# Integrated single cell spatial multi-omics landscape of WHO grades 2-4 diffuse gliomas identifies locoregional metabolomic regulators of glioma growth

**DOI:** 10.1101/2025.04.30.651361

**Authors:** Yanxia Ma, Shamini Ayyadhury, Sanjay Singh, Yashu Vashishath, Cagri Ozdemir, Trevor D. McKee, Nhat Nguyen, Akshay Basi, Duncan Mak, Javier A. Gomez, Jason T. Huse, Sana Noor, Dan Winkowski, Regan Baird, Jeffrey S. Weinberg, Frederick F. Lang, Jared K. Burks, Serdar Bozdag, Erin H. Seeley, Chibawanye I. Ene

## Abstract

Diffuse infiltrating gliomas are aggressive tumors of the central nervous system driven by intra-tumoral heterogeneity and aberrant normal-tumor cell-cell interactions. Grade specific and locoregional metabolic dependencies driving aberrant cell-states linked to treatment resistance, seizures and infiltration of gliomas remain elusive. Here, we applied spatial transcriptomics (stRNAseq), imaging mass cytometry (IMC) and mass spectrometry imaging (MSI; metabolites, peptides and glycans) to the core and edge tumor tissue from patients with World Health Organization (WHO) grades 2-4 diffuse infiltrating gliomas including isocitrate dehydrogenase (IDH) mutant oligodendrogliomas (WHO Grades 2 and 3) and IDH wildtype astrocytomas including ‘anaplastic’ astrocytoma (prior 2016 WHO histological grade 3) and glioblastoma (GBM, WHO grade 4) stRNAseq identified regions-specific differentially expressed genes with significant overall survival implications particularly in IDH wildtype GBM. Integration of stRNA seq and MSI-derived metabolite expression demonstrated enrichment of L-glutamine in *SOX4*+ Neural progenitor-like (NPC-like) cells and DL-dopamine in *GPNMB*^+^ Mesenchymal-like (MES-like) GBM cells at the tumor edge relative to the core. Our results uncover clinically relevant and locoregional cell state-specific metabolites that may contribute to GBM proliferation, infiltration and seizures. This comprehensive pan-diffuse infiltrating glioma multi-omics study could serve as a resource for uncovering region-specific metabolic vulnerabilities encompassing metabolites, glycans and peptides within transcriptionally defined cell states across WHO 2-4 diffuse glioma.

## Introduction

Gliomas are a diverse group of tumors that originating from cells within the brain.^1^ The World Health Organization (WHO) classifies gliomas into 4 grades based on histological and molecular features.^2^ Diffuse infiltrating gliomas (WHO grade 2-4) are an incurable form of glioma due to their infiltration nature and resistance to treatment.^3^ Despite aggressive surgery followed by cytotoxic chemotherapies and radiation therapy, diffuse infiltrating gliomas ultimately recur leading to significant morbidity and mortality.^3–5^ The complexity and heterogeneity of the glioma tumor micro-environment (TME) contributes to treatment resistance.^6^ There are also locoregional differences in metabolic drivers of tumor and non-tumor cells within the glioma TME that contribute to poor clinical outcomes.^7–11^ Up to now, however, spatial profiling of diffuse infiltrating gliomas has been limited to glioblastoma (GBM, WHO grade 4).^12,13^ So far, there are no spatially resolved single cell multi-omics studies from metabolically distinct regions of isocitrate dehydrogenase (IDH) mutant and IDH wildtype diffuse gliomas encompassing oligodendrogliomas and astrocytoma histologies. In this study, we integrated spatial transcriptomics (stRNAseq), imaging mass cytometry (IMC) and mass spectrometry imaging (MSI) data from core and edge tissues of IDH mutant oligodendrogliomas (WHO grades 2 and 3) and IDH wildtype astrocytomas including ‘anaplastic’ astrocytoma (prior 2016 WHO histological grade 3) and glioblastoma (WHO grade 4). Our results from isocitrate dehydrogenase wildtype (IDHwt) GBM, the most aggressive form of diffuse infiltrating glioma, revealed regional transcriptional defined cell state-enriched metabolites that may contribute to tumor, progression, infiltration, seizures. Our study provides an atlas of spatially resolved transcriptomics, proteomics, metabolomics (metabolites, glycans and peptides) of core and edge tissue from WHO grades 2-4 diffuse infiltrating gliomas.

## Materials and Methods

### Tissue preparation

We utilized a glioma tissue micro-array (TMA) to establish the neoplastic and non-neoplastic heterogeneity in the GBM infiltration edge. TMA tissue samples are meticulously constructed through a multi-step process. In briefly, a cylindrical core of tissue is extracted from a donor paraffin block using a specialized hollow needle. This core, typically ranging from 1mm to 4mm in diameter, was carefully selected to represent the area of interest within the original tissue sample. Following core removal, it was precisely positioned and embedded into a recipient paraffin block at a predefined location. This recipient block can accommodate numerous cores from various donor blocks, allowing for the simultaneous analysis of multiple tissue samples under uniform experimental conditions. Following TMA assembly, sections were cut on a rotary microtome at 5um thickness, then mounted onto the appropriate slide required by the experiment. Formalin fixed paraffin embedded (FFPE) tissue was obtained from residual clinical material with the patients’ consent under protocols LAB03-0320 or 2012-0441 and analyzed under protocol 2022-0979. A waiver of consent was granted to 2022-0979 by MD Anderson’s institutional review board (IRB) due to the research involving no more than minimal risk to subjects. In this study, each TMA slide consists of pathologist-annotated tumor core and edge samples of Glioblastoma (IDH wildtype GBM, WHO Grade 4; n=10), ‘Anaplastic’ Astrocytoma (IDH wildtype AA, prior 2016 WHO Grade 3; n=3), Oligodendroglioma (IDH mutant Oligo, WHO Grade 2 and 3; n=5) and 2 non-brain control samples.

### 10x Genomics Xenium Single Cell Spatial Transcriptomics

To analyze gene expression while preserving tissue architecture, we performed 10x Genomics Xenium in Situ gene expression assay per 10x protocol. For a slide of glioma tissue micro-array (TMA) with the neoplastic and non-neoplastic heterogeneity in the GBM infiltration edge, a targeted gene panel, Xenium human brain cancer In Situ Gene Expression Panel (10x Genomics, In Situ Gene Expression v1.0, FFPE Human Brain Cancer Data with Human Immuno-Oncology Profiling Panel and Custom Add-on is shown in Supplemental Table 1), was designed to detect hundreds of mRNA transcripts in TMA tissue sections. After deparaffinization with 2 times of Xylene bath for 10 mins each, and a series of Ethanol bath (100%, 96%, 96% and 70%) for 3 minutes each, the tissue slide was processed to decrosslinking at 80°C for 30 mins followed by 2 times of PBS-T incubation for 1 minute each at room temperature (CG000580). The slide with the tissue section was then processed as described in the Xenium In Situ Gene Expression (CG000582) workflow. In brief, the targeted barcoded probes were hybridized and specifically bound to the target RNA sequences within the tissue, followed by rolling circle amplification (RCA) to generate rolling circle products (RCPs), which can then be sequenced. The RCP readout and data were processed by 10x Genomics Xenium Analyzer instrument and on-board analysis (Xenium Instrument Bundle, PN-1000569 - Xenium Analyzer, Analysis Computer, Instrument Accessory Kits). The single cell spatial RNA sequencing data can be visualized using Xenium Explorer visualization software (CG000584).

### Imaging mass cytometry (IMC)

To identify major cell types based on surface markers, we performed imaging mass cytometry on the TMA section adjacent to the 10x Genomics Xenium slide. First, we optimized an antibody panel (Supplemental Fig. 2) for our TMA samples. Antibodies were selected as follows (anti-GFAP, Standard BioTool, Cat#3143030D ; anti-NeuN, Standard BioTool, Cat#3145019D; anti-MAP2,Standard BioTool, Cat#3148023D; anti-O4, R&D system, Cat#MAB1326; anti-lba1, Standard BioTool, Cat#3142020D; anti-CX3CR1, abcam, Cat#ab308614; anti-CCK, LSBio, Cat#LS-B15905-50; anti-CD34, antibodies, Cat#A85659; anti-P2Y12, abcam, Cat#ab240285; anti-FAM20C, abcam, Cat#ab154740; anti-CD133, CST, Cat #51917; anti-CD15/SSEA1 (MC480), CST, Cat#74180). Antibodies conjugated with metal labels are shown in Supplemental Table 2. Cell segmentation was determined using Maxpar® IMC™ Cell Segmentation kit (Standard BioTools, Cat# 201500). Other antibodies (anti-CD3, anti-CD45, anti-E-Cad, anti-CD68, anti-CD276/B7-H3, anti-Arginase 1, anti-EGFR, anti-CD163, anti-Ki67, anti-Vimentin, anti-CD4, anti-CD8a) were obtained from Flow Cytometry & Cellular Imaging Core Facility, Department of Leukemia, The University of Texas M. D. Anderson Cancer Center. TMA section was processed in antigen retrieval EZ-AR2 Elegance RTU (pH9, Cat#HK547-XAK) buffer at 107°C for 15 mins. The slide was then blocked with 3% BSA in TBS buffer and 1% horse serum (ThermoFisher, Cat#16050130) for 2 hours at room temperature. Metal conjugated antibodies were added to TMA slide for incubation at 4°C overnight. After washing with TBS-T buffer and TBS buffer at room temperature, TMA slide was incubated with Ir-intercalator solution for 5 minutes at room temperature. Slide was then washed with TBS buffer and rinsed with ddH2O. It was then dry thoroughly for analyzed with the Hyperion™ XTi Imaging Mass Cytometer (Flow Cytometry & Cellular Imaging Core Facility, Department of Leukemia, The University of Texas M. D. Anderson Cancer Center). The high-dimensional spatial data at subcellular resolution from the Hyperion™ XTi Imaging System were generated and visualized using MCD SmartViewer v1.0 (FLDM-400316).

### Mass spectrometry imaging (MSI)

As described in a previous study^14^, the next adjacent TMA section from the 10x Xenium tissue were dewaxed using xylene for 2 x 3 minutes with fresh xylene for each step. The slides were allowed to dry before fiducial points were placed at the corners of the slides by etching with a diamond scribe. Digital images were acquired at 4800 dpi using an Epson Perfection V600 flatbed scanner. The slides were coated with 1,5-diaminonaphthalene using an HTX M5 Robotic Reagent Sprayer (HTX Technologies, Chapel Hill, NC). Mass spectrometry images of metabolites were acquired in negative ion mode at 20 μm resolution using a Bruker timsTOF fleX Mass Spectrometer (Bruker Daltonics, Billerica, MA). After metabolite imaging, the matrix was removed using 100% ethanol. The sections were rehydrated in graded ethanol and antigen retrieved with 100 mM Tris, pH 9.0 using a Biocare Medical Decloaker (Biocare Medical, Pacheco, CA) at 95°C for 20 minutes. After cooling for 15 minutes, the slides were buffer exchanged to water over 5 changes and allowed to dry completely. The sections were coated with PNGaseF (Bulldog Bio, Portsmouth, NH) and incubated at 37°C for 2 hours in humidity chambers1 before being coated with α-cyano-4-hydroxycinnamic acid (CHCA) for N-linked glycan imaging using the timsTOF fleX in positive ion mode. After glycan imaging, the matrix was removed, and the slides were rehydrated and again antigen retrieved as described above. The slides were coated with trypsin (MS Grade, Fisher Scientific, Pittsburgh, PA) and incubated at 37°C for 4 hours in humidity chambers before again being coated with CHCA. Peptide images were acquired in positive ion mode on the timsTOF fleX. Finally, the matrix was removed using ethanol and the slides were stained with hematoxylin and eosin using standard protocols. Digital microscopy images were acquired of the stained slides using a Hamamatsu NanoZoomerSQ Digital Slide Scanner (Hamamatsu Photonics, Bridgewater, NJ) at 20X magnification. A complete summary of parameters used for matrix/enzyme sprayer and mass spectrometry data acquisition are provided in Supplemental Tables 3 and 4, respectively. Files were loaded into SCiLS Lab 2024a (Bruker) and root mean square normalized for visualization of the ion images. Each dataset was manually peak picked, avoiding known matrix and non-monoisotopic peaks. The attribute editor was used to add core location, tumor type, and tumor grade information for each core that was used for calculation of p-values and ROC curves. Metabolites and glycans were putatively identified using the SCiLS MetaboScape 2023b (Bruker) plugin. Metabolites were searched against the human metabolite database with [M-H]-, [M]-, and [M+Cl]-primary ions considered while glycans were searched as [M+H]+, [M+Na]+, and [M+K]+ primary ions.

### Pre-processing Xenium Cell-by-Gene Matrix of GBM TMA

The cell feature matrix file was imported using sc.read_10x_h5 (scanpy 1.11.0) creating an ‘AnnDat’ object. A total of 174, 261 xenium segmented cells were imported representing 20 TMA cores. The median transcript counts, and median gene counts for the entire GBM cohort was 141 and 61 respectively. Individually, each TMA core had median transcript and median gene numbers ranging from 11-312 and 63-158. Cells were filtered by individual cores based on the following criteria: dropping all cells that had more than 9 transcripts (equivalent to the median control probe transcript detected per cell) and cells that had *n* genes <6 (equivalent to the median unique control probes identified per cell). Peripheral cells away from the tumor cores were removed.

### Scaling, Normalization and Transformation

The TMA cores were grouped by their respective tumor types by grade: IDH mutant oligodendroglioma (Grade 2 and 3), IDH wildtype ‘anaplastic’ astrocytoma (prior 2016 WHO grade 3), IDH wildtype glioblastoma (Grade 4), and merged within their tumor types. A scale factor of 500 was applied to all tumor types for normalization of transcript counts across cells before applying a log1p transformation across each tumor type.

### Cell Type Annotation

Next, cell type label transfer was carried out using a reference dataset. We used GBmap, a single cell reference dataset that has been extensively characterized and annotated at multiple hierarchical levels.^15^ The extended GBmap cohort comprising of 1,135,677 cells was downloaded. Only the droplet-based single cell transcriptome datasets that were profiled using the 10xGenomics assays were included (10x 3’ v2,v3 and 10x 5’ v1 chemistries). This truncated GBmap dataset was further trimmed to include only genes present in the Xenium gene panel set resulting in the final dataset which we called GBmapX. We selected ‘annotation level 3’ from GBmap to compute our cell type label transfer.

To assign cell type labels to Xenium single cells, we implemented a correlation-based label transfer method using the GBmapX reference dataset. We first build a reference expression profile, denoted μ, for each cell type c and gene g, by averaging the expression values of gene g across all cells of type c in the GBmapX reference dataset. For each Xenium cell j, with expression vector X_j_, we computed the Pearson correlation coefficient r between X_j_ and each reference profile μ. Each Xenium cell j is assigned the label corresponding to the cell type ĉ_j_ with the highest Pearson correlation coefficient. A label is only assigned if the corresponding correlation is statistically significant (p-value ≤ 0.001).

### Cell Type Proportion

Cell type proportions were calculated by determining the fraction of each cell type within the tumor type cohorts. This was achieved by aggregating the number of cells of each type across all cores and dividing by the total number of cells, then multiplying by 100 to express the proportion as a percentage as depicted by the following function:

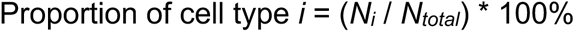

Where *N_i_* is the number of cells of type *i*, and *N_total_* is the total number of cells.

Similar calculations were performed to determine cell type proportions for each TMA core from the tumor core and edge separately, as well as for each individual TMA core.

### Selection of Highly Variable Genes using Moran’s I

To emphasize spatial variability in clustering, Moran’s *I* statistic, embedded within the Squidpy package (V1.5.0) was used to select for highly variable genes. Moran’s I is a spatial autocorrelation method where positive values identify spatially autocorrelated or aggregated genes.^16^ Spatially variable genes were calculated for each TMA core individually. Each TMA core, representing a patient sample derived from the ‘Edge’ or ‘Core’ was scaled as described above and log transformed. Subsequently spatial nearest neighbors were computed by calling the squidpy.gr.spatial_neighbors() tool before applying the sq.gr.spatial_autocorr tool (using mode=’moran’) to identify the top variable genes for each sample. We selected the set of genes with *I* > 0.2 (*Gj : G = number of genes with i>0.2, j = Sample)*. To derive the final spatially variable gene set, we calculated the **union** of these gene sets across all samples. For *n* samples, the final gene set, *G_final*, is given by:

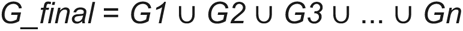

G_final for each tumor type was calculated and used for downstream dimensional reduction and clustering as outlined below.

### Dimensional Reduction, Batch Correction, and Leiden Clustering

We performed dimensional reduction and leiden clustering on the log1p normalized tumor types and found significant batch effects were present, despite all samples stained on the same slide. To mitigate batch effects, we performed batch correction using Harmony across samples in a tumor type (oligodendroglioma, ‘anaplastic’ astrocytoma, glioblastoma,). We used G_final to subset each tumor type dataset. Then we applied principal component analysis (PCA) to reduce the dimensionality of the data, retaining 21 principal components, determined by measuring the minimum number of components required to include 90% of total variance. Nearest neighbors were selected using *n* = 15 neighbors via the ‘sc.pp.neighbors’ function from scanpy. The leiden method, a community detection algorithm, was applied to the compressed data to identify spatial clusters. Leiden clusters were computed using resolution parameters ranging from 0.1 to 1.0 at 0.1 resolution intervals for a total of ten resolutions. The *Calinksi Harabasz Score, Davies Bouldin Score* and the Adjusted *Rand Index* were used to determine the optimal number of clusters, which were determined as 0.4, 0.4, 0.3 for the glioblastoma, ‘anaplastic’ astrocytoma and oligodendroglioma batch corrected tumor type datasets.

### Differential Gene Expression between ‘Core’ and ‘Edge’

Differential gene expression between the ‘Core’ and ‘Edge’ samples in all tumor type datasets was computed using the ‘scanpy.tl.rank_genes_groups’ function with default settings. The Wilcoxon rank-sum test was used to calculate p-values. Significantly expressed genes from the ‘Core’ versus ‘Edge’ were utilized for survival analysis.

### Survival Analysis

Gene expression and clinical data for 525 glioblastoma multiforme (GBM) patients were obtained from The Cancer Genome Atlas (TCGA), specifically from the Broad Institute’s Affymetrix HT Human Genome U133A microarray platform (Level 3, RMA-normalized expression data). Survival analysis was conducted to evaluate associations between individual gene expression levels and overall survival. For each gene, patients were stratified into three expression-based groups: *upregulated*, *downregulated*, and *neutral*. Grouping was based on standardized thresholds defined as ± 0.5 standard deviations (SD) from the gene-wise mean expression value across all samples. Patients with expression values greater than (mean + 0.5 × SD) were classified as *upregulated*, and those with values below (mean − 0.5 × SD) as *downregulated*. The remaining samples were classified as neutral. To evaluate the association of each gene with survival, we first excluded samples categorized as “neutral.” The remaining patients (upregulated vs. downregulated) were included in a multivariable Cox proportional hazards model using the coxph() function from the survival R package. The model was adjusted for available covariates including age, sex, and race. Model fitting was performed using the Breslow method for handling tied survival times and robust variance estimation. For genes yielding significant associations (Wald test *p* < 0.05), Kaplan–Meier survival curves were generated using the survfit() function and visualized with the ggsurvplot() function from the survminer package. All statistical analyses and visualizations were performed in R version 4.4.2.

### Cell-Cell Interaction

To investigate spatial cluster enrichment and relationships between cell types within these clusters, SCIMAP(v2.2.11), a scalable toolkit for analyzing spatial molecular data, was used to perform spatial interaction mapping.^17^ The ‘scimap.tl.spatial.interaction’ method was called to compute the strength of spatial interactions between the cell types. It computes this by calculating the likelihood of cell types being adjacent to each other against a permutated random background. Only tumor cell types were used to compute interactions (immune and accessory cell types were removed from each tumor type dataset). Default parameters were used except for the following changes – permutations=500, knn=3. The discriminator in interaction mapping was the region of the dataset – ‘Core’ versus ‘Edge’. Only interactions with p-value < 0.001 and interaction strength > 0.25 were included in further analysis and for the construction of a circular directed network map.

### Xenium-derived DAPI, Mass Spectrometry Image and Image Mass Cytometry Co-Registration

Virtual Alignment of pathoLogy Image Series or VALIS, a fully automated pipeline for image registration was used to align the xenium-derived DAPI and mass spectrometry derived whole slide images (WSI) using rigid transformations. Briefly, WSIs are converted to numpy arrays using low resolution images and processed to single channel images and masks are automatically defined for tissue focused registration. Image features are extracted and matched using the metabolites image as a reference. The morphology focus DAPI images were used for image alignment. A single channel, with broad intensity distribution across all the TMAs, was selected as a representative from the peptide, glycan and metabolite dataset for image alignment. The Xenium DAPI image, together with the representative peptide and glycan intensity image from a single channel, were then aligned towards the reference image (metabolite intensity channel image), with rigid registration performed serially. This was followed by non-rigid registration. Error estimation was calculated using the distance between registered matched features from the full resolution images. The transformations from the registered VALIS object were used to warp the x,y coordinates (converted to pixels) of the xenium derived cell centroids and single cell polygon vertices to align with the reference image coordinate framework. Intensity values from the glycans, peptides and metabolites images were extracted and tabularized along with the warped x,y coordinates of each pixel. Intensities from 287, 818, 670 channels for glycans, peptides and metabolites were extracted.

### Metabolite, Glycan and Peptide Enrichment

We performed a second round of differential gene expression analysis, this time between cell types within each tumor type dataset. Following this, for the genes indicated as most significant in the survival analysis, we identified their highest enrichment in tumor transcriptional cell-states (MES-like, AC-like, OPC-like and NPC-like). We next generated polygons from cells labelled with the cell types enriched for these genes and further filtered cells that showed non-zero expression of these genes to generate binary masks. Using our co-registered images for the glycans, metabolites and peptides, we extracted pixels whose centroid coordinates fell within the enclosed masked regions. To show metabolite co-localization with proliferating tumor cells (Ki67+) in figure 4e, analysis was performed in *Visiopharm’s* Discovery platform. Images were co-registered using automated TissueAlign module. Nuclei in the IMC image were segmented using pre-trained AI-driven algorithm available in the Discovery platform. The average intensity of the full IMC panel and MSI images were calculated.

### Code and data availability

Data will be made available upon publication of manuscript. The analysis code was developed by Astraea Bio and is available upon reasonable request for academic use.

## Results

### Tissue micro-array generation from WHO diffuse glioma grades 2-4 core and edge

We generated a tissue micro-array (TMA; Supplemental Table 5) comprising a neuropathologist annotated tumor core and edge tissue from WHO grade 2 and 3 (IDH mutant Oligodendrogliomas or IDH mut Oligo; n=5), prior 2016 WHO grade 3 (IDH wildtype ‘anaplastic’ astrocytoma or IDH wt AA, n=4) and WHO grade 4 (IDH wild-type glioblastoma or IDH wt GBM, n=10) in addition to control spleens (n=2; Clinical information Supplemental Table 6). FFPE embedded tissue on the TMA was analyzed using the Xenium spatial transcriptomics platform. Following spatial transcriptomics analysis, the same slide was stained with H&E. IMC and MSI were performed on adjacent sections respectively. (See Fig. 1 for workflow of tissue collection and analysis).

**Figure 1.**
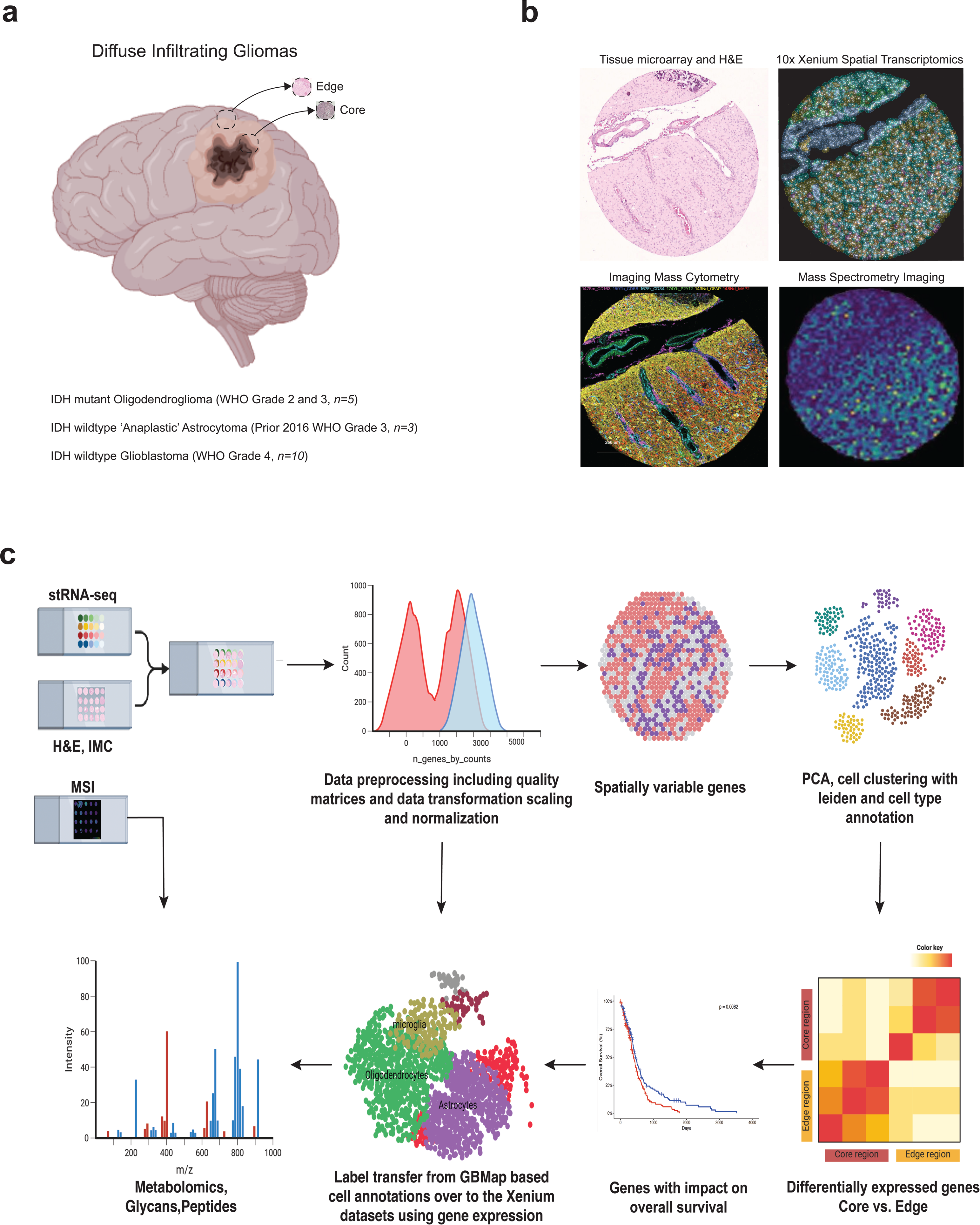
Overview of workflow and multi-omic integration. **a**. Surgical resections from the tumor core and edge were carried out for three tumor types. Parallel sections of these tumor resections were interrogated by three different modalities to spatially profile transcriptomics, protein expression and biomolecular content (peptides, metabolites and glycans). **b.** Representative TMA sections from a single patient, geometrically aligned, showing the molecular profile from each modality. **c.** Computational workflow highlights the main pre-processing and post-processing steps leading up to integration between MSI intensity values and gene expression.

### Spatial transcriptomics reveals cellular heterogeneity across the tumor core and edge from diffuse infiltrating glioma WHO grade 2-4

Using the Xenium platform, we evaluated cellular heterogeneity across the landscape of diffuse infiltrating gliomas. GBmap, a single cell brain cancer atlas was used to resolve cellular identity and characterize the cellular heterogeneity across diffuse gliomas on the TMA. We specifically used the annotation-level-3 cell type labels that allowed the identification of the following cell states and types - ‘RG’, ‘Neuron’, ‘Oligodendrocyte’, ‘OPC’, ‘’Astrocyte’, ‘OPC-like’, ‘AC-like’, ‘MES-like’, ‘NPC-like’, ‘Mono’, ‘TAM-BDM’, ‘DC’, ‘CD4/CD8’, ‘Neutrophil’, ‘B cell’, ‘Plasma B’, ‘NK’, ‘TAM-MG’, ‘Mast’, ‘Mural’ and ‘Endothelial’ cell types. Though GBmap is a reference database derived from IDHwt GBM database, we reasoned that the generic cell-type labelling would be reflected accurately across the other diffuse gliomas namely IDHmut Oligos (WHO grade 2 and 3) and IDHwt AA (prior 2016 WHO grade 3).

We reduced GBmap to include only the genes that were present in the xenium custom gene panel (333 genes; Supplemental Table 1) and derived the gene expression vector for each cell type (*cell type annotation in methods*). Using this reference cell-type gene expression vector, we computed the Pearson correlation for each cell from the spatial transcriptomics datasets and assigned cell type labels. In addition, we performed k-nearest neighbor and subsequent Uniform Manifold Approximation Projection (UMAP) to compress the datasets into a 2-dimensional projection. Based on this GBmap reference label transfer, we noted no distinct enrichment of cell types between ‘Core’ versus ‘Edge’ and that all cell types were present in both ‘Core’ and ‘Edge’ samples (Fig. 2a-b, d-e, g-h). Next, we evaluated each grade of diffuse gliomas. Analysis of the landscape of the IDHmut Oligos (WHO Grades 2 and 3) dataset showed that the tumor type was comprised primarily of OPC-like and NPC-like cell states (Fig. 2a-b). Strikingly, the cell type composition for IDHmut Oligos was restricted to a few cell types and more homogenous compared to the other diffuse glioma grades. Relative to the tumor core, the IDHmut Oligo tumor edge showed an increase in non-neoplastic tumor associated oligodendrocytes (TAO) and astrocytes (TAA) (Fig. 2b). The cellular landscape of the IDHwt AA (prior 2016 WHO grade 3) dataset was more heterogenous compared to IDHmut Oligo but concordant between the ‘Edge’ and ‘Core’ (Fig. 2d-e). IDHwt AA tumors were comprised predominantly of MES-like tumor cell states and TAAs (Fig. 2d-e). There was a slight increase in the MES-like tumor cell state and TAAs at the IDHwt AA tumor edge dataset relative to the core (Fig. 2e). The AC-like tumor cell state within IDHwt AA, however, were significantly more enriched in the tumor core relative to the edge (Fig. 2e). There was significantly lower proportions of non-neoplastic and peripherally derived tumor-associated bone marrow derived macrophages (TAM-BDM) at the IDHwt AA tumor edge with no significant change to the brain resident tumor-associated microglia (TAM-MG) proportion (Fig. 2e). IDHwt GBM were also noted to be significantly more heterogenous relative to IDHmut Oligo (Fig. 2g-h). The IDHwt GBM tumor core was comprised primarily of tumor cells in the AC-like and MES-like states (Fig. 2g-h). The ‘Astrocyte’ or TAA, ‘OPC-like’ and ‘NPC-like’ populations were significantly enriched in the ‘Edge’ dataset and rare in the ‘Core’ dataset from IDHwt GBM (Fig. 2h). We also found a significant increase in TAA and TAM-MG populations at the IDHwt GBM edge (Fig. 2h). Across all diffuse glioma grades, the most predominant immune subset were myeloid cells including TAM-MG, TAM-BDM and monocytes, while non-myeloid subsets including CD4+, CD8+ T-cells, NK cells and neutrophils were rare (Fig. 2b,e,h).

**Figure 2.**
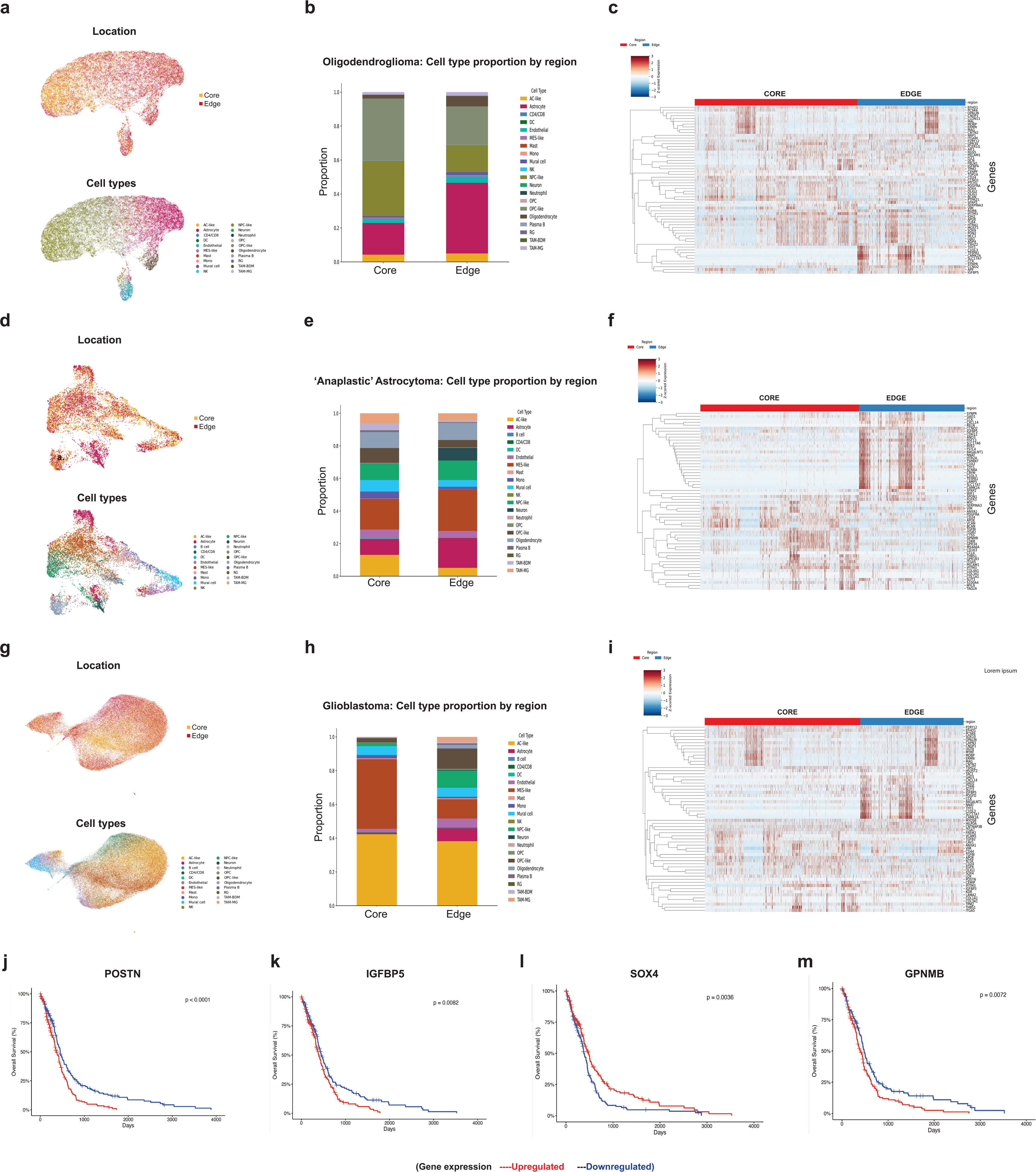
Locoregional cellular and transcriptional heterogeneity across diffuse infiltrating gliomas (WHO grades 2-4). **a-i**. Top left UMAP colored by ‘Core’ versus ‘Edge’ and bottom left UMAP is colored by the cell type labels. Right bar plot showing cell type proportions present in ‘Core’ versus ‘Edge’. Heat map showing differentially expressed genes between ‘Core’ versus ‘Edge’. IDH mutant Oligodendroglioma **(a-c)**, IDH wildtype ‘anaplastic’ Astrocytoma **(prior 2016 WHO glioma classification; d-f)**, and IDH wildtype GBM **(g-i)**. **j-m**. Kaplan meier survival curves showing impact of upregulated or downregulated differentially expressed genes between ‘Core’ v. ‘Edge’ of IDH wildtype GBM in The Cancer Genome Atlas (TCGA), p<0.05 is significant.

### Clinical significance of locoregional differentially expressed genes across diffuse infiltrating gliomas

We performed differential expression analysis using spatially variable genes to discover differentially expressed genes from the core and edge of within each diffuse glioma grade (Fig. 2c,f,i). We identified several differentially expressed genes at the core and edge specific to each grade (Fig. 2c,f,i and Supplemental Table 7). To determine if any differentially expressed genes between the core and edge of IDHwt GBM influenced overall survival (OS), we analyzed the association between OS and gene expression levels (low vs. high) in IDHwt GBM samples from the cancer genome atlas (TCGA). At least 4 genes (*POSTN, TGFBI, LOX and IGFBP5*) that are significantly up regulated in the IDHwt GBM core were associated with significantly worse OS (Fig. 2j-k; Supplemental Table 8). Other genes (*ADRA1A, CORO1A, ELOVL2, GPNMB, and SOX4)* that were significantly up regulated at the IDHwt GBM edge also significantly impacted OS (Fig. 2l-m; Supplemental Table 8).

### Cell-Cell interactions

To explore cellular neighborhoods and interactions associated with poor clinical outcomes, we performed SCIMAP, an analytical package that includes functions for various spatial analysis, including computing cell type proximity associations. We focused on identifying enrichment of tumor cell-states and non-neoplastic central nervous system (CNS) cell types because of their spatial proximity and removed immune cells from our analysis. In the IDH mutant Oligo tumor core, there were strong interactions between AC-like tumor cell states and TAA as well as between non-neoplastic TAOs and TAAs (Fig. 3a and Supplemental Fig. 1). At the IDHmut Oligo tumor edge, the strongest interaction occurred between NPC-like tumor cell states and TAOs (Fig. 3b and Supplemental Fig. 1). In IDHwt AA (prior 2016 WHO grade 3) tumor core, there were strong interactions between MES-like and AC-like tumor cell states as well as between MES-like and OPC-like tumor cell states (Fig. 3c, and Supplemental Fig. 1). At the IDHwt AA (prior 2016 WHO grade 3) tumor edge, the strongest interactions occurred between AC-like tumor cell states and TAAs (Fig. 3d and Supplemental Fig. 1). In IDHwt GBM core, the strongest interactions occurred between AC-like tumor cells and TAA as well between MES-like tumor cells and TAOs (Fig. 3e and Supplemental Fig. 1). At the IDHwt GBM tumor edge, the strongest interactions occurred between OPC-like tumor cell states and TAA as well as between TAA and endothelial cells (Fig. 3f and Supplemental Fig. 1).

**Figure 3.**
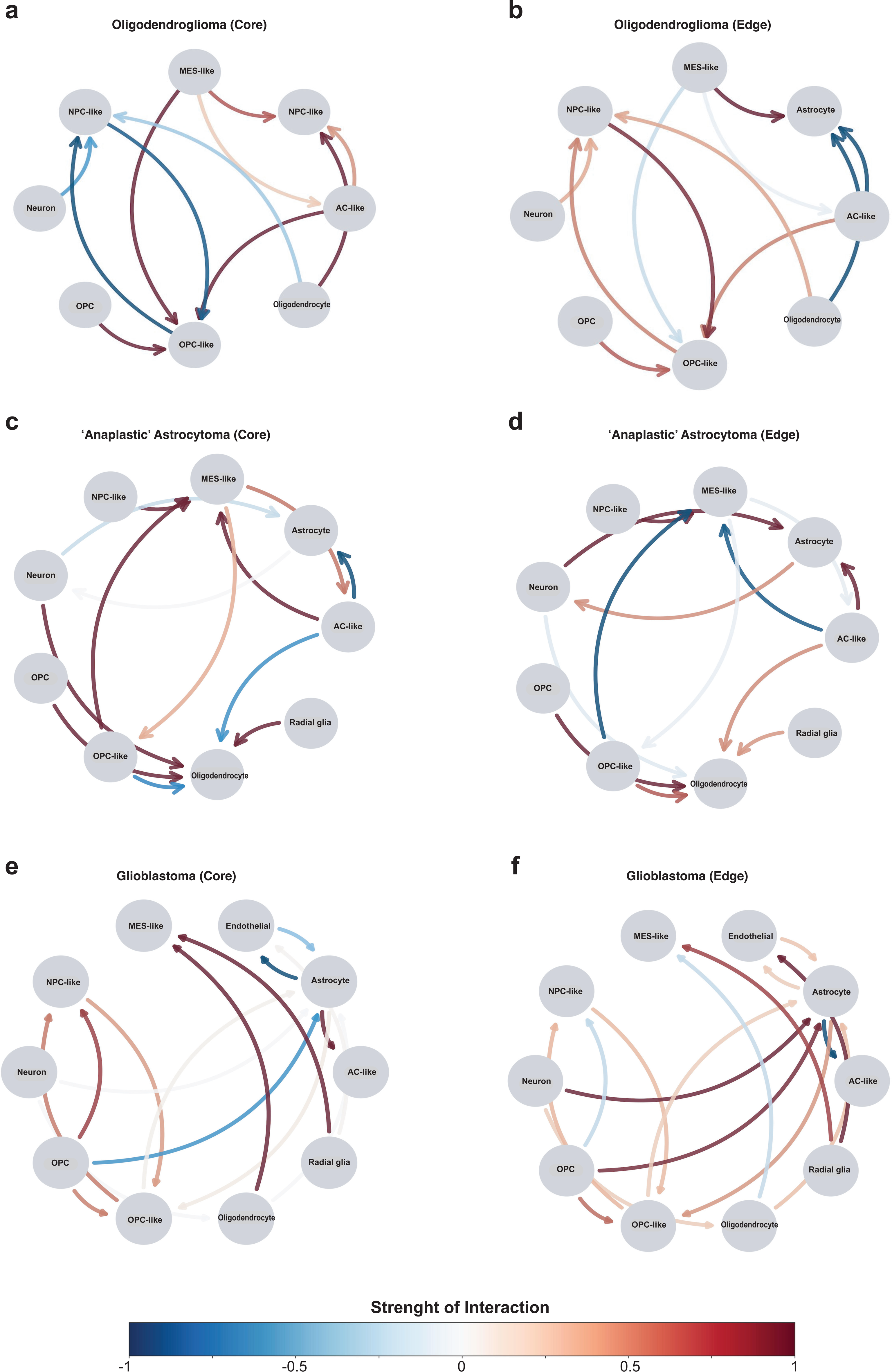
Region-specific tumor cellular interactions across diffuse gliomas WHO grades 2-4. **a-f.** Circular interaction plots for ‘Core’ versus ‘Edge’ from IDH mutant Oligodendroglioma **(a-b)**, IDH wildtype ‘Anaplastic’ Astrocytomas **(prior 2016 WHO glioma classification; c-d)** and IDH wildtype GBM **(e-f).** Each plot showing the directed pairs of interactions for tumor and non-tumor cell types, filtered for two levels of significance (p-value < 0.0001 and interaction strength between -1 to 1).

### Spatially resolved cell-state specific metabolite enrichment in IDH wildtype GBM

Next to identify metabolites enriched in specific transcriptional cell states with high expression of clinically relevant differentially expressed genes between the core and edge of IDHwt GBM (Fig. 4a), we co-registered and aligned the MSI data with stRNA-seq resolved cell states expressing the clinically relevant differentially expressed genes within IDHwt GBM. We find that all differentially expressed genes with survival implications were significantly higher expression within tumor cells compared to non-tumor cells (Fig. 4a). MES-like tumor cell states had higher expression of *GPNMB, TGFBI, IGFBP5* and *LOX* mRNA. AC-like tumor cell states had higher expression of *POSTN* and *ELOVL2* mRNA. NPC-like tumor cell states had higher expression of *SOX4* mRNA. We also find that the most significant overlaps between metabolites and tumor cell states within IDHwt GBM enriched with clinically relevant differentially expressed genes occurred at the tumor edge (Fig. 4b-d). The exceptions were *IGFBP5^+^* MES-like tumor cells which showed significant metabolite overlaps in both the core and the edge of IDHwt GBM (Fig. 4b) and *ELAVL2^+^*AC-like tumor cells where the overlap with metabolites was most significant at the tumor core (Fig. 4c). Known metabolites that were significantly enriched in specific cell states expressing clinically relevant differentially expressed genes at the IDHwt GBM edge included 4-hydroxycyclohexanecarboxylic acid (4-Hchc) within *IGFBP5*^+^ MES-like tumor cells (Fig. 4b), DL-dopamine within *GPNMB^+^ MES*-like tumor cells (Fig. 4b) as well as L-glutamine within *SOX4*^+^ NPC-like tumor cells (Fig. 4d).

**Figure 4.**
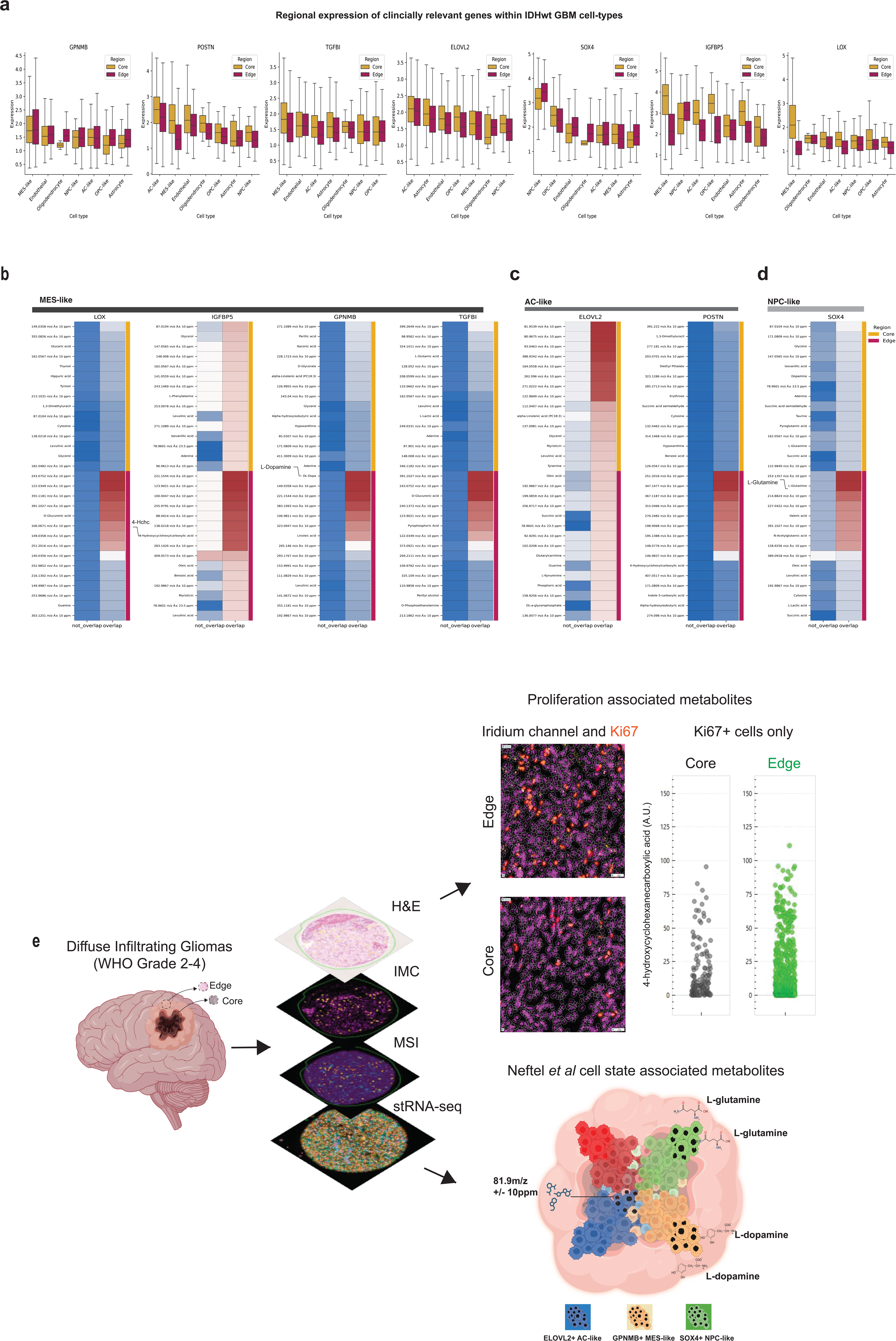
Identification of region-specific IDH wildtype GBM cell state enriched metabolites based on integrated mass spectrometry imaging and spatial transcriptomics. **a**. Enrichment of differentially expressed genes with survival implications within IDHwt GBM cell states and cell types. **b-d**. Using a common co-ordinate and co-registration approach from adjacent slides we identified overlaps between MSI derived metabolites and MES-like (**b**), AC-like (**c**) and NPC-like (**d**) tumor cell states in IDHwt GBM. **e.** Illustrative display showing co-registered images from multiple imaging modalities on diffuse glioma core or edge tissue. *Top-*For metabolite overlay on Ki67^+^ proliferating tumor cells, H&E, IMC, MSI are co-registered. Top arrow leads to zoomed view of IMC image showing iridium (DNA) channels and Ki67. Overlay results showing segmented cells. Ki67+ cells are displayed in Red (Edge tissue sample on top, Core tissue sample on bottom). Comparison plot of Ki67+ cells organized by 4-Hydroxy intensity (Grey: Core tissue sample; Green: Edge Tissue sample). *Bottom-*Showing Neftel *et al* ^21^ transcriptional cell states in IDHwt GBM enriched with clinically relevant genes ELAVL2 in AC-like tumor cells, GPNMB in MES-like tumor cells and SOX4 in NPC-like tumor cells. Following co-registration and overlay of stRNA-seq and MSI, annotated and unannotated metabolites corresponding to transcriptional cell states expressing clinically relevant genes in specific regions of the tumor are identified. 81.9m/z +/- 10ppm in ELAVL2+ AC-like cells in tumor core (blue), L-dopamine in GPNMB+ MES-like cells in tumor edge (orange), L-glutamine in SOX4 expressing NPC-like cells in tumor edge (green).

## Discussion

Diffuse infiltrating gliomas remains one of the most complex and heterogenous TMEs across cancers.^18^ Spatial transcriptomics (stRNA-seq) has revolutionized understanding of the single cell pathology landscape of WHO grade 4 diffuse infiltrating gliomas (IDHmutant astrocytoma and IDH wildtype glioblastoma).^12,19^ Consistent with our findings, these studies demonstrated invasive niche specific enrichment of specific cell states such as OPC-like and NPC-like tumor cell states in the invasive front of GBM.^20,21^ Few studies, however, have performed a comprehensive multi-omic (transcriptional and metabolomic) profiling of all diffuse infiltrating gliomas encompassing WHO grades 2-4. In this study, we integrated multi-omics data from all diffuse glioma grades and our results could provide a roadmap for uncovering putative locoregional and grade specific metabolic vulnerabilities at single cell resolution.

The clinically relevant differentially expressed genes between the core and edge of IDHwt GBM had significantly higher mRNA expression within tumor cells relative to non-tumor cells in the TME (Fig. 4a). This indicates that tumor cell expression of the niche-specific differentially expressed genes in our study likely drove the observed clinical outcomes. The AC-like and MES-like tumor cell states within gliomas are associated with poor clinical outcomes.^22^ The distribution and cell-cell interactions of these specific cell states within the glioma TME ultimately determine the overall clinical impact.^23^ Metabolic and hypoxic gradients organize the structural continuum of these tumor cell states within the glioma TME.^24^ In this study, we combined stRNA-seq, IMC and MSI to profile metabolically distinct regions (core v. edge) of all diffuse infiltrating gliomas including IDHmut oligo (WHO grade 2 and 3), IDHwt AA (prior 2016 WHO grade 3) and IDHwt GBM (WHO grade 4). We identified a strong cell-cell interaction between AC-like tumor cell states and TAA in the cores of IDH mutant Oligo and IDHwt GBM as well as the edge of IDHwt AA. Based on prior *in-vitro* studies, a bidirectional relationship between glioma cells and TAA is associated with chemotherapy resistance, tumor cell proliferation and inflitration.^25,26,27,28^ Therefore, our results indicate that the AC-like tumor cell state interaction with TAA occurs across all diffuse gliomas. The clinical consequence of this interaction in diffuse gliomas may be grade and location specific. OPC-like tumor cells were exclusively enriched in the IDHwt GBM edge (relative to the core) where they demonstrated a strong interaction with TAA relative to the tumor core. Based on the pro-infiltration effect of TAA on gliomas at the tumor edge, these results indicate that OPC-like tumor cell state within IDHwt GBM may contribute to tumor infiltration via a cross talk with TAAs in the tumor periphery.^20^

The MES-like tumor cell states were enriched in the cores of IDHwt AA and IDHwt GBM. In the IDHwt AA core, MES-like tumor cells showed the strongest interaction with all the other cell states including NPC-like, OPC-like and AC like tumor cells. In IDHwt GBM core, MES-like tumor cells had the strongest interaction TAOs and radial glia (RG). These findings are consistent with prior reports on the role of MES-like cells in the GBM, where there are physical interactions between MES-like cells and non-neoplastic cells using gap junctions and glioma microtubes^24,29^ This makes the MES-like cell state the most suitable to form tumor networks that can be targeted. To this end, it was shown that GBM networks can withstand chemical damage from Temozolomide (TMZ) and can regenerate its connections among remaining cells after surgery by increasing microtube formation.^30^ The hubs of this network are called ‘periodic cells’ and they represent around 4% of all GBM tumor cells. Importantly ‘periodic cells’ possess a MES-like cell state and initiate intracellular calcium currents that boost tumor proliferation and regulate microtube dynamics for networking and invasion. Selective ablation of the hub cells impaired the overall tumor growth; however, inherent tumor plasticity drove a change from ‘non-periodic’ to ‘period cells’ in the tumor TME. Altogether, these findings indicate that targeting cell-states involved in pro-tumor cell-cell interactions or networks that drive plasticity, tumor growth and invasion may be a clinically relevant and feasible therapeutic strategy.

Cell state plasticity is responsible for the intra-tumoral and inter-tumoral heterogeneity that makes gliomas elusive to treatment.^31–33^ Using stRNA-seq and MSI, Ravi *et al* identified spatially distinct and clinically relevant transcriptional programs within IDHwt GBM that are influenced by immunological and metabolic stress factors within the tumor micro-environment.^13^ These results indicated that the various cell states within IDHwt GBM originate from plasticity and dynamic adaptation to various environmental cues including metabolites. Therefore, they concluded that there is a bi-directional tumor -host interdependence that drives GBM infiltration and clinical outcomes. To uncover cell state specific metabolites, peptides and glycans that may contribute to plasticity and poor clinical outcome in diffuse infiltrating gliomas, we integrated spatially resolved stRNA-seq and MSI data from 2 distinct regions (core v. edge) of all grades of diffuse infiltrating gliomas. *SOX4* and *GPNMB* mRNA were significantly enriched in NPC-like cell and MES-like cell states respectively at the IDHwt GBM edge compared to the tumor core. SOX4 is a known stemness and neural development marker that is expressed in NPC-like GBM cells where it contributes to the maintenance of a stem-like phenotype.^34^ Unlike the other differentially expressed genes between the core and edge of IDHwt GBM, high *SOX4* mRNA expression was associated with improved OS. This indicates that the biological impact of *SOX4* expression within NPC-like tumor cells at the IDHwt GBM edge may not be increased proliferation or infiltration. Based on metabolomics data, we find enrichment of L-glutamine within *SOX4*^+^ NPC-like tumor cells in the edge of IDHwt GBM. NPC-like tumor GBM cells are known to form glutaminergic-neuronal synapses which induce epileptic activity within the brain.^35^ Therefore, these results indicate that *SOX4*^+^ NPC-like tumor cells at the tumor edge may contribute to seizure activity in IDHwt GBM.

Gpnmb (glycoprotein non-metastatic melanoma protein B or Osteoactivin) plays a crucial role in inducing a MES-like cell state within GBM.^36^ We find that *GPNMB*+ MES-like cells in the IDHwt GBM edge are enriched with L-dopamine. Dopamine receptor activation promotes GBM stemness and growth via MET activation.^37^ *In-vitro*, it has been shown that dopamine induces transferrin receptor 1 expression to assist with iron update and iron addition within MES-like GBM cells.^38^ This provided a clinical rationale for eradicating treatment resistant MES-like GBM stem cells by targeting ferroptosis. Our integrated stRNA-seq and MSI data corroborate these *in-vitro* studies demonstrating an oncogenic role of L-dopamine in *GPNBM+* MES-like cells in the IDHwt edge. Ultimately, with a larger dataset from all diffuse glioma grade that encompasses previously un-identified and un-annotated metabolites, glycans and peptides, our study provides an unprecedented opportunity to uncover targetable locoregional metabolic vulnerabilities associated with proliferative (Ki67^+^) and poor-prognosis transcriptional cell states in diffuse gliomas (Fig. 4e).

In conclusion, we performed a comprehensive spatial multi-omics profiling (stRNA-seq, IMC and MSI) on tissue samples derived from both the core and edge of diffuse gliomas across all grades. We present a common coordinate and co-registration methodology for integrating and analyzing these multi-omics datasets. Our results from IDHwt GBM implicate annotated and un-annotated metabolites enriched within transcriptionally defined cell states in distinct regions of IDHwt GBM. These results provide insights into cell-state specific locoregional metabolic vulnerabilities that may be targeted in diffuse gliomas clinically to minimize tumor proliferation, infiltration, seizures and plasticity-driven treatment resistance.

### Limitations

The TMA used in this study included 4 IDHwt ‘anaplastic’ astrocytomas, which represent a histopathologic diagnosis based on the prior 2016 WHO glioma classification.^39^ The 2021 WHO glioma molecular classification phased out the ‘anaplastic’ designation in these IDHwt astrocytomas and these tumors are now classified as IDHwt GBM if there is evidence EGFR amplification, TERT promoter mutation and chromosome 7 gain and chromosome 10 loss.^2^ In support of this, our single cell spatial profiling demonstrated significant similarity in cellular heterogeneity across IDHwt ‘anaplastic’ astrocytoma and IDHwt GBM samples. There are ongoing efforts to obtain molecular information on the historical IDHwt ‘anaplastic’ astrocytoma samples to determine if these tumors are IDHwt GBM based on the 2021 WHO glioma molecular classification.

## Supporting information

All Supplemental Tables

## Acknowledgements

We thank Dr. Gregory Fuller (Neuropathologist) for providing GBM TMAs for optimizing IMC conditions. We would also like to thank the MD Anderson Brain Tumor Center histopathology core including Lisa Norberg, Truc Kuo and Jennifer Ritchie for their assistance with creating the TMA used in the study. This work was supported by The University of Texas MD Anderson Cancer Center SPORE in Brain Cancer Career Enhancement Program Award (P50CA127001 to C.E), University of Texas MD Anderson Cancer Center Physician Scientist Program Award (600378-125266 to C.E), Cancer Prevention and Research Institute of Texas (CPRIT; RP190617 and RP240559 to E.S) and Sachs Family Funds (J.W), National Institute of General Medical Sciences of the National Institutes of Health (R35GM133657 to S.B).

## Author contributions

C.E. and S.S. conceived the study. J.H. annotated core v. edge from surgical samples. Y.M., S.S. and N.N. performed stRNA-seq and IMC. S.A., T.M., S.N., D.W., R.G. integrated and analyzed stRNA-seq, IMC and MSI datasets. Y.M., S.S., A.B., D.M., J.G. and J.B. optimized IMC antibodies and conditions. Y.V., C.O. and S.B. performed clinical outcomes analysis from differentially expressed genes. E.S. performed and analyzed MSI datasets. J.W. and F.L. contributed surgical samples. Y.M., S.A. and C.E. wrote the paper.

## Supplemental Figure

**Figure 1.** Heatmap showing Cell-Cell interaction strengths across regions all grades of diffuse gliomas.

## Supplemental Table Legends

**Table 1.** 10x Custom Spatial Transcriptomics Gene Expression Panel

A targeted gene panel, Xenium human brain cancer In Situ Gene Expression Panel (10x Genomics, In Situ Gene Expression v1.0, FFPE Human Brain Cancer Data with Human Immuno-Oncology Profiling Panel and Custom Add-on), is shown in this table. This Custom Add-on gene panel was designed to detect hundreds of mRNA transcripts in a slide of glioma tissue micro-array (TMA) with the neoplastic and non-neoplastic heterogeneity in the GBM infiltration edge.

**Table 2.** Imaging mass cytometry panel

An optimized antibody panel was selected to simultaneously stain and quantify multiple markers in an adjacent TMA section, which is close to the tissue section used in 10x Genomics Xenium analysis. The cell population was identified based on expression of the indicated protein markers.

**Table 3.** Mass spectrometry imaging acquisition parameters

Bruker timsTOF fleX Imaging Acquisition Parameters used in our MSI analysis are listed in the table.

**Table 4.** Mass spectrometry sprayer parameters

HTX M5 Robotic Reagent Sprayer Parameters used in our MSI analysis are shown in the table.

**Table 5.** Brain tumor tissue microarray layout

The layout of glioma tissue micro-array comprising a neuropathologist annotated tumor core and edge tissue from WHO grade 2 (IDH mutant Oligodendrogliomas; n=5), WHO grade 3 (IDH wildtype ‘anaplastic’ astrocytoma, n=4) and WHO grade 4 (IDH wild-type glioblastoma, n=10) in addition to control spleens (n=2)

**Table 6.** Clinical information of samples on brain tumor tissue microarray

Basic clinical information of samples on brain tumor tissue microarray are listed.

**Table 7.** Spatial transcriptomics derived differentially expressed genes across all diffuse gliomas

Expression levels of genes that show a significant difference in Core v. Edge across all samples.

**Table 8.** Differentially expressed genes with significant overall survival impact from the cancer genome atlas for GBM

**Table 9.** MSI derived metabolomics overlap with 10x spatial transcriptomics derived cell states and differentially expressed genes in core v. edge

