## Supplemental Figure 1 for "Integrated single cell spatial multi-omics landscape of WHO grades 2-4 diffuse gliomas identifies locoregional metabolomic regulators of glioma growth"

Supplemental Figure 1. Heatmap showing Cell-Cell interaction strengths across regions all grades of diffuse gliomas

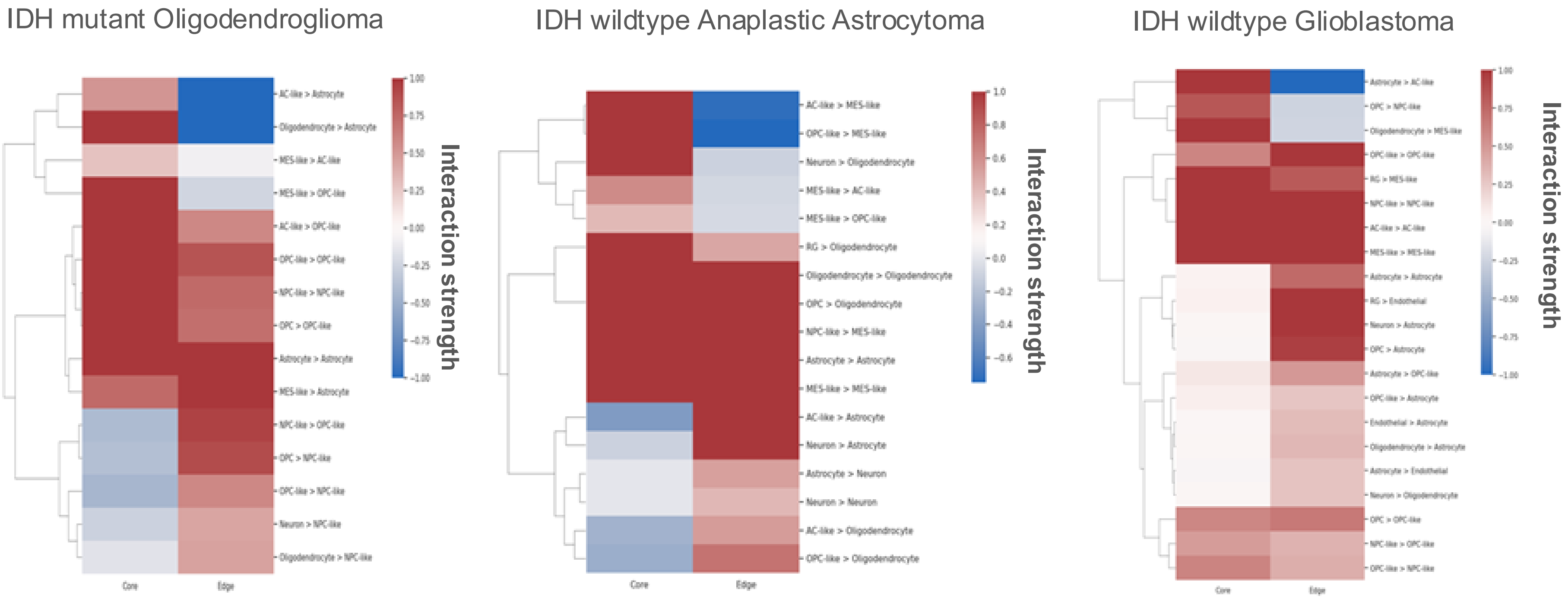
