## Supplementary material for "Integrated single cell spatial multi-omics landscape of WHO grades 2-4 diffuse gliomas identifies locoregional metabolomic regulators of glioma growth": All Supplemental Tables

**Supplemental Table 1. 10X Xenium Custom Panel**

| Gene | Annotation | Ensembl ID |
|---|---|---|
| BANK1 | B cells | ENSG00000153064 |
| CD19 | B cells | ENSG00000177455 |
| CD27 | B cells | ENSG00000139193 |
| CD79A | B cells | ENSG00000105369 |
| CD79B | B cells | ENSG00000007312 |
| MS4A1 | B cells | ENSG00000156738 |
| TCL1A | B cells | ENSG00000100721 |
| CDH4 | Bulk CN Signature | ENSG00000179242 |
| ELOVL2 | Bulk CN Signature | ENSG00000197977 |
| FGFR3 | Bulk CN Signature | ENSG00000068078 |
| MEOX2 | Bulk CN Signature | ENSG00000106511 |
| PAX6 | Bulk CN Signature | ENSG00000007372 |
| PTPRA | Bulk CN Signature | ENSG00000132670 |
| SEPTIN11 | Bulk CN Signature | ENSG00000138758 |
| SOX9 | Bulk CN Signature | ENSG00000125398 |
| BCL3 | Bulk MES Signature | ENSG00000069399 |
| COL1A2 | Bulk MES Signature | ENSG00000164692 |
| IL6 | Bulk MES Signature | ENSG00000136244 |
| ITGB1 | Bulk MES Signature | ENSG00000150093 |
| LOX | Bulk MES Signature | ENSG00000113083 |
| MMP7 | Bulk MES Signature | ENSG00000137673 |
| VDR | Bulk MES Signature | ENSG00000111424 |
| CDKN1C | Bulk PN Signature | ENSG00000129757 |
| EPHB1 | Bulk PN Signature | ENSG00000154928 |
| ERBB3 | Bulk PN Signature | ENSG00000065361 |
| GABRB3 | Bulk PN Signature | ENSG00000166206 |
| HDAC2 | Bulk PN Signature | ENSG00000196591 |
| HOXD3 | Bulk PN Signature | ENSG00000128652 |
| CD1A | Dendritic cells | ENSG00000158477 |
| CD1C | Dendritic cells | ENSG00000158481 |
| PECAM1 | Endothelial cells | ENSG00000261371 |
| EPCAM | Epithelial cells | ENSG00000119888 |
| KRT14 | Epithelial cells | ENSG00000186847 |
| BRINP3 | Excitatory neurons | ENSG00000162670 |
| C1QL3 | Excitatory neurons | ENSG00000165985 |
| CA10 | Excitatory neurons | ENSG00000154975 |
| CCK | Excitatory neurons | ENSG00000187094 |
| CDH12 | Excitatory neurons | ENSG00000154162 |
| CDH13 | Excitatory neurons | ENSG00000140945 |
| CNTN5 | Excitatory neurons | ENSG00000149972 |
| DCC | Excitatory neurons | ENSG00000187323 |
| DCN | Excitatory neurons | ENSG00000011465 |

| | | |
|---|---|---|
| DNER | Excitatory neurons | ENSG00000187957 |
| EPHA4 | Excitatory neurons | ENSG00000116106 |
| FBXO36 | Excitatory neurons | ENSG00000153832 |
| GPC6 | Excitatory neurons | ENSG00000183098 |
| GRIK1 | Excitatory neurons | ENSG00000171189 |
| GRM8 | Excitatory neurons | ENSG00000179603 |
| HAT1 | Excitatory neurons | ENSG00000128708 |
| HPCA | Excitatory neurons | ENSG00000121905 |
| HS3ST4 | Excitatory neurons | ENSG00000182601 |
| IGFBP4 | Excitatory neurons | ENSG00000141753 |
| IL1RAPL2 | Excitatory neurons | ENSG00000189108 |
| KIRREL3 | Excitatory neurons | ENSG00000149571 |
| LMO4 | Excitatory neurons | ENSG00000143013 |
| MAML2 | Excitatory neurons | ENSG00000184384 |
| MEF2C | Excitatory neurons | ENSG00000081189 |
| MEIS2 | Excitatory neurons | ENSG00000134138 |
| MGAT4C | Excitatory neurons | ENSG00000182050 |
| NFIA | Excitatory neurons | ENSG00000162599 |
| NKAIN3 | Excitatory neurons | ENSG00000185942 |
| NPTXR | Excitatory neurons | ENSG00000221890 |
| NR3C2 | Excitatory neurons | ENSG00000151623 |
| NRGN | Excitatory neurons | ENSG00000154146 |
| NRP1 | Excitatory neurons | ENSG00000099250 |
| NTNG1 | Excitatory neurons | ENSG00000162631 |
| NWD2 | Excitatory neurons | ENSG00000174145 |
| PDZRN3 | Excitatory neurons | ENSG00000121440 |
| PLCXD3 | Excitatory neurons | ENSG00000182836 |
| PROX1 | Excitatory neurons | ENSG00000117707 |
| PTCHD4 | Excitatory neurons | ENSG00000244694 |
| PTPRK | Excitatory neurons | ENSG00000152894 |
| PTPRT | Excitatory neurons | ENSG00000196090 |
| RAB3C | Excitatory neurons | ENSG00000152932 |
| RELN | Excitatory neurons | ENSG00000189056 |
| RFX3 | Excitatory neurons | ENSG00000080298 |
| RGS4 | Excitatory neurons | ENSG00000117152 |
| RGS6 | Excitatory neurons | ENSG00000182732 |
| ROBO1 | Excitatory neurons | ENSG00000169855 |
| RORB | Excitatory neurons | ENSG00000198963 |
| SGCD | Excitatory neurons | ENSG00000170624 |
| SGCZ | Excitatory neurons | ENSG00000185053 |
| SHISA6 | Excitatory neurons | ENSG00000188803 |
| TENM3 | Excitatory neurons | ENSG00000218336 |
| TRPM3 | Excitatory neurons | ENSG00000083067 |
| TRPS1 | Excitatory neurons | ENSG00000104447 |
| TSHZ2 | Excitatory neurons | ENSG00000182463 |
| UNC5D | Excitatory neurons | ENSG00000156687 |
| WIPF3 | Excitatory neurons | ENSG00000122574 |
| ZBTB20 | Excitatory neurons | ENSG00000181722 |
| ZMAT4 | Excitatory neurons | ENSG00000165061 |
| ZNF804B | Excitatory neurons | ENSG00000182348 |
| POSTN | Fibroblasts | ENSG00000133110 |

| | | |
|---|---|---|
| SFRP2 | Fibroblasts | ENSG00000145423 |
| AKAP12 | Glioblastoma | ENSG00000131016 |
| ANXA1 | Glioblastoma | ENSG00000135046 |
| AQP4 | Glioblastoma | ENSG00000171885 |
| B4GALNT1 | Glioblastoma | ENSG00000135454 |
| BCAN | Glioblastoma | ENSG00000132692 |
| C1orf194 | Glioblastoma | ENSG00000179902 |
| C9orf24 | Glioblastoma | ENSG00000164972 |
| CAV1 | Glioblastoma | ENSG00000105974 |
| CDK4 | Glioblastoma | ENSG00000135446 |
| CDKN2A | Glioblastoma | ENSG00000147889 |
| CHI3L1 | Glioblastoma | ENSG00000133048 |
| CNN3 | Glioblastoma | ENSG00000117519 |
| DLL3 | Glioblastoma | ENSG00000090932 |
| EGFR | Glioblastoma | ENSG00000146648 |
| ETV1 | Glioblastoma | ENSG00000006468 |
| FABP7 | Glioblastoma | ENSG00000164434 |
| FBLN1 | Glioblastoma | ENSG00000077942 |
| FOXJ1 | Glioblastoma | ENSG00000129654 |
| GAP43 | Glioblastoma | ENSG00000172020 |
| GATM | Glioblastoma | ENSG00000171766 |
| GJA1 | Glioblastoma | ENSG00000152661 |
| HEPN1 | Glioblastoma | ENSG00000221932 |
| HES6 | Glioblastoma | ENSG00000144485 |
| HILPDA | Glioblastoma | ENSG00000135245 |
| HOPX | Glioblastoma | ENSG00000171476 |
| IDH1 | Glioblastoma | ENSG00000138413 |
| IDH2 | Glioblastoma | ENSG00000182054 |
| IDO1 | Glioblastoma | ENSG00000131203 |
| IDO2 | Glioblastoma | ENSG00000188676 |
| IGFBP3 | Glioblastoma | ENSG00000146674 |
| IGFBP5 | Glioblastoma | ENSG00000115461 |
| IGFBP7 | Glioblastoma | ENSG00000163453 |
| IL13RA2 | Glioblastoma | ENSG00000123496 |
| ITGB8 | Glioblastoma | ENSG00000105855 |
| KISS1R | Glioblastoma | ENSG00000116014 |
| LANCL2 | Glioblastoma | ENSG00000132434 |
| LIFR | Glioblastoma | ENSG00000113594 |
| LPL | Glioblastoma | ENSG00000175445 |
| MAP1B | Glioblastoma | ENSG00000131711 |
| MAP2 | Glioblastoma | ENSG00000078018 |
| MDM2 | Glioblastoma | ENSG00000135679 |
| MEST | Glioblastoma | ENSG00000106484 |
| METRN | Glioblastoma | ENSG00000103260 |
| MGST1 | Glioblastoma | ENSG00000008394 |
| MRPS17 | Glioblastoma | ENSG00000239789 |
| NF1 | Glioblastoma | ENSG00000196712 |
| NMB | Glioblastoma | ENSG00000197696 |
| NNAT | Glioblastoma | ENSG00000053438 |
| NPDC1 | Glioblastoma | ENSG00000107281 |
| NRN1 | Glioblastoma | ENSG00000124785 |
| NTRK2 | Glioblastoma | ENSG00000148053 |
| OTOS | Glioblastoma | ENSG00000178602 |
| PMP2 | Glioblastoma | ENSG00000147588 |
| PRDX2 | Glioblastoma | ENSG00000167815 |

| | | |
|---|---|---|
| PTEN | Glioblastoma | ENSG00000171862 |
| PTPRZ1 | Glioblastoma | ENSG00000106278 |
| RBP1 | Glioblastoma | ENSG00000114115 |
| S100A16 | Glioblastoma | ENSG00000188643 |
| SAA1 | Glioblastoma | ENSG00000173432 |
| SCRG1 | Glioblastoma | ENSG00000164106 |
| SLPI | Glioblastoma | ENSG00000124107 |
| SOX11 | Glioblastoma | ENSG00000176887 |
| SOX2 | Glioblastoma | ENSG00000181449 |
| SOX4 | Glioblastoma | ENSG00000124766 |
| SPARC | Glioblastoma | ENSG00000113140 |
| SRI | Glioblastoma | ENSG00000075142 |
| STAT3 | Glioblastoma | ENSG00000168610 |
| STMN2 | Glioblastoma | ENSG00000104435 |
| TGFB1 | Glioblastoma | ENSG00000105329 |
| TGFB2 | Glioblastoma | ENSG00000092969 |
| TOP2A | Glioblastoma | ENSG00000131747 |
| TRIL | Glioblastoma | ENSG00000255690 |
| TSC22D1 | Glioblastoma | ENSG00000102804 |
| TSPAN3 | Glioblastoma | ENSG00000140391 |
| TTYH1 | Glioblastoma | ENSG00000167614 |
| TUBB2B | Glioblastoma | ENSG00000137285 |
| VGF | Glioblastoma | ENSG00000128564 |
| GAPDH | Housekeeping | ENSG00000111640 |
| AIF1 | Immune | ENSG00000204472 |
| ALOX5AP | Immune | ENSG00000132965 |
| AREG | Immune | ENSG00000109321 |
| ARHGDIB | Immune | ENSG00000111348 |
| BIN1 | Immune | ENSG00000136717 |
| C15orf48 | Immune | ENSG00000166920 |
| C1orf162 | Immune | ENSG00000143110 |
| CAPG | Immune | ENSG00000042493 |
| CCL20 | Immune | ENSG00000115009 |
| CCL5 | Immune | ENSG00000271503 |
| CD163 | Immune | ENSG00000177575 |
| CD2 | Immune | ENSG00000116824 |
| CD274 | Immune | ENSG00000120217 |
| CD3D | Immune | ENSG00000167286 |
| CD48 | Immune | ENSG00000117091 |
| CD52 | Immune | ENSG00000169442 |
| CD69 | Immune | ENSG00000110848 |
| CD7 | Immune | ENSG00000173762 |
| CD83 | Immune | ENSG00000112149 |
| CD86 | Immune | ENSG00000114013 |
| CH25H | Immune | ENSG00000138135 |
| CHCHD7 | Immune | ENSG00000170791 |
| CORO1A | Immune | ENSG00000102879 |
| COTL1 | Immune | ENSG00000103187 |
| CSTA | Immune | ENSG00000121552 |
| CTLA4 | Immune | ENSG00000163599 |
| CTSS | Immune | ENSG00000163131 |
| CX3CR1 | Immune | ENSG00000168329 |
| CXCR4 | Immune | ENSG00000121966 |
| CYTIP | Immune | ENSG00000115165 |
| DUSP2 | Immune | ENSG00000158050 |

| | | |
|---|---|---|
| FCGBP | Immune | ENSG00000275395 |
| FCGR1A | Immune | ENSG00000150337 |
| FCGR3A | Immune | ENSG00000203747 |
| G0S2 | Immune | ENSG00000123689 |
| GNLY | Immune | ENSG00000115523 |
| GPNMB | Immune | ENSG00000136235 |
| GPR34 | Immune | ENSG00000171659 |
| GZMA | Immune | ENSG00000145649 |
| HAMP | Immune | ENSG00000105697 |
| HLA-DMB | Immune | ENSG00000242574 |
| HLA-DQA1 | Immune | ENSG00000196735 |
| HLA-DQB1 | Immune | ENSG00000179344 |
| HLA-DRB5 | Immune | ENSG00000198502 |
| HMOX1 | Immune | ENSG00000100292 |
| IER3 | Immune | ENSG00000137331 |
| IFI27 | Immune | ENSG00000165949 |
| IL32 | Immune | ENSG00000008517 |
| ITGB2 | Immune | ENSG00000160255 |
| KLF2 | Immune | ENSG00000127528 |
| KLF4 | Immune | ENSG00000136826 |
| KLRB1 | Immune | ENSG00000111796 |
| KLRC1 | Immune | ENSG00000134545 |
| LST1 | Immune | ENSG00000204482 |
| LTB | Immune | ENSG00000227507 |
| LY86 | Immune | ENSG00000112799 |
| MS4A6A | Immune | ENSG00000110077 |
| MT1G | Immune | ENSG00000125144 |
| MT1H | Immune | ENSG00000205358 |
| NKG7 | Immune | ENSG00000105374 |
| NR4A2 | Immune | ENSG00000153234 |
| NUPR1 | Immune | ENSG00000176046 |
| PDCD1 | Immune | ENSG00000188389 |
| PLIN2 | Immune | ENSG00000147872 |
| RASGEF1B | Immune | ENSG00000138670 |
| RB1 | Immune | ENSG00000139687 |
| REL | Immune | ENSG00000162924 |
| RGS10 | Immune | ENSG00000148908 |
| RNASET2 | Immune | ENSG00000026297 |
| S100A12 | Immune | ENSG00000163221 |
| S100A4 | Immune | ENSG00000196154 |
| SLC11A1 | Immune | ENSG00000018280 |
| SPI1 | Immune | ENSG00000066336 |
| STXBP2 | Immune | ENSG00000076944 |
| TGFBI | Immune | ENSG00000120708 |
| TMIGD3 | Immune | ENSG00000121933 |
| TNFAIP3 | Immune | ENSG00000118503 |
| TRAC | Immune | ENSG00000277734 |
| TRBC1 | Immune | ENSG00000211751 |
| TRBC2 | Immune | ENSG00000211772 |
| TREM2 | Immune | ENSG00000095970 |
| TYMP | Immune | ENSG00000025708 |
| VAMP8 | Immune | ENSG00000118640 |
| VCAN | Immune | ENSG00000038427 |
| VSIG4 | Immune | ENSG00000155659 |
| ADGRE1 | Microglia | ENSG00000174837 |

| | | |
|---|---|---|
| APOE | Microglia | ENSG00000130203 |
| CD40 | Microglia | ENSG00000101017 |
| CD68 | Microglia | ENSG00000129226 |
| CD74 | Microglia | ENSG00000019582 |
| CST3 | Microglia | ENSG00000101439 |
| CTSH | Microglia | ENSG00000103811 |
| CXCR1 | Microglia | ENSG00000163464 |
| IL1B | Microglia | ENSG00000125538 |
| ITGAM | Microglia | ENSG00000169896 |
| ITGAX | Microglia | ENSG00000140678 |
| LYVE1 | Microglia | ENSG00000133800 |
| MAF | Microglia | ENSG00000178573 |
| P2RY12 | Microglia | ENSG00000169313 |
| P2RY13 | Microglia | ENSG00000181631 |
| PTPRC | Microglia | ENSG00000081237 |
| CD14 | Monocytes | ENSG00000170458 |
| FCER1A | Myeloid dendritic cells | ENSG00000179639 |
| RBFOX3 | Neurons | ENSG00000167281 |
| CD247 | NK cells | ENSG00000198821 |
| FCER1G | NK cells | ENSG00000158869 |
| CSPG4 | Oligodendrocyte precursor cel | ENSG00000173546 |
| PDGFRA | Oligodendrocyte precursor cel | ENSG00000134853 |
| MBP | Oligodendrocytes | ENSG00000197971 |
| OLIG1 | Oligodendrocytes | ENSG00000184221 |
| OLIG2 | Oligodendrocytes | ENSG00000205927 |
| OLIG3 | Oligodendrocytes | ENSG00000177468 |
| SOX10 | Oligodendrocytes | ENSG00000100146 |
| LILRA4 | Plasmacytoid dendritic cells | ENSG00000239961 |
| CCNA1 | Proliferation | ENSG00000133101 |
| CCNB2 | Proliferation | ENSG00000157456 |
| CDK1 | Proliferation | ENSG00000170312 |
| CENPF | Proliferation | ENSG00000117724 |
| KIT | Proliferation | ENSG00000157404 |
| MKI67 | Proliferation | ENSG00000148773 |
| PCNA | Proliferation | ENSG00000132646 |
| RACGAP1 | Proliferation | ENSG00000161800 |
| CCR7 | T cells | ENSG00000126353 |
| CD28 | T cells | ENSG00000178562 |
| CD3G | T cells | ENSG00000160654 |
| CD4 | T cells | ENSG00000010610 |
| CD8A | T cells | ENSG00000153563 |
| CD8B | T cells | ENSG00000172116 |
| FASLG | T cells | ENSG00000117560 |
| FOXP3 | T cells | ENSG00000049768 |
| GPR183 | T cells | ENSG00000169508 |
| GZMB | T cells | ENSG00000100453 |
| GZMK | T cells | ENSG00000113088 |
| IL2RA | T cells | ENSG00000134460 |
| IL7R | T cells | ENSG00000168685 |
| AATK | TME (non-immune) | ENSG00000181409 |
| ABCA2 | TME (non-immune) | ENSG00000107331 |
| APOD | TME (non-immune) | ENSG00000189058 |
| CAPN3 | TME (non-immune) | ENSG00000092529 |
| CLDN11 | TME (non-immune) | ENSG00000013297 |
| CNDP1 | TME (non-immune) | ENSG00000150656 |

| | | |
|---|---|---|
| CNTN2 | TME (non-immune) | ENSG00000184144 |
| EDIL3 | TME (non-immune) | ENSG00000164176 |
| EFHD1 | TME (non-immune) | ENSG00000115468 |
| ERMN | TME (non-immune) | ENSG00000136541 |
| FGFR2 | TME (non-immune) | ENSG00000066468 |
| HHATL | TME (non-immune) | ENSG00000010282 |
| KCNMB4 | TME (non-immune) | ENSG00000135643 |
| KLK6 | TME (non-immune) | ENSG00000167755 |
| LHPP | TME (non-immune) | ENSG00000107902 |
| MAG | TME (non-immune) | ENSG00000105695 |
| MAL | TME (non-immune) | ENSG00000172005 |
| MOBP | TME (non-immune) | ENSG00000168314 |
| MOG | TME (non-immune) | ENSG00000204655 |
| MYRF | TME (non-immune) | ENSG00000124920 |
| OPALIN | TME (non-immune) | ENSG00000197430 |
| RAPGEF5 | TME (non-immune) | ENSG00000136237 |
| S100A1 | TME (non-immune) | ENSG00000160678 |
| SLC24A2 | TME (non-immune) | ENSG00000155886 |
| SPOCK3 | TME (non-immune) | ENSG00000196104 |
| UGT8 | TME (non-immune) | ENSG00000174607 |

**Table 2. Antibody Panel for IMC Staining**

| Ab | Label | Target | Catalog |
|---|---|---|---|
| 1 | 143Nd | Gfap (Astrocytes, glial tumor cells) | 3143030D |
| 2 | 145Nd | NeuN (Neurons) | 145019D |
| 3 | 148Nd | MAP2 (Neurons) | 3148023D |
| 4 | 150Nd | O4† (Oligodendocytes) | AB1326 |
| 5 | 169Tm | Iba1 | 3142020D |
| 6 | 161Dy | CX3CR1† (Chemokine receptor for migratory cells) | ab308614 |
| 7 | 162Dy | CCK† (Infiltration marker) | LS-B15905-50 |
| 8 | 167Er | CD34 (Blood vessels) | A85659 |
| 9 | 174Yb | P2Y12† (Activated microglia) | |
| | | **Maxpar IMC Cell Segmentation Kit** | |
| 10 | 195Pt | ICSK1 | 201500 |
| 11 | 196Pt | ICSK2 | 201500 |
| 12 | 198Pt | ICSK3 | 201500 |
| | | **Target** | |
| 13 | 170Er | CD3 (T-cells) | CORE |
| 14 | 152Sm | CD45 (Immune cell) | CORE |
| 15 | 158Gd | E-Cad (Infiltration) | CORE |
| 16 | 159Tb | CD68 (Peripheral macrophage) | CORE |
| 17 | 173Yb | CD276/B7-H3 | CORE |
| 18 | 164Dy | Arginase 1 (Pro-tumor macrophage) | CORE |
| 19 | 142Nd | EGFR (Tumor) | CORE |
| 20 | 147Sm | CD163 (Peripheral macrophages) | CORE |
| 21 | 168Er | Ki67 (Proliferating tumor cells) | CORE |
| 22 | 141Pr | Vimentin (Tumor) | CORE |

| | | | |
|---|---|---|---|
| 23 | CD4 156Gd (IMC) | CD4 | CORE |
| 24 | 160Gd | CD8a | CORE |

**Supplemental Table 3. Bruker timsTOF fleX Imaging Acquisition Parameters**

| | Metabolites | Glycans | Peptides |
|---|---|---|---|
| Polarity | Negative | Positive | Positive |
| *m/z* range | 50-600 | 700-3500 | 600-4500 |
| Number of Laser Shots | 300 | 300 | 400 |
| Funnel 1 RF (Vpp) | 75.0 | 450.0 | 450.0 |
| Funnel 2 RF (Vpp) | 100.0 | 500.0 | 500.0 |
| Multipole RF (Vpp) | 150.0 | 500.0 | 600.0 |
| Collision Energy (eV) | 5.0 | 10.0 | 10.0 |
| Collision RF (Vpp) | 500.0 | 2700.0 | 3400.0 |
| Quadrupole Ion Energy (eV) | 5 | 5 | 5 |
| Transfer Time (μs) | 35.0 | 140.0 | 180.0 |
| Pre Pulse Storage (μs) | 3.0 | 14.0 | 18.0 |

**Supplemental Table 4. HTX M5 Robotic Reagent Sprayer Parameters**

| | Metabolite Matrix | Glycan Matrix | Peptide Matrix | PNGaseF | Trypsin |
|---|---|---|---|---|---|
| Matrix/Enzyme | DAN | CHCA | CHCA | PNGaseF | Trypsin |
| Concentration (mg/mL) | 10 | 10 | 10 | 0.1 | 0.05 |
| Solvent | 50% ACN | 70% ACN, 0.1% TFA, 10 mM AmPhos | 70% ACN, 0.1% TFA, 10 mM AmPhos | Water | 9% ACN, 100 mM AmBic |
| Number of Passes | 4 | 3 | 4 | 12 | 15 |
| Flow Rate (mL/min) | 0.1 | 0.12 | 0.12 | 0.025 | 0.01 |
| Nozzle Temperature (°C) | 60 | 75 | 75 | 45 | 30 |
| Track Speed (mm/min) | 1200 | 1200 | 1200 | 1200 | 750 |
| Track Spacing (mm) | 3 | 3 | 3 | 3 | 3 |
| Nozzle Height (mm) | 40 | 40 | 40 | 40 | 40 |
| Track Pattern | CC | HH | HH | CC | HH |

DAN – 1,5-diaminonaphthalene, CHCA – α-cyano-4-hydroxycinnamic acid, ACN – acetonitrile, TFA – trifluoroacetic acid, AmPhos – ammonium phosphate, AmBic – ammonium bicarbonate, CC- crisscross, HH – horizontal-horizontal

**Supplemental Table 5. Brain Tumor TMA Layout**

BTB TMA #3 - Gliomas for Xenium (4x10 array)

| | | | |
|---|---|---|---|
| GBM1-Edge | GBM6-Edge | AA1- Edge | Oligo1- Edge |
| GBM1-Core | GBM6-Core | AA1- Core | Oligo1- Core |
| GBM2-Edge | GBM7-Edge | AA2- Edge | Oligo2- Edge |
| GBM2-Core | GBM7-Core | AA2 - Core | Oligo2- Core |
| GBM3-Edge | GBM8-Edge | AA3- Edge | Oligo3- Edge |
| GBM3-Core | GBM8-Core | AA3- Core | Oligo3 - Core |
| GBM4-Edge | GBM9-Edge | OligoGrIII- Edge | Oligo4- Edge |
| GBM4-Core | GBM9-Core | OligoGrIII- Core | Oligo4- Core |
| GBM5-Edge | GBM10-Edge | Control 1 | Oligo5- Edge |
| GBM5-Core | GBM10-Core | Control 2 | Oligo5- Core |

| | | | |
|---|---|---|---|
| 1 | 11 | 21 | 31 |
| 2 | 12 | 22 | 32 |
| 3 | 13 | 23 | 33 |
| 4 | 14 | 24 | 34 |
| 5 | 15 | 25 | 35 |
| 6 | 16 | 26 | 36 |
| 7 | 17 | 27 | 37 |
| 8 | 18 | 28 | 38 |
| 9 | 19 | 29 | 39 |
| 10 | 20 | 30 | 40 |

**Supplemental Table 6. TMA Clinical Information**

| Pathology | DOB | Seizures pre-op | Date of surgery | IDHR132H Mutation | 1p19q co-del | Date of tumor progression | Date of death |
|---|---|---|---|---|---|---|---|
| GBM1 | 7/11/51 | No | 3/28/13 | No | No | 7/28/15 | |
| GBM2 | 10/27/62 | No | 8/1/13 | No | No | 9/19/13 | 7/22/16 |
| GBM3 | 3/5/80 | No | 2/13/14 | No | No | 8/18/15 | |
| GBM4 | 7/7/59 | No | 2/5/15 | No | No | | 5/23/16 |
| GBM5 | 2/16/61 | Yes | 3/17/16 | No | No | 10/16/17 | |
| GBM6 | 10/27/68 | No | 4/4/16 | No | No | 1/8/17 | 9/15/17 |
| GBM7 | 1/21/46 | No | 7/19/16 | No | No | 9/7/17 | 11/4/17 |
| GBM8 | 8/18/46 | No | 7/25/16 | No | No | | 1/22/18 |
| GBM9 | 6/1/54 | No | 7/28/16 | No | No | 9/8/17 | 11/3/17 |
| GBM10 | 1/9/56 | No | 12/21/16 | No | No | 3/23/17 | |
| AA1 | 7/29/88 | Yes | 1/22/18 | Yes | No | 2/10/20 | |
| AA2 | 10/18/81 | No | 1/24/18 | Yes | No | | |
| AA3 | 1/4/90 | Yes | 3/27/18 | Yes | No | | |
| OligGr3 | 6/3/68 | No | 1/16/20 | Yes | Yes | | |
| Olig1 | 4/20/78 | Yes | 10/17/07 | Yes | Yes | | |
| Olig2 | 12/3/74 | No | 8/13/18 | Yes | Yes | 12/28/22 | |
| Olig3 | 8/30/59 | Yes | 9/11/18 | Yes | Yes | | |
| Olig4 | 8/7/81 | Yes | 12/21/18 | Yes | Yes | | |
| Olig5 | 4/21/72 | Yes | 12/5/19 | Yes | Yes | | |
| Control spleen 1 | 7/11/26 | | | | | | |
| Control spleen 2 | 1/18/51 | | | | | | |

**Supplemental Table 7. Spatial transcriptomics derived differentially expressed genes (Oligo)**

| | gene | region | logfoldchange | pval_adj | score |
|---|---|---|---|---|---|
| 0 | PDGFRA | Core | 4.0566297 | 0 | 87.991486 |
| 1 | OLIG2 | Core | 4.248039 | 0 | 72.839645 |
| 2 | OLIG1 | Core | 6.1979194 | 0 | 70.31132 |
| 3 | PTPRZ1 | Core | 7.73765 | 0 | 63.668736 |
| 363 | PON2 | Edge | 3.3614955 | 0 | 57.16311 |
| 362 | EFHD1 | Edge | 1.798796 | 0 | 57.762527 |
| 361 | GJA1 | Edge | 3.1314714 | 0 | 61.622257 |
| 364 | EGFR | Edge | -2.4757304 | 0 | -57.034863 |
| 367 | APOE | Edge | 4.078381 | 0 | 55.214417 |
| 366 | AQP4 | Edge | 5.7990417 | 0 | 55.31656 |
| 365 | SOX11 | Edge | -1.8743243 | 0 | -56.68783 |
| 372 | CCND1 | Edge | -2.269381 | 0 | -48.1786 |
| 371 | TLR4 | Edge | 1.8789823 | 0 | 48.95025 |
| 370 | SOX4 | Edge | -3.251696 | 0 | -51.470585 |
| 369 | KCNJ16 | Edge | 1.8215945 | 0 | 52.42469 |
| 368 | BCAN | Edge | -3.0239496 | 0 | -54.391476 |
| 375 | MYO16 | Edge | -1.5864166 | 0 | -39.697792 |
| 374 | MLC1 | Edge | 1.2804438 | 0 | 44.168354 |
| 373 | MOBP | Edge | 1.8483167 | 0 | 47.166103 |
| 379 | MGST1 | Edge | 1.3186041 | 0 | 37.93247 |
| 376 | MYC | Edge | -1.480059 | 0 | -38.961266 |
| 378 | DNER | Edge | -1.3606641 | 0 | -38.224426 |
| 377 | ADRA1A | Edge | 1.3997303 | 0 | 38.358044 |
| 360 | CD24 | Edge | -2.2846293 | 0 | -61.74372 |
| 380 | ERMN | Edge | 1.8123178 | 0 | 37.795803 |
| 19 | MYO16 | Core | 1.5864166 | 0 | 39.697792 |
| 17 | MOBP | Core | -1.8483167 | 0 | -47.166103 |
| 11 | APOE | Core | -4.078381 | 0 | -55.214417 |
| 356 | PDGFRA | Edge | -4.0566297 | 0 | -87.991486 |
| 357 | OLIG2 | Edge | -4.248039 | 0 | -72.839645 |
| 358 | OLIG1 | Edge | -6.1979194 | 0 | -70.31132 |
| 359 | PTPRZ1 | Edge | -7.73765 | 0 | -63.668736 |
| 12 | BCAN | Core | 3.0239496 | 0 | 54.391476 |
| 13 | KCNJ16 | Core | -1.8215945 | 0 | -52.42469 |
| 14 | SOX4 | Core | 3.251696 | 0 | 51.470585 |
| 15 | TLR4 | Core | -1.8789823 | 0 | -48.95025 |
| 16 | CCND1 | Core | 2.269381 | 0 | 48.1786 |
| 8 | EGFR | Core | 2.4757304 | 0 | 57.034863 |
| 10 | AQP4 | Core | -5.7990417 | 0 | -55.31656 |

| | | | | | |
|---|---|---|---|---|---|
| 23 | MGST1 | Core | -1.3186041 | 0 | -37.93247 |
| 4 | CD24 | Core | 2.2846293 | 0 | 61.74372 |
| 22 | DNER | Core | 1.3606641 | 0 | 38.224426 |
| 21 | ADRA1A | Core | -1.3997303 | 0 | -38.358044 |
| 24 | ERMN | Core | -1.8123178 | 0 | -37.795803 |
| 7 | PON2 | Core | -3.3614955 | 0 | -57.16311 |
| 6 | EFHD1 | Core | -1.798796 | 0 | -57.762527 |
| 5 | GJA1 | Core | -3.1314714 | 0 | -61.622257 |
| 20 | MYC | Core | 1.480059 | 0 | 38.961266 |
| 18 | MLC1 | Core | -1.2804438 | 0 | -44.168354 |
| 9 | SOX11 | Core | 1.8743243 | 0 | 56.68783 |
| 25 | SFRP2 | Core | -1.5053595 | 7.83E-308 | -37.573982 |
| 381 | SFRP2 | Edge | 1.5053595 | 7.83E-308 | 37.573982 |
| 382 | PCSK6 | Edge | 1.4992359 | 4.17E-232 | 32.60795 |
| 26 | PCSK6 | Core | -1.4992359 | 4.17E-232 | -32.60795 |
| 27 | FGFR3 | Core | -1.5817012 | 3.46E-230 | -32.471153 |
| 383 | FGFR3 | Edge | 1.5817012 | 3.46E-230 | 32.471153 |
| 384 | TP53 | Edge | -0.9712106 | 4.15E-222 | -31.892582 |
| 28 | TP53 | Core | 0.9712106 | 4.15E-222 | 31.892582 |
| 29 | IDH1 | Core | 1.0421216 | 1.41E-213 | 31.270224 |
| 385 | IDH1 | Edge | -1.0421216 | 1.41E-213 | -31.270224 |
| 386 | UGT8 | Edge | -0.7817959 | 3.05E-204 | -30.574703 |
| 30 | UGT8 | Core | 0.78179586 | 3.05E-204 | 30.574703 |
| 31 | PAX6 | Core | -1.1657003 | 2.37E-201 | -30.355404 |
| 387 | PAX6 | Edge | 1.1657003 | 2.37E-201 | 30.355404 |
| 32 | FGFR2 | Core | -1.4543738 | 6.20E-195 | -29.864176 |
| 388 | FGFR2 | Edge | 1.4543738 | 6.20E-195 | 29.864176 |
| 389 | VCAN | Edge | -0.861787 | 6.32E-193 | -29.708118 |
| 33 | VCAN | Core | 0.86178696 | 6.32E-193 | 29.708118 |
| 34 | THY1 | Core | 0.4615311 | 5.04E-190 | 29.481604 |
| 390 | THY1 | Edge | -0.4615311 | 5.04E-190 | -29.481604 |
| 391 | MYCN | Edge | -1.891814 | 1.51E-176 | -28.409842 |
| 35 | MYCN | Core | 1.891814 | 1.51E-176 | 28.409842 |
| 36 | GAD1 | Core | 1.1747823 | 8.72E-173 | 28.102747 |
| 392 | GAD1 | Edge | -1.1747823 | 8.72E-173 | -28.102747 |
| 393 | SPON1 | Edge | 0.8794924 | 6.55E-170 | 27.865496 |
| 37 | SPON1 | Core | -0.8794924 | 6.55E-170 | -27.865496 |
| 38 | SOX9 | Core | -0.7686822 | 5.30E-167 | -27.623547 |
| 394 | SOX9 | Edge | 0.7686822 | 5.30E-167 | 27.623547 |
| 395 | GPR37L1 | Edge | -0.9202901 | 1.52E-166 | -27.584438 |
| 39 | GPR37L1 | Core | 0.9202901 | 1.52E-166 | 27.584438 |

| | | | | | |
|---|---|---|---|---|---|
| 396 | PTCH1 | Edge | -1.3703552 | 3.63E-158 | -26.876116 |
| 40 | PTCH1 | Core | 1.3703552 | 3.63E-158 | 26.876116 |
| 41 | CDK6 | Core | 1.3752836 | 3.74E-158 | 26.874111 |
| 397 | CDK6 | Edge | -1.3752836 | 3.74E-158 | -26.874111 |
| 42 | MOG | Core | -1.627679 | 1.33E-154 | -26.567612 |
| 398 | MOG | Edge | 1.627679 | 1.33E-154 | 26.567612 |
| 399 | CD44 | Edge | 0.5585383 | 2.31E-144 | 25.665268 |
| 43 | CD44 | Core | -0.5585383 | 2.31E-144 | -25.665268 |
| 44 | P2RY12 | Core | -1.2840438 | 2.02E-142 | -25.48987 |
| 400 | P2RY12 | Edge | 1.2840438 | 2.02E-142 | 25.48987 |
| 401 | SMARCB1 | Edge | -0.7479384 | 1.77E-129 | -24.29356 |
| 45 | SMARCB1 | Core | 0.7479384 | 1.77E-129 | 24.29356 |
| 46 | CAMK2A | Core | -1.7194096 | 1.42E-123 | -23.72747 |
| 402 | CAMK2A | Edge | 1.7194096 | 1.42E-123 | 23.72747 |
| 403 | CCND2 | Edge | -0.6051589 | 1.44E-123 | -23.726194 |
| 47 | CCND2 | Core | 0.6051589 | 1.44E-123 | 23.726194 |
| 48 | HEPACAM | Core | -0.8832935 | 1.11E-116 | -23.048244 |
| 404 | HEPACAM | Edge | 0.8832935 | 1.11E-116 | 23.048244 |
| 405 | CLDN11 | Edge | 1.3780439 | 6.81E-100 | 21.308052 |
| 49 | CLDN11 | Core | -1.3780439 | 6.81E-100 | -21.308052 |
| 50 | TTYH1 | Core | -0.6959034 | 2.26E-91 | -20.367558 |
| 406 | TTYH1 | Edge | 0.6959034 | 2.26E-91 | 20.367558 |
| 407 | AIF1 | Edge | 0.8046593 | 9.68E-87 | 19.837448 |
| 51 | AIF1 | Core | -0.8046593 | 9.68E-87 | -19.837448 |
| 52 | ELOVL2 | Core | -1.2795196 | 1.60E-86 | -19.811249 |
| 408 | ELOVL2 | Edge | 1.2795196 | 1.60E-86 | 19.811249 |
| 409 | S100A1 | Edge | 1.091353 | 5.94E-85 | 19.627436 |
| 53 | S100A1 | Core | -1.091353 | 5.94E-85 | -19.627436 |
| 54 | APP | Core | 0.6363028 | 1.77E-83 | 19.453375 |
| 410 | APP | Edge | -0.6363028 | 1.77E-83 | -19.453375 |
| 411 | CAPN3 | Edge | 1.4679103 | 2.19E-80 | 19.083755 |
| 55 | CAPN3 | Core | -1.4679103 | 2.19E-80 | -19.083755 |
| 56 | GPR34 | Core | -0.8739823 | 5.37E-79 | -18.91486 |
| 412 | GPR34 | Edge | 0.8739823 | 5.37E-79 | 18.91486 |
| 413 | TRIL | Edge | 0.69460905 | 7.40E-79 | 18.897043 |
| 57 | TRIL | Core | -0.6946091 | 7.40E-79 | -18.897043 |
| 58 | CNDP1 | Core | -1.3422494 | 1.59E-70 | -17.855024 |
| 414 | CNDP1 | Edge | 1.3422494 | 1.59E-70 | 17.855024 |
| 415 | RASGRP1 | Edge | 1.1808763 | 1.60E-70 | 17.85374 |
| 59 | RASGRP1 | Core | -1.1808763 | 1.60E-70 | -17.85374 |
| 60 | MAL | Core | -1.3017688 | 1.17E-68 | -17.611626 |

| | | | | | |
|---|---|---|---|---|---|
| 416 | MAL | Edge | 1.3017688 | 1.17E-68 | 17.611626 |
| 417 | CALCRL | Edge | -0.4016616 | 1.19E-61 | -16.672562 |
| 61 | CALCRL | Core | 0.40166163 | 1.19E-61 | 16.672562 |
| 62 | ARHGAP24 | Core | 0.7169662 | 7.56E-60 | 16.421686 |
| 418 | ARHGAP24 | Edge | -0.7169662 | 7.56E-60 | -16.421686 |
| 419 | SLC24A3 | Edge | -0.5458621 | 3.27E-59 | -16.33158 |
| 63 | SLC24A3 | Core | 0.54586214 | 3.27E-59 | 16.33158 |
| 420 | TGFB2 | Edge | 0.923367 | 1.79E-56 | 15.941347 |
| 64 | TGFB2 | Core | -0.923367 | 1.79E-56 | -15.941347 |
| 65 | MAG | Core | -1.5642161 | 3.66E-52 | -15.307751 |
| 421 | MAG | Edge | 1.5642161 | 3.66E-52 | 15.307751 |
| 422 | BRINP3 | Edge | -0.4950482 | 1.38E-50 | -15.068758 |
| 66 | BRINP3 | Core | 0.4950482 | 1.38E-50 | 15.068758 |
| 67 | SETD2 | Core | -0.4414551 | 1.97E-49 | -14.891025 |
| 423 | SETD2 | Edge | 0.44145513 | 1.97E-49 | 14.891025 |
| 424 | CXCL14 | Edge | 0.8072389 | 2.35E-49 | 14.878227 |
| 68 | CXCL14 | Core | -0.8072389 | 2.35E-49 | -14.878227 |
| 69 | NCAM1 | Core | 0.5370375 | 1.82E-47 | 14.583583 |
| 425 | NCAM1 | Edge | -0.5370375 | 1.82E-47 | -14.583583 |
| 426 | RNASET2 | Edge | 0.7379525 | 3.31E-47 | 14.541581 |
| 70 | RNASET2 | Core | -0.7379525 | 3.31E-47 | -14.541581 |
| 71 | PTCHD4 | Core | -1.3168108 | 9.85E-47 | -14.465809 |
| 427 | PTCHD4 | Edge | 1.3168108 | 9.85E-47 | 14.465809 |
| 428 | SERPINA3 | Edge | -1.6779081 | 2.33E-45 | -14.245506 |
| 72 | SERPINA3 | Core | 1.6779081 | 2.33E-45 | 14.245506 |
| 73 | SLC17A7 | Core | -1.0645193 | 4.56E-45 | -14.197562 |
| 429 | SLC17A7 | Edge | 1.0645193 | 4.56E-45 | 14.197562 |
| 430 | VIM | Edge | 0.5648302 | 5.95E-45 | 14.178042 |
| 74 | VIM | Core | -0.5648302 | 5.95E-45 | -14.178042 |
| 75 | AXL | Core | -0.6793969 | 1.67E-44 | -14.10438 |
| 431 | AXL | Edge | 0.67939687 | 1.67E-44 | 14.10438 |
| 76 | IGFBP5 | Core | -0.6369706 | 7.68E-44 | -13.995515 |
| 432 | IGFBP5 | Edge | 0.63697064 | 7.68E-44 | 13.995515 |
| 433 | VCP | Edge | -0.3685866 | 1.34E-43 | -13.955137 |
| 77 | VCP | Core | 0.3685866 | 1.34E-43 | 13.955137 |
| 78 | MDM4 | Core | 0.37982392 | 2.44E-43 | 13.911099 |
| 434 | MDM4 | Edge | -0.3798239 | 2.44E-43 | -13.911099 |
| 435 | RORB | Edge | 0.77711296 | 5.97E-43 | 13.846238 |
| 79 | RORB | Core | -0.777113 | 5.97E-43 | -13.846238 |
| 80 | ABCC9 | Core | -0.7167785 | 8.30E-43 | -13.821591 |
| 436 | ABCC9 | Edge | 0.71677846 | 8.30E-43 | 13.821591 |

| | | | | | |
|---|---|---|---|---|---|
| 437 | CNTN2 | Edge | 1.17741 | 9.61E-40 | 13.303375 |
| 81 | CNTN2 | Core | -1.17741 | 9.61E-40 | -13.303375 |
| 82 | ST18 | Core | -1.4184465 | 2.22E-39 | -13.239597 |
| 438 | ST18 | Edge | 1.4184465 | 2.22E-39 | 13.239597 |
| 439 | IDH2 | Edge | 0.45280808 | 1.21E-37 | 12.934953 |
| 83 | IDH2 | Core | -0.4528081 | 1.21E-37 | -12.934953 |
| 84 | CCK | Core | -1.598412 | 5.60E-35 | -12.453573 |
| 440 | CCK | Edge | 1.598412 | 5.60E-35 | 12.453573 |
| 441 | CUX2 | Edge | -1.2318517 | 1.22E-33 | -12.204618 |
| 85 | CUX2 | Core | 1.2318517 | 1.22E-33 | 12.204618 |
| 86 | MSH6 | Core | 0.5462543 | 6.43E-33 | 12.06728 |
| 442 | MSH6 | Edge | -0.5462543 | 6.43E-33 | -12.06728 |
| 443 | IFITM3 | Edge | 0.3040228 | 7.03E-33 | 12.059098 |
| 87 | IFITM3 | Core | -0.3040228 | 7.03E-33 | -12.059098 |
| 88 | IGFBP4 | Core | 0.5441856 | 1.18E-32 | 12.015107 |
| 444 | IGFBP4 | Edge | -0.5441856 | 1.18E-32 | -12.015107 |
| 89 | HHATL | Core | -1.2257978 | 5.64E-32 | -11.884492 |
| 445 | HHATL | Edge | 1.2257978 | 5.64E-32 | 11.884492 |
| 90 | B4GALNT1 | Core | 0.45049182 | 4.21E-29 | 11.317379 |
| 446 | B4GALNT1 | Edge | -0.4504918 | 4.21E-29 | -11.317379 |
| 447 | OPALIN | Edge | 0.90561795 | 7.84E-29 | 11.261748 |
| 91 | OPALIN | Core | -0.905618 | 7.84E-29 | -11.261748 |
| 93 | STAT3 | Core | 0.314278 | 1.74E-28 | 11.189619 |
| 92 | HTR2A | Core | -0.8627183 | 1.74E-28 | -11.190133 |
| 448 | HTR2A | Edge | 0.86271834 | 1.74E-28 | 11.190133 |
| 449 | STAT3 | Edge | -0.314278 | 1.74E-28 | -11.189619 |
| 94 | CAMK2D | Core | 0.24222682 | 1.63E-27 | 10.9881735 |
| 450 | CAMK2D | Edge | -0.2422268 | 1.63E-27 | -10.988174 |
| 451 | RYR1 | Edge | 1.2344826 | 5.79E-27 | 10.8723135 |
| 95 | RYR1 | Core | -1.2344826 | 5.79E-27 | -10.872314 |
| 96 | CX3CR1 | Core | -1.2027664 | 1.09E-25 | -10.600478 |
| 452 | CX3CR1 | Edge | 1.2027664 | 1.09E-25 | 10.600478 |
| 453 | HTRA1 | Edge | -0.4537197 | 1.27E-25 | -10.585029 |
| 97 | HTRA1 | Core | 0.4537197 | 1.27E-25 | 10.585029 |
| 98 | CTSH | Core | -0.6461533 | 1.79E-25 | -10.551919 |
| 454 | CTSH | Edge | 0.6461533 | 1.79E-25 | 10.551919 |
| 100 | NOTCH1 | Core | 0.2003388 | 3.91E-24 | 10.256241 |
| 99 | TENM1 | Core | 0.725969 | 3.91E-24 | 10.256999 |
| 456 | NOTCH1 | Edge | -0.2003388 | 3.91E-24 | -10.256241 |
| 455 | TENM1 | Edge | -0.725969 | 3.91E-24 | -10.256999 |
| 101 | TSPAN7 | Core | 0.13281922 | 9.20E-23 | 9.945594 |

| | | | | | |
|---|---|---|---|---|---|
| 457 | TSPAN7 | Edge | -0.1328192 | 9.20E-23 | -9.945594 |
| 458 | EZH2 | Edge | -0.6662955 | 2.24E-22 | -9.855586 |
| 102 | EZH2 | Core | 0.6662955 | 2.24E-22 | 9.855586 |
| 103 | HES1 | Core | -0.6321654 | 5.83E-21 | -9.521676 |
| 459 | HES1 | Edge | 0.6321654 | 5.83E-21 | 9.521676 |
| 460 | ITGA7 | Edge | 0.6019212 | 1.51E-20 | 9.421067 |
| 104 | ITGA7 | Core | -0.6019212 | 1.51E-20 | -9.421067 |
| 105 | RGS10 | Core | -0.4785325 | 3.30E-20 | -9.338015 |
| 461 | RGS10 | Edge | 0.4785325 | 3.30E-20 | 9.338015 |
| 462 | ERBB2 | Edge | -0.475886 | 1.13E-19 | -9.206053 |
| 106 | ERBB2 | Core | 0.475886 | 1.13E-19 | 9.206053 |
| 107 | ERBB3 | Core | 0.29014492 | 3.56E-19 | 9.080614 |
| 463 | ERBB3 | Edge | -0.2901449 | 3.56E-19 | -9.080614 |
| 464 | MYRF | Edge | 1.0093244 | 4.13E-18 | 8.808866 |
| 108 | MYRF | Core | -1.0093244 | 4.13E-18 | -8.808866 |
| 109 | HHIP | Core | -0.8501227 | 6.12E-18 | -8.763632 |
| 465 | HHIP | Edge | 0.8501227 | 6.12E-18 | 8.763632 |
| 466 | PLCH1 | Edge | 0.82231426 | 9.04E-17 | 8.453814 |
| 110 | PLCH1 | Core | -0.8223143 | 9.04E-17 | -8.453814 |
| 111 | DDR2 | Core | -0.7309706 | 1.04E-16 | -8.43629 |
| 467 | DDR2 | Edge | 0.7309706 | 1.04E-16 | 8.43629 |
| 468 | CXCR4 | Edge | 0.82059395 | 1.18E-16 | 8.420541 |
| 112 | CXCR4 | Core | -0.820594 | 1.18E-16 | -8.420541 |
| 113 | C1orf162 | Core | -0.703684 | 1.49E-16 | -8.39208 |
| 469 | C1orf162 | Edge | 0.703684 | 1.49E-16 | 8.39208 |
| 470 | FLT1 | Edge | 0.623051 | 1.01E-15 | 8.163522 |
| 114 | FLT1 | Core | -0.623051 | 1.01E-15 | -8.163522 |
| 115 | FCGR1A | Core | -0.6692705 | 3.32E-15 | -8.017186 |
| 471 | FCGR1A | Edge | 0.66927046 | 3.32E-15 | 8.017186 |
| 472 | PCNA | Edge | -0.3485926 | 3.62E-15 | -8.005352 |
| 116 | PCNA | Core | 0.34859258 | 3.62E-15 | 8.005352 |
| 117 | SCN8A | Core | 0.13183023 | 1.08E-14 | 7.868889 |
| 473 | SCN8A | Edge | -0.1318302 | 1.08E-14 | -7.868889 |
| 474 | NTNG2 | Edge | -0.2521243 | 2.47E-14 | -7.763464 |
| 118 | NTNG2 | Core | 0.25212425 | 2.47E-14 | 7.763464 |
| 119 | P2RY13 | Core | -1.1039401 | 3.96E-14 | -7.7025247 |
| 475 | P2RY13 | Edge | 1.1039401 | 3.96E-14 | 7.7025247 |
| 476 | CTSS | Edge | 0.5973101 | 4.09E-14 | 7.69717 |
| 120 | CTSS | Core | -0.5973101 | 4.09E-14 | -7.69717 |
| 121 | MS4A6A | Core | -0.5569366 | 1.33E-13 | -7.544244 |
| 477 | MS4A6A | Edge | 0.55693656 | 1.33E-13 | 7.544244 |

| | | | | | |
|---|---|---|---|---|---|
| 478 | TSHZ2 | Edge | 0.37813112 | 6.24E-13 | 7.338824 |
| 122 | TSHZ2 | Core | -0.3781311 | 6.24E-13 | -7.338824 |
| 123 | HLA-DMB | Core | -0.599632 | 7.59E-13 | -7.3113265 |
| 479 | HLA-DMB | Edge | 0.59963197 | 7.59E-13 | 7.3113265 |
| 480 | CD68 | Edge | 0.32455343 | 1.03E-12 | 7.2692223 |
| 124 | CD68 | Core | -0.3245534 | 1.03E-12 | -7.2692223 |
| 125 | DCN | Core | -0.7033809 | 1.62E-12 | -7.206981 |
| 481 | DCN | Edge | 0.7033809 | 1.62E-12 | 7.206981 |
| 482 | HS3ST4 | Edge | 1.405675 | 2.02E-12 | 7.1753554 |
| 126 | HS3ST4 | Core | -1.405675 | 2.02E-12 | -7.1753554 |
| 127 | FBLN1 | Core | -0.9262114 | 2.54E-12 | -7.142962 |
| 483 | FBLN1 | Edge | 0.9262114 | 2.54E-12 | 7.142962 |
| 484 | LY86 | Edge | 0.6435516 | 4.13E-12 | 7.074936 |
| 128 | LY86 | Core | -0.6435516 | 4.13E-12 | -7.074936 |
| 129 | BGN | Core | -0.8293523 | 6.61E-12 | -7.0081787 |
| 485 | BGN | Edge | 0.82935226 | 6.61E-12 | 7.0081787 |
| 486 | BMI1 | Edge | -0.2451487 | 1.82E-11 | -6.864224 |
| 130 | BMI1 | Core | 0.24514869 | 1.82E-11 | 6.864224 |
| 132 | TESPA1 | Core | -2.0282686 | 2.18E-11 | -6.835811 |
| 131 | SYNPR | Core | -1.3327714 | 2.18E-11 | -6.836518 |
| 488 | TESPA1 | Edge | 2.0282686 | 2.18E-11 | 6.835811 |
| 487 | SYNPR | Edge | 1.3327714 | 2.18E-11 | 6.836518 |
| 134 | TMEM132C | Core | 0.49020058 | 2.21E-11 | 6.8319645 |
| 133 | CNTNAP3B | Core | -0.6314933 | 2.21E-11 | -6.832376 |
| 489 | CNTNAP3B | Edge | 0.6314933 | 2.21E-11 | 6.832376 |
| 490 | TMEM132C | Edge | -0.4902006 | 2.21E-11 | -6.8319645 |
| 135 | SOX2 | Core | 0.13175757 | 3.28E-11 | 6.773901 |
| 491 | SOX2 | Edge | -0.1317576 | 3.28E-11 | -6.773901 |
| 136 | CENPF | Core | 0.9379881 | 5.59E-11 | 6.695335 |
| 492 | CENPF | Edge | -0.9379881 | 5.59E-11 | -6.695335 |
| 493 | LYVE1 | Edge | 0.70199114 | 6.46E-11 | 6.6730423 |
| 137 | LYVE1 | Core | -0.7019911 | 6.46E-11 | -6.6730423 |
| 138 | LAMP5 | Core | -0.6023358 | 8.86E-11 | -6.625591 |
| 494 | LAMP5 | Edge | 0.6023358 | 8.86E-11 | 6.625591 |
| 495 | NCSTN | Edge | -0.2185124 | 2.13E-10 | -6.4937673 |
| 139 | NCSTN | Core | 0.21851236 | 2.13E-10 | 6.4937673 |
| 496 | TGFB1 | Edge | 0.22966553 | 3.02E-10 | 6.439768 |
| 140 | TGFB1 | Core | -0.2296655 | 3.02E-10 | -6.439768 |
| 141 | FGF13 | Core | 0.1439184 | 1.03E-09 | 6.2498183 |
| 497 | FGF13 | Edge | -0.1439184 | 1.03E-09 | -6.2498183 |
| 498 | SULF1 | Edge | 0.5458159 | 1.52E-09 | 6.1872444 |

| | | | | | |
|---|---|---|---|---|---|
| 142 | SULF1 | Core | -0.5458159 | 1.52E-09 | -6.1872444 |
| 143 | KIT | Core | -0.5201916 | 2.66E-09 | -6.0978093 |
| 499 | KIT | Edge | 0.5201916 | 2.66E-09 | 6.0978093 |
| 500 | SPHKAP | Edge | -0.303584 | 2.90E-09 | -6.082635 |
| 144 | SPHKAP | Core | 0.30358395 | 2.90E-09 | 6.082635 |
| 145 | SOX10 | Core | -0.8605683 | 5.41E-09 | -5.980886 |
| 501 | SOX10 | Edge | 0.86056834 | 5.41E-09 | 5.980886 |
| 502 | ITGB2 | Edge | 0.39097655 | 5.75E-09 | 5.9698954 |
| 146 | ITGB2 | Core | -0.3909766 | 5.75E-09 | -5.9698954 |
| 147 | OTOGL | Core | 0.59693766 | 5.97E-09 | 5.9625344 |
| 503 | OTOGL | Edge | -0.5969377 | 5.97E-09 | -5.9625344 |
| 504 | FGFR1 | Edge | -0.1172851 | 6.25E-09 | -5.9541335 |
| 148 | FGFR1 | Core | 0.1172851 | 6.25E-09 | 5.9541335 |
| 149 | PECAM1 | Core | -0.2615408 | 7.88E-09 | -5.9148946 |
| 505 | PECAM1 | Edge | 0.2615408 | 7.88E-09 | 5.9148946 |
| 506 | TMIGD3 | Edge | 0.25010225 | 1.11E-08 | 5.8569837 |
| 150 | TMIGD3 | Core | -0.2501023 | 1.11E-08 | -5.8569837 |
| 151 | SEMA5A | Core | -0.5212941 | 2.04E-08 | -5.7538967 |
| 507 | SEMA5A | Edge | 0.5212941 | 2.04E-08 | 5.7538967 |
| 508 | PDGFRB | Edge | 0.3183614 | 2.05E-08 | 5.752584 |
| 152 | PDGFRB | Core | -0.3183614 | 2.05E-08 | -5.752584 |
| 153 | PTPN12 | Core | -0.1914539 | 3.55E-08 | -5.657654 |
| 509 | PTPN12 | Edge | 0.19145393 | 3.55E-08 | 5.657654 |
| 510 | NRP1 | Edge | 0.47026253 | 4.28E-08 | 5.6241665 |
| 154 | NRP1 | Core | -0.4702625 | 4.28E-08 | -5.6241665 |
| 155 | RYR2 | Core | -1.0101963 | 5.83E-08 | -5.5694156 |
| 511 | RYR2 | Edge | 1.0101963 | 5.83E-08 | 5.5694156 |
| 512 | TOP2A | Edge | -0.6285361 | 2.54E-07 | -5.3061137 |
| 156 | TOP2A | Core | 0.6285361 | 2.54E-07 | 5.3061137 |
| 157 | ITGAM | Core | -0.775946 | 3.10E-07 | -5.2684503 |
| 513 | ITGAM | Edge | 0.775946 | 3.10E-07 | 5.2684503 |
| 514 | ANGPT1 | Edge | 0.53262705 | 7.78E-07 | 5.095779 |
| 158 | ANGPT1 | Core | -0.5326271 | 7.78E-07 | -5.095779 |
| 159 | PROX1 | Core | 0.15811184 | 7.89E-07 | 5.0917788 |
| 515 | PROX1 | Edge | -0.1581118 | 7.89E-07 | -5.0917788 |
| 516 | ANK1 | Edge | -0.5411641 | 1.13E-06 | -5.0227766 |
| 160 | ANK1 | Core | 0.5411641 | 1.13E-06 | 5.0227766 |
| 161 | RGS16 | Core | 1.0252002 | 8.35E-06 | 4.6220713 |
| 517 | RGS16 | Edge | -1.0252002 | 8.35E-06 | -4.6220713 |
| 518 | LYPD6 | Edge | 0.79299533 | 1.29E-05 | 4.5300813 |
| 162 | LYPD6 | Core | -0.7929953 | 1.29E-05 | -4.5300813 |

| | | | | | |
|---|---|---|---|---|---|
| 163 | LAMA2 | Core | -0.6403815 | 1.59E-05 | -4.4842267 |
| 519 | LAMA2 | Edge | 0.6403815 | 1.59E-05 | 4.4842267 |
| 520 | CDK1 | Edge | -0.8229792 | 1.85E-05 | -4.449952 |
| 164 | CDK1 | Core | 0.8229792 | 1.85E-05 | 4.449952 |
| 165 | HLA-DQA1 | Core | -0.6439404 | 2.12E-05 | -4.4201326 |
| 521 | HLA-DQA1 | Edge | 0.6439404 | 2.12E-05 | 4.4201326 |
| 522 | PTPRC | Edge | 0.3810481 | 2.17E-05 | 4.4136834 |
| 166 | PTPRC | Core | -0.3810481 | 2.17E-05 | -4.4136834 |
| 167 | FCGR3A | Core | -0.3679986 | 3.91E-05 | -4.28261 |
| 523 | FCGR3A | Edge | 0.36799863 | 3.91E-05 | 4.28261 |
| 524 | VCAM1 | Edge | 0.81351614 | 4.59E-05 | 4.2457294 |
| 168 | VCAM1 | Core | -0.8135161 | 4.59E-05 | -4.2457294 |
| 169 | RIT2 | Core | 0.43888137 | 5.59E-05 | 4.200081 |
| 525 | RIT2 | Edge | -0.4388814 | 5.59E-05 | -4.200081 |
| 526 | NTNG1 | Edge | 0.54750955 | 6.34E-05 | 4.1700816 |
| 170 | NTNG1 | Core | -0.5475096 | 6.34E-05 | -4.1700816 |
| 171 | FCER1G | Core | 0.13079716 | 7.61E-05 | -4.1269135 |
| 527 | FCER1G | Edge | -0.1307972 | 7.61E-05 | 4.1269135 |
| 528 | CD86 | Edge | 0.40498534 | 8.62E-05 | 4.0968533 |
| 172 | CD86 | Core | -0.4049853 | 8.62E-05 | -4.0968533 |
| 173 | PSENEN | Core | -0.2506106 | 9.16E-05 | -4.081353 |
| 529 | PSENEN | Edge | 0.2506106 | 9.16E-05 | 4.081353 |
| 530 | KRAS | Edge | -0.0054834 | 0.00013005 | -3.9978242 |
| 174 | KRAS | Core | 0.00548338 | 0.00013005 | 3.9978242 |
| 175 | CRYM | Core | -0.8828223 | 0.0001409 | -3.977454 |
| 531 | CRYM | Edge | 0.8828223 | 0.0001409 | 3.977454 |
| 532 | ANKRD18A | Edge | 0.6905419 | 0.0001935 | 3.899964 |
| 176 | ANKRD18A | Core | -0.6905419 | 0.0001935 | -3.899964 |
| 177 | VWC2L | Core | 0.21103227 | 0.00033454 | 3.7639227 |
| 533 | VWC2L | Edge | -0.2110323 | 0.00033454 | -3.7639227 |
| 178 | RYR3 | Core | -0.6203403 | 0.00052657 | -3.6475382 |
| 534 | RYR3 | Edge | 0.6203403 | 0.00052657 | 3.6475382 |
| 535 | RGS5 | Edge | 0.14223683 | 0.0007079 | 3.5693069 |
| 179 | RGS5 | Core | -0.1422368 | 0.0007079 | -3.5693069 |
| 180 | ITGAX | Core | -0.1531036 | 0.00082455 | -3.5276766 |
| 536 | ITGAX | Edge | 0.1531036 | 0.00082455 | 3.5276766 |

**Supplemental Table 7. Spatial transcriptomics derived differentially expressed genes (AA)**

| | gene | region | logfoldchange | pval_adj | score |
|---|---|---|---|---|---|
| 0 | CAMK2A | Core | -9.177222 | 0 | -41.541985 |
| 361 | CAMK2A | Edge | 9.177222 | 0 | 41.541985 |
| 362 | THY1 | Edge | 4.011147 | 6.46E-283 | 36.08737 |
| 1 | THY1 | Core | -4.011147 | 6.46E-283 | -36.08737 |
| 2 | SLC17A7 | Core | -6.8036795 | 1.71E-254 | -34.215572 |
| 363 | SLC17A7 | Edge | 6.8036795 | 1.71E-254 | 34.215572 |
| 364 | FGF13 | Edge | 2.7551522 | 6.66E-240 | 33.211365 |
| 3 | FGF13 | Core | -2.7551522 | 6.66E-240 | -33.211365 |
| 4 | APOE | Core | 7.943713 | 2.27E-204 | 30.644318 |
| 365 | APOE | Edge | -7.943713 | 2.27E-204 | -30.644318 |
| 366 | GPNMB | Edge | -4.0283017 | 6.04E-184 | -29.064655 |
| 5 | GPNMB | Core | 4.0283017 | 6.04E-184 | 29.064655 |
| 6 | CCK | Core | -4.468705 | 1.77E-162 | -27.3078 |
| 367 | CCK | Edge | 4.468705 | 1.77E-162 | 27.3078 |
| 368 | CCND2 | Edge | 2.4456341 | 1.48E-160 | 27.140455 |
| 7 | CCND2 | Core | -2.4456341 | 1.48E-160 | -27.140455 |
| 8 | CXCL14 | Core | -4.6131606 | 8.81E-152 | -26.382126 |
| 369 | CXCL14 | Edge | 4.6131606 | 8.81E-152 | 26.382126 |
| 9 | MS4A6A | Core | 2.0964823 | 1.25E-132 | 24.652477 |
| 370 | MS4A6A | Edge | -2.0964823 | 1.25E-132 | -24.652477 |
| 371 | LAMP5 | Edge | 5.0937304 | 3.25E-113 | 22.766285 |
| 10 | LAMP5 | Core | -5.0937304 | 3.25E-113 | -22.766285 |
| 11 | TESPA1 | Core | -5.711076 | 1.96E-102 | -21.646864 |
| 372 | TESPA1 | Edge | 5.711076 | 1.96E-102 | 21.646864 |
| 373 | CD24 | Edge | -1.8156096 | 1.22E-97 | -21.128147 |
| 12 | CD24 | Core | 1.8156096 | 1.22E-97 | 21.128147 |
| 13 | CUX2 | Core | -4.197691 | 3.97E-90 | -20.291136 |
| 374 | CUX2 | Edge | 4.197691 | 3.97E-90 | 20.291136 |
| 375 | VCAN | Edge | -2.1143625 | 5.64E-86 | -19.812086 |
| 14 | VCAN | Core | 2.1143625 | 5.64E-86 | 19.812086 |
| 15 | CRYM | Core | -4.298945 | 2.57E-74 | -18.407629 |
| 376 | CRYM | Edge | 4.298945 | 2.57E-74 | 18.407629 |
| 377 | VIM | Edge | -3.412486 | 2.23E-71 | -18.034075 |
| 16 | VIM | Core | 3.412486 | 2.23E-71 | 18.034075 |
| 17 | BCAN | Core | 3.5615184 | 8.46E-69 | 17.699657 |
| 378 | BCAN | Edge | -3.5615184 | 8.46E-69 | -17.699657 |
| 379 | SCN8A | Edge | 2.9474914 | 7.79E-67 | 17.440027 |
| 18 | SCN8A | Core | -2.9474914 | 7.79E-67 | -17.440027 |
| 19 | NNAT | Core | -3.7677417 | 1.11E-66 | -17.416708 |

| | | | | | |
|---|---|---|---|---|---|
| 380 | NNAT | Edge | 3.7677417 | 1.11E-66 | 17.416708 |
| 381 | IGFBP5 | Edge | 2.1559207 | 1.12E-65 | 17.281313 |
| 20 | IGFBP5 | Core | -2.1559207 | 1.12E-65 | -17.281313 |
| 21 | CD163 | Core | 1.449369 | 7.90E-65 | 17.16552 |
| 382 | CD163 | Edge | -1.449369 | 7.90E-65 | -17.16552 |
| 383 | MYC | Edge | -1.8593186 | 1.03E-63 | -17.01295 |
| 22 | MYC | Core | 1.8593186 | 1.03E-63 | 17.01295 |
| 23 | C1QL3 | Core | -4.718191 | 1.02E-62 | -16.875584 |
| 384 | C1QL3 | Edge | 4.718191 | 1.02E-62 | 16.875584 |
| 385 | SYNPR | Edge | 2.5074909 | 4.25E-61 | 16.651686 |
| 24 | SYNPR | Core | -2.5074909 | 4.25E-61 | -16.651686 |
| 25 | ANXA1 | Core | 1.3851919 | 4.81E-60 | 16.50343 |
| 386 | ANXA1 | Edge | -1.3851919 | 4.81E-60 | -16.50343 |
| 387 | CTSS | Edge | -1.4428506 | 3.59E-56 | -15.953759 |
| 26 | CTSS | Core | 1.4428506 | 3.59E-56 | 15.953759 |
| 27 | B4GALNT1 | Core | -1.8423593 | 2.70E-55 | -15.82491 |
| 388 | B4GALNT1 | Edge | 1.8423593 | 2.70E-55 | 15.82491 |
| 389 | NCAM1 | Edge | 1.2193859 | 3.73E-54 | 15.656563 |
| 28 | NCAM1 | Core | -1.2193859 | 3.73E-54 | -15.656563 |
| 29 | IDH2 | Core | -0.9266681 | 5.60E-52 | -15.332272 |
| 390 | IDH2 | Edge | 0.92666805 | 5.60E-52 | 15.332272 |
| 391 | TSPAN7 | Edge | 3.5309918 | 4.82E-51 | 15.189673 |
| 30 | TSPAN7 | Core | -3.5309918 | 4.82E-51 | -15.189673 |
| 31 | PDGFRA | Core | 1.0558863 | 3.96E-50 | 15.048877 |
| 392 | PDGFRA | Edge | -1.0558863 | 3.96E-50 | -15.048877 |
| 393 | WIF1 | Edge | 3.4387546 | 9.11E-46 | 14.367085 |
| 32 | WIF1 | Core | -3.4387546 | 9.11E-46 | -14.367085 |
| 33 | BTBD11 | Core | -1.8741666 | 1.36E-45 | -14.337421 |
| 394 | BTBD11 | Edge | 1.8741666 | 1.36E-45 | 14.337421 |
| 34 | BMI1 | Core | -0.8318576 | 5.64E-45 | -14.236105 |
| 395 | BMI1 | Edge | 0.8318576 | 5.64E-45 | 14.236105 |
| 35 | SERPINA3 | Core | 0.9536019 | 1.20E-42 | 13.854622 |
| 396 | SERPINA3 | Edge | -0.9536019 | 1.20E-42 | -13.854622 |
| 36 | SNTB2 | Core | -1.3348029 | 1.54E-41 | -13.668024 |
| 397 | SNTB2 | Edge | 1.3348029 | 1.54E-41 | 13.668024 |
| 37 | SEMA5A | Core | 0.8371093 | 1.60E-41 | 13.663335 |
| 398 | SEMA5A | Edge | -0.8371093 | 1.60E-41 | -13.663335 |
| 399 | VCP | Edge | 0.8414543 | 3.71E-41 | 13.599903 |
| 38 | VCP | Core | -0.8414543 | 3.71E-41 | -13.599903 |
| 400 | CTSH | Edge | -1.0953193 | 1.76E-38 | -13.139862 |
| 39 | CTSH | Core | 1.0953193 | 1.76E-38 | 13.139862 |

| | | | | | |
|---|---|---|---|---|---|
| 40 | S100A1 | Core | -0.8367419 | 1.96E-38 | -13.129586 |
| 401 | S100A1 | Edge | 0.8367419 | 1.96E-38 | 13.129586 |
| 402 | TLR4 | Edge | -1.0201129 | 4.93E-38 | -13.057755 |
| 41 | TLR4 | Core | 1.0201129 | 4.93E-38 | 13.0577545 |
| 42 | NOTCH1 | Core | 0.78730184 | 6.98E-38 | 13.029484 |
| 403 | NOTCH1 | Edge | -0.7873018 | 6.98E-38 | -13.029484 |
| 43 | FGFR2 | Core | -0.9863306 | 2.02E-37 | -12.946403 |
| 404 | FGFR2 | Edge | 0.98633057 | 2.02E-37 | 12.946403 |
| 405 | CD68 | Edge | -1.32085 | 8.86E-36 | -12.651029 |
| 44 | CD68 | Core | 1.32085 | 8.86E-36 | 12.651029 |
| 45 | IFITM3 | Core | 1.234878 | 1.70E-34 | 12.415069 |
| 406 | IFITM3 | Edge | -1.234878 | 1.70E-34 | -12.415069 |
| 46 | RYR2 | Core | -2.7399166 | 1.18E-33 | -12.257027 |
| 407 | RYR2 | Edge | 2.7399166 | 1.18E-33 | 12.257027 |
| 408 | APP | Edge | 1.3495426 | 2.91E-33 | 12.182113 |
| 47 | APP | Core | -1.3495426 | 2.91E-33 | -12.182113 |
| 48 | HTR2A | Core | -2.905464 | 7.12E-33 | -12.107315 |
| 409 | HTR2A | Edge | 2.905464 | 7.12E-33 | 12.107315 |
| 410 | SPON1 | Edge | -0.7624061 | 7.98E-33 | -12.096217 |
| 49 | SPON1 | Core | 0.7624061 | 7.98E-33 | 12.096217 |
| 50 | FGFR3 | Core | -1.5099567 | 8.49E-33 | -12.089564 |
| 411 | FGFR3 | Edge | 1.5099567 | 8.49E-33 | 12.089564 |
| 412 | VCAM1 | Edge | -2.0522432 | 4.47E-31 | -11.757831 |
| 51 | VCAM1 | Core | 2.0522432 | 4.47E-31 | 11.757831 |
| 52 | THBS1 | Core | 2.9211092 | 1.73E-30 | 11.64138 |
| 413 | THBS1 | Edge | -2.9211092 | 1.73E-30 | -11.64138 |
| 414 | CCKBR | Edge | 2.767312 | 4.94E-29 | 11.350285 |
| 53 | CCKBR | Core | -2.767312 | 4.94E-29 | -11.350285 |
| 54 | DNER | Core | -0.9886161 | 2.63E-28 | -11.201631 |
| 415 | DNER | Edge | 0.98861605 | 2.63E-28 | 11.201631 |
| 416 | CD47 | Edge | 1.0501337 | 4.38E-28 | 11.154766 |
| 55 | CD47 | Core | -1.0501337 | 4.38E-28 | -11.154766 |
| 56 | ITGAX | Core | 1.3085576 | 6.55E-28 | 11.117287 |
| 417 | ITGAX | Edge | -1.3085576 | 6.55E-28 | -11.117287 |
| 418 | NPY1R | Edge | 2.0429878 | 3.29E-27 | 10.970579 |
| 57 | NPY1R | Core | -2.0429878 | 3.29E-27 | -10.970579 |
| 58 | KCNJ16 | Core | -1.3637148 | 6.19E-27 | -10.91188 |
| 419 | KCNJ16 | Edge | 1.3637148 | 6.19E-27 | 10.91188 |
| 420 | FSTL4 | Edge | 3.42851 | 2.78E-26 | 10.772864 |
| 59 | FSTL4 | Core | -3.42851 | 2.78E-26 | -10.772864 |
| 60 | CD48 | Core | 2.0249534 | 6.44E-26 | 10.693736 |

| | | | | | |
|---|---|---|---|---|---|
| 421 | CD48 | Edge | -2.0249534 | 6.44E-26 | -10.693736 |
| 422 | GJA1 | Edge | -1.007338 | 1.09E-25 | -10.64358 |
| 61 | GJA1 | Core | 1.007338 | 1.09E-25 | 10.6435795 |
| 423 | TGFB1 | Edge | -0.7475553 | 1.74E-25 | -10.598141 |
| 62 | TGFB1 | Core | 0.7475553 | 1.74E-25 | 10.598141 |
| 63 | GPR34 | Core | 0.9309423 | 3.43E-25 | 10.533056 |
| 424 | GPR34 | Edge | -0.9309423 | 3.43E-25 | -10.533056 |
| 64 | SLC17A6 | Core | -2.90195 | 3.67E-25 | -10.525159 |
| 425 | SLC17A6 | Edge | 2.90195 | 3.67E-25 | 10.525159 |
| 426 | KRAS | Edge | 0.98080075 | 8.81E-25 | 10.441009 |
| 65 | KRAS | Core | -0.9808008 | 8.81E-25 | -10.441009 |
| 66 | MEIS2 | Core | -0.7042193 | 1.99E-24 | -10.362126 |
| 427 | MEIS2 | Edge | 0.7042193 | 1.99E-24 | 10.362126 |
| 428 | CALCRL | Edge | -0.8696382 | 8.02E-24 | -10.226424 |
| 67 | CALCRL | Core | 0.8696382 | 8.02E-24 | 10.226424 |
| 429 | CDH12 | Edge | 2.2762415 | 2.02E-23 | 10.135103 |
| 68 | CDH12 | Core | -2.2762415 | 2.02E-23 | -10.135103 |
| 69 | PECAM1 | Core | 0.7945876 | 4.84E-23 | 10.047947 |
| 430 | PECAM1 | Edge | -0.7945876 | 4.84E-23 | -10.047947 |
| 70 | RYR3 | Core | -1.2393408 | 3.79E-22 | -9.841655 |
| 431 | RYR3 | Edge | 1.2393408 | 3.79E-22 | 9.841655 |
| 432 | RSPO2 | Edge | 2.0480564 | 1.01E-20 | 9.50404 |
| 71 | RSPO2 | Core | -2.0480564 | 1.01E-20 | -9.50404 |
| 433 | SETD2 | Edge | 0.589967 | 1.66E-20 | 9.451019 |
| 72 | SETD2 | Core | -0.589967 | 1.66E-20 | -9.451019 |
| 434 | HLA-DMB | Edge | -0.4474823 | 2.55E-20 | -9.404315 |
| 73 | HLA-DMB | Core | 0.44748226 | 2.55E-20 | 9.404315 |
| 74 | GAD2 | Core | -2.5776663 | 3.69E-20 | -9.364229 |
| 435 | GAD2 | Edge | 2.5776663 | 3.69E-20 | 9.364229 |
| 436 | MGP | Edge | -0.344115 | 8.28E-20 | 9.277017 |
| 75 | MGP | Core | 0.344115 | 8.28E-20 | -9.277017 |
| 76 | ITGB2 | Core | 0.9500034 | 1.92E-19 | 9.185686 |
| 437 | ITGB2 | Edge | -0.9500034 | 1.92E-19 | -9.185686 |
| 438 | COL4A1 | Edge | -0.8889483 | 2.91E-19 | -9.139127 |
| 77 | COL4A1 | Core | 0.8889483 | 2.91E-19 | 9.139127 |
| 78 | RELN | Core | -3.338587 | 1.40E-18 | -8.96639 |
| 439 | RELN | Edge | 3.338587 | 1.40E-18 | 8.96639 |
| 440 | ITGA5 | Edge | -1.4522016 | 1.83E-18 | -8.935272 |
| 79 | ITGA5 | Core | 1.4522016 | 1.83E-18 | 8.935272 |
| 441 | CXCR4 | Edge | -0.9445773 | 3.79E-18 | -8.853115 |
| 80 | CXCR4 | Core | 0.94457734 | 3.79E-18 | 8.853115 |

| | | | | | |
|---|---|---|---|---|---|
| 442 | GPR183 | Edge | -1.6608279 | 8.15E-18 | -8.766049 |
| 81 | GPR183 | Core | 1.6608279 | 8.15E-18 | 8.766049 |
| 443 | CAPG | Edge | -1.0352498 | 1.07E-17 | -8.734364 |
| 82 | CAPG | Core | 1.0352498 | 1.07E-17 | 8.734364 |
| 444 | CCL4 | Edge | -2.3848903 | 1.20E-17 | -8.7198105 |
| 83 | CCL4 | Core | 2.3848903 | 1.20E-17 | 8.7198105 |
| 445 | AXL | Edge | -0.7116143 | 1.33E-17 | -8.706337 |
| 84 | AXL | Core | 0.71161425 | 1.33E-17 | 8.706337 |
| 85 | ANO3 | Core | -1.6930943 | 1.63E-17 | -8.682306 |
| 446 | ANO3 | Edge | 1.6930943 | 1.63E-17 | 8.682306 |
| 447 | ACTA2 | Edge | 0.0213617 | 3.59E-17 | 8.59055 |
| 86 | ACTA2 | Core | -0.0213617 | 3.59E-17 | -8.59055 |
| 87 | ADRA1A | Core | -1.0587177 | 4.40E-17 | -8.56596 |
| 448 | ADRA1A | Edge | 1.0587177 | 4.40E-17 | 8.56596 |
| 449 | CD4 | Edge | -0.6475279 | 5.19E-17 | -8.545515 |
| 88 | CD4 | Core | 0.64752793 | 5.19E-17 | 8.545515 |
| 450 | RORB | Edge | 1.1646426 | 8.31E-17 | 8.489676 |
| 89 | RORB | Core | -1.1646426 | 8.31E-17 | -8.489676 |
| 90 | GAD1 | Core | -1.5157402 | 9.30E-17 | -8.475378 |
| 451 | GAD1 | Edge | 1.5157402 | 9.30E-17 | 8.475378 |
| 452 | PCNA | Edge | 0.603574 | 1.46E-16 | 8.421212 |
| 91 | PCNA | Core | -0.603574 | 1.46E-16 | -8.421212 |
| 92 | SULF1 | Core | 0.87242746 | 2.07E-16 | 8.379199 |
| 453 | SULF1 | Edge | -0.8724275 | 2.07E-16 | -8.379199 |
| 454 | CD44 | Edge | 0.4892051 | 2.49E-16 | -8.356016 |
| 93 | CD44 | Core | -0.4892051 | 2.49E-16 | 8.356016 |
| 94 | UGT8 | Core | 0.6605209 | 2.89E-16 | 8.337283 |
| 455 | UGT8 | Edge | -0.6605209 | 2.89E-16 | -8.337283 |
| 456 | CORO1A | Edge | 0.97617435 | 4.59E-16 | 8.28099 |
| 95 | CORO1A | Core | -0.9761744 | 4.59E-16 | -8.28099 |
| 96 | TGFBI | Core | 0.8787602 | 1.05E-15 | 8.181206 |
| 457 | TGFBI | Edge | -0.8787602 | 1.05E-15 | -8.181206 |
| 458 | ELOVL2 | Edge | 1.1783965 | 2.40E-15 | 8.079389 |
| 97 | ELOVL2 | Core | -1.1783965 | 2.40E-15 | -8.079389 |
| 98 | CD86 | Core | 0.81636155 | 2.51E-15 | 8.0727215 |
| 459 | CD86 | Edge | -0.8163616 | 2.51E-15 | -8.0727215 |
| 460 | FCGR1A | Edge | -0.6972515 | 6.16E-15 | -7.961051 |
| 99 | FCGR1A | Core | 0.6972515 | 6.16E-15 | 7.961051 |
| 100 | HLA-DQA1 | Core | 0.4900173 | 2.20E-14 | 7.8008385 |
| 461 | HLA-DQA1 | Edge | -0.4900173 | 2.20E-14 | -7.8008385 |
| 462 | FCGR3A | Edge | -0.5208198 | 4.25E-14 | -7.7158985 |

| | | | | | |
|---|---|---|---|---|---|
| 101 | FCGR3A | Core | 0.52081984 | 4.25E-14 | 7.7158985 |
| 102 | RFTN1 | Core | -0.7514907 | 1.43E-13 | -7.5582232 |
| 463 | RFTN1 | Edge | 0.75149065 | 1.43E-13 | 7.5582232 |
| 464 | PDGFD | Edge | 2.0154853 | 1.87E-13 | 7.522423 |
| 103 | PDGFD | Core | -2.0154853 | 1.87E-13 | -7.522423 |
| 465 | PTPRC | Edge | -0.5719973 | 3.78E-13 | -7.4282722 |
| 104 | PTPRC | Core | 0.57199734 | 3.78E-13 | 7.4282722 |
| 105 | HTRA1 | Core | -0.6863778 | 7.43E-13 | -7.3372045 |
| 466 | HTRA1 | Edge | 0.68637776 | 7.43E-13 | 7.3372045 |
| 467 | NR2F2 | Edge | 1.2875615 | 9.43E-13 | 7.30391 |
| 106 | NR2F2 | Core | -1.2875615 | 9.43E-13 | -7.30391 |
| 107 | PROX1 | Core | -0.7690858 | 2.23E-12 | -7.186275 |
| 468 | PROX1 | Edge | 0.7690858 | 2.23E-12 | 7.186275 |
| 469 | PAX6 | Edge | 0.62869215 | 3.20E-12 | 7.135024 |
| 108 | PAX6 | Core | -0.6286922 | 3.20E-12 | -7.135024 |
| 109 | CX3CR1 | Core | -0.7739572 | 2.60E-11 | -6.8398414 |
| 470 | CX3CR1 | Edge | 0.7739572 | 2.60E-11 | 6.8398414 |
| 471 | ADRA1B | Edge | 2.225001 | 3.24E-11 | 6.8069415 |
| 110 | ADRA1B | Core | -2.225001 | 3.24E-11 | -6.8069415 |
| 111 | SLIT3 | Core | -1.3484665 | 4.63E-11 | -6.7540555 |
| 472 | SLIT3 | Edge | 1.3484665 | 4.63E-11 | 6.7540555 |
| 473 | SPI1 | Edge | -0.7628487 | 1.14E-10 | -6.620956 |
| 112 | SPI1 | Core | 0.7628487 | 1.14E-10 | 6.620956 |
| 113 | SMARCB1 | Core | -0.3848002 | 3.47E-10 | -6.4532843 |
| 474 | SMARCB1 | Edge | 0.38480023 | 3.47E-10 | 6.4532843 |
| 114 | LY86 | Core | 0.6013808 | 1.33E-09 | 6.245367 |
| 475 | LY86 | Edge | -0.6013808 | 1.33E-09 | -6.245367 |
| 476 | TTYH1 | Edge | -0.303548 | 1.94E-09 | -6.184323 |
| 115 | TTYH1 | Core | 0.30354795 | 1.94E-09 | 6.184323 |
| 477 | OLIG2 | Edge | -1.0379643 | 3.49E-09 | -6.089788 |
| 116 | OLIG2 | Core | 1.0379643 | 3.49E-09 | 6.089788 |
| 117 | CNTN2 | Core | -0.4151978 | 5.39E-09 | -6.0185075 |
| 478 | CNTN2 | Edge | 0.41519776 | 5.39E-09 | 6.0185075 |
| 118 | TAGLN | Core | 1.2666669 | 7.09E-09 | 5.972457 |
| 479 | TAGLN | Edge | -1.2666669 | 7.09E-09 | -5.972457 |
| 119 | NES | Core | 0.3922096 | 9.37E-09 | 5.925511 |
| 480 | NES | Edge | -0.3922096 | 9.37E-09 | -5.925511 |
| 120 | PLD5 | Core | -1.0292279 | 2.77E-08 | -5.743344 |
| 481 | PLD5 | Edge | 1.0292279 | 2.77E-08 | 5.743344 |
| 482 | ENG | Edge | -0.3571298 | 3.27E-08 | -5.713957 |
| 121 | ENG | Core | 0.35712975 | 3.27E-08 | 5.713957 |

| | | | | | |
|---|---|---|---|---|---|
| 122 | LYPD6 | Core | -1.2269826 | 9.25E-08 | -5.5327263 |
| 483 | LYPD6 | Edge | 1.2269826 | 9.25E-08 | 5.5327263 |
| 484 | RYK | Edge | 0.41308558 | 9.46E-08 | 5.527442 |
| 123 | RYK | Core | -0.4130856 | 9.46E-08 | -5.527442 |
| 124 | HS3ST4 | Core | -1.2907977 | 1.74E-07 | -5.41817 |
| 485 | HS3ST4 | Edge | 1.2907977 | 1.74E-07 | 5.41817 |
| 486 | VIP | Edge | 1.7514553 | 2.41E-07 | 5.357732 |
| 125 | VIP | Core | -1.7514553 | 2.41E-07 | -5.357732 |
| 126 | C1orf162 | Core | 0.5010756 | 2.49E-07 | 5.3505287 |
| 487 | C1orf162 | Edge | -0.5010756 | 2.49E-07 | -5.3505287 |
| 127 | HILPDA | Core | -0.7668486 | 2.53E-07 | -5.3461456 |
| 488 | HILPDA | Edge | 0.76684856 | 2.53E-07 | 5.3461456 |
| 489 | RNASET2 | Edge | -0.5489979 | 5.56E-07 | -5.2006454 |
| 128 | RNASET2 | Core | 0.54899794 | 5.56E-07 | 5.2006454 |
| 129 | P2RY12 | Core | -0.4857907 | 8.35E-07 | -5.123109 |
| 490 | P2RY12 | Edge | 0.4857907 | 8.35E-07 | 5.123109 |
| 491 | TRHDE | Edge | 1.7924082 | 8.43E-07 | 5.1198363 |
| 130 | TRHDE | Core | -1.7924082 | 8.43E-07 | -5.1198363 |
| 131 | COL3A1 | Core | 1.0534093 | 9.28E-07 | 5.1002736 |
| 492 | COL3A1 | Edge | -1.0534093 | 9.28E-07 | -5.1002736 |
| 493 | RASGRP1 | Edge | 0.8986768 | 1.02E-06 | 5.0811124 |
| 132 | RASGRP1 | Core | -0.8986768 | 1.02E-06 | -5.0811124 |
| 494 | SOX9 | Edge | 0.09927601 | 1.39E-06 | 5.0196733 |
| 133 | SOX9 | Core | -0.099276 | 1.39E-06 | -5.0196733 |
| 134 | SNCG | Core | -1.5751036 | 1.83E-06 | -4.965324 |
| 495 | SNCG | Edge | 1.5751036 | 1.83E-06 | 4.965324 |
| 496 | HHATL | Edge | 0.34938246 | 2.50E-06 | 4.9032373 |
| 135 | HHATL | Core | -0.3493825 | 2.50E-06 | -4.9032373 |
| 136 | OPALIN | Core | -1.1457139 | 2.65E-06 | -4.8906198 |
| 497 | OPALIN | Edge | 1.1457139 | 2.65E-06 | 4.8906198 |
| 498 | CDH4 | Edge | 0.98419625 | 2.66E-06 | 4.8882484 |
| 137 | CDH4 | Core | -0.9841963 | 2.66E-06 | -4.8882484 |
| 138 | NWD2 | Core | -1.9504639 | 3.16E-06 | -4.8526363 |
| 499 | NWD2 | Edge | 1.9504639 | 3.16E-06 | 4.8526363 |
| 500 | RGS10 | Edge | -0.4189161 | 3.59E-06 | -4.8261995 |
| 139 | RGS10 | Core | 0.41891605 | 3.59E-06 | 4.8261995 |
| 140 | CNDP1 | Core | -0.8491208 | 4.12E-06 | -4.797102 |
| 501 | CNDP1 | Edge | 0.8491208 | 4.12E-06 | 4.797102 |
| 141 | CAMK2D | Core | -0.2870506 | 5.23E-06 | -4.7479086 |
| 502 | CAMK2D | Edge | 0.2870506 | 5.23E-06 | 4.7479086 |
| 503 | PLCH1 | Edge | 1.2308193 | 6.22E-06 | 4.711159 |

| | | | | | |
|---|---|---|---|---|---|
| 142 | PLCH1 | Core | -1.2308193 | 6.22E-06 | -4.711159 |
| 143 | NRP1 | Core | -0.6612922 | 7.45E-06 | -4.6727023 |
| 504 | NRP1 | Edge | 0.6612922 | 7.45E-06 | 4.6727023 |
| 505 | FCER1G | Edge | -0.2422527 | 9.47E-06 | -4.62183 |
| 144 | FCER1G | Core | 0.24225266 | 9.47E-06 | 4.62183 |
| 506 | TREM2 | Edge | -0.564815 | 9.96E-06 | -4.6099977 |
| 145 | TREM2 | Core | 0.564815 | 9.96E-06 | 4.6099977 |
| 146 | HIF1A | Core | 0.75164616 | 1.44E-05 | 4.5319157 |
| 507 | HIF1A | Edge | -0.7516462 | 1.44E-05 | -4.5319157 |
| 508 | MYL9 | Edge | -1.1024685 | 1.58E-05 | -4.5101123 |
| 147 | MYL9 | Core | 1.1024685 | 1.58E-05 | 4.5101123 |
| 148 | ITGA7 | Core | -0.3730696 | 1.66E-05 | -4.4977674 |
| 509 | ITGA7 | Edge | 0.3730696 | 1.66E-05 | 4.4977674 |
| 510 | MLC1 | Edge | -0.000715 | 2.29E-05 | -4.427736 |
| 149 | MLC1 | Core | 0.00071499 | 2.29E-05 | 4.427736 |
| 150 | OLIG1 | Core | 1.3164393 | 2.53E-05 | 4.4048014 |
| 511 | OLIG1 | Edge | -1.3164393 | 2.53E-05 | -4.4048014 |
| 512 | PCSK1 | Edge | 1.5903957 | 2.59E-05 | 4.3986406 |
| 151 | PCSK1 | Core | -1.5903957 | 2.59E-05 | -4.3986406 |
| 152 | TP53 | Core | 0.30038962 | 3.94E-05 | 4.304837 |
| 513 | TP53 | Edge | -0.3003896 | 3.94E-05 | -4.304837 |
| 514 | ADAMTS16 | Edge | 2.0280719 | 4.18E-05 | 4.290187 |
| 153 | ADAMTS16 | Core | -2.0280719 | 4.18E-05 | -4.290187 |
| 154 | BRINP3 | Core | -0.5101247 | 4.50E-05 | -4.272593 |
| 515 | BRINP3 | Edge | 0.51012474 | 4.50E-05 | 4.272593 |
| 516 | SLC24A3 | Edge | 0.71390736 | 4.89E-05 | 4.2527037 |
| 155 | SLC24A3 | Core | -0.7139074 | 4.89E-05 | -4.2527037 |
| 156 | PTPRZ1 | Core | 1.5068619 | 5.56E-05 | 4.2224092 |
| 517 | PTPRZ1 | Edge | -1.5068619 | 5.56E-05 | -4.2224092 |
| 518 | STXBP2 | Edge | -0.4912386 | 6.39E-05 | -4.189369 |
| 157 | STXBP2 | Core | 0.4912386 | 6.39E-05 | 4.189369 |
| 158 | KIT | Core | -1.3169664 | 7.51E-05 | -4.151191 |
| 519 | KIT | Edge | 1.3169664 | 7.51E-05 | 4.151191 |
| 520 | SORCS1 | Edge | 0.96391237 | 9.09E-05 | 4.1058908 |
| 159 | SORCS1 | Core | -0.9639124 | 9.09E-05 | -4.1058908 |
| 160 | S100A4 | Core | 0.9396413 | 0.00010237 | 4.076816 |
| 521 | S100A4 | Edge | -0.9396413 | 0.00010237 | -4.076816 |
| 522 | COL1A1 | Edge | -0.6336114 | 0.00011277 | -4.0528126 |
| 161 | COL1A1 | Core | 0.63361144 | 0.00011277 | 4.0528126 |
| 162 | TRPC5 | Core | -1.7321573 | 0.00011702 | -4.042715 |
| 523 | TRPC5 | Edge | 1.7321573 | 0.00011702 | 4.042715 |

| | | | | | |
|---|---|---|---|---|---|
| 524 | NTNG1 | Edge | 0.66023326 | 0.0001739 | 3.9474108 |
| 163 | NTNG1 | Core | -0.6602333 | 0.0001739 | -3.9474108 |
| 164 | PLCE1 | Core | 0.4075995 | 0.00018045 | 3.937084 |
| 525 | PLCE1 | Edge | -0.4075995 | 0.00018045 | -3.937084 |
| 526 | PTCH1 | Edge | 0.05030369 | 0.00027622 | 3.8321762 |
| 165 | PTCH1 | Core | -0.0503037 | 0.00027622 | -3.8321762 |
| 166 | CYTIP | Core | 0.7517921 | 0.00038218 | 3.7500544 |
| 527 | CYTIP | Edge | -0.7517921 | 0.00038218 | -3.7500544 |
| 528 | MAL | Edge | 0.2719683 | 0.00081764 | 3.5532455 |
| 167 | MAL | Core | -0.2719683 | 0.00081764 | -3.5532455 |
| 169 | PSEN1 | Core | -0.2064178 | 0.00093284 | -3.5152826 |
| 168 | CDK6 | Core | -0.2706671 | 0.00093284 | -3.5159483 |
| 530 | PSEN1 | Edge | 0.20641784 | 0.00093284 | 3.5152826 |
| 529 | CDK6 | Edge | 0.27066714 | 0.00093284 | 3.5159483 |

**Supplemental Table 7. Spatial transcriptomics derived differentially expressed genes (GBM)**

| | gene | region | logfoldchange | pval_adj | score |
|---|---|---|---|---|---|
| 0 | IGFBP5 | Core | 8.142127 | 0 | 113.22575 |
| 1 | VIM | Core | 7.2052455 | 0 | 96.49696 |
| 18 | IGFBP3 | Core | 2.1281564 | 0 | 45.772583 |
| 19 | ERMN | Core | -3.225273 | 0 | -42.956837 |
| 20 | OLIG2 | Core | -1.3107042 | 0 | -42.441578 |
| 21 | B4GALNT1 | Core | -3.110078 | 0 | -42.18719 |
| 17 | NCAM1 | Core | -1.7600701 | 0 | -46.563038 |
| 16 | GPR34 | Core | -1.7912499 | 0 | -46.81891 |
| 22 | MOBP | Core | -3.8789766 | 0 | -41.84925 |
| 23 | CD24 | Core | -1.9014045 | 0 | -40.27531 |
| 24 | IDH2 | Core | -1.0187265 | 0 | -39.447567 |
| 25 | P2RY12 | Core | -2.0119808 | 0 | -38.831863 |
| 26 | AIF1 | Core | -1.126893 | 0 | -38.34014 |
| 10 | THY1 | Core | -1.6694611 | 0 | -53.00779 |
| 3 | OLIG1 | Core | -2.7958424 | 0 | -80.35199 |
| 2 | APOE | Core | -5.6286345 | 0 | -91.51818 |
| 11 | DNER | Core | -1.4487908 | 0 | -51.877907 |
| 6 | HILPDA | Core | 3.3012857 | 0 | 59.727417 |
| 5 | POSTN | Core | 3.07778 | 0 | 60.236286 |
| 8 | CD44 | Core | 1.359423 | 0 | 57.744923 |
| 7 | BCAN | Core | -1.809346 | 0 | -59.7272 |
| 4 | ANXA1 | Core | 1.2733365 | 0 | 70.47087 |
| 366 | POSTN | Edge | -3.07778 | 0 | -60.236286 |
| 365 | ANXA1 | Edge | -1.2733365 | 0 | -70.47087 |
| 364 | OLIG1 | Edge | 2.7958424 | 0 | 80.35199 |
| 363 | APOE | Edge | 5.6286345 | 0 | 91.51818 |
| 362 | VIM | Edge | -7.2052455 | 0 | -96.49696 |
| 361 | IGFBP5 | Edge | -8.142127 | 0 | -113.22575 |
| 15 | CAV1 | Core | 1.8105588 | 0 | 47.01185 |
| 14 | MDM4 | Core | -2.905533 | 0 | -49.307125 |
| 13 | SOX4 | Core | -2.7725527 | 0 | -50.622673 |
| 12 | EFHD1 | Core | -1.734702 | 0 | -51.85827 |
| 368 | BCAN | Edge | 1.809346 | 0 | 59.7272 |
| 369 | CD44 | Edge | -1.359423 | 0 | -57.744923 |
| 367 | HILPDA | Edge | -3.3012857 | 0 | -59.727417 |
| 9 | TSPAN7 | Core | -2.1990712 | 0 | -54.543846 |
| 377 | GPR34 | Edge | 1.7912499 | 0 | 46.81891 |

| | | | | | |
|---|---|---|---|---|---|
| 378 | NCAM1 | Edge | 1.7600701 | 0 | 46.563038 |
| 379 | IGFBP3 | Edge | -2.1281564 | 0 | -45.772583 |
| 380 | ERMN | Edge | 3.225273 | 0 | 42.956837 |
| 381 | OLIG2 | Edge | 1.3107042 | 0 | 42.441578 |
| 382 | B4GALNT1 | Edge | 3.110078 | 0 | 42.18719 |
| 383 | MOBP | Edge | 3.8789766 | 0 | 41.84925 |
| 385 | IDH2 | Edge | 1.0187265 | 0 | 39.447567 |
| 384 | CD24 | Edge | 1.9014045 | 0 | 40.27531 |
| 370 | TSPAN7 | Edge | 2.1990712 | 0 | 54.543846 |
| 387 | AIF1 | Edge | 1.126893 | 0 | 38.34014 |
| 386 | P2RY12 | Edge | 2.0119808 | 0 | 38.831863 |
| 373 | EFHD1 | Edge | 1.734702 | 0 | 51.85827 |
| 374 | SOX4 | Edge | 2.7725527 | 0 | 50.622673 |
| 375 | MDM4 | Edge | 2.905533 | 0 | 49.307125 |
| 376 | CAV1 | Edge | -1.8105588 | 0 | -47.01185 |
| 371 | THY1 | Edge | 1.6694611 | 0 | 53.00779 |
| 372 | DNER | Edge | 1.4487908 | 0 | 51.877907 |
| 27 | LOX | Core | 3.5990028 | 9.96E-302 | 37.196663 |
| 388 | LOX | Edge | -3.5990028 | 9.96E-302 | -37.196663 |
| 389 | SOX11 | Edge | 2.4284177 | 9.36E-296 | 36.824398 |
| 28 | SOX11 | Core | -2.4284177 | 9.36E-296 | -36.824398 |
| 29 | UGT8 | Core | -1.6175233 | 1.12E-278 | -35.740578 |
| 390 | UGT8 | Edge | 1.6175233 | 1.12E-278 | 35.740578 |
| 391 | NES | Edge | 1.1254011 | 5.22E-257 | 34.31631 |
| 30 | NES | Core | -1.1254011 | 5.22E-257 | -34.31631 |
| 31 | NOTCH1 | Core | -1.0538911 | 1.83E-240 | -33.187717 |
| 392 | NOTCH1 | Edge | 1.0538911 | 1.83E-240 | 33.187717 |
| 393 | BMI1 | Edge | 0.88420254 | 9.03E-238 | 32.999504 |
| 32 | BMI1 | Core | -0.8842025 | 9.03E-238 | -32.999504 |
| 33 | AQP4 | Core | -4.4814067 | 1.66E-235 | -32.84037 |
| 394 | AQP4 | Edge | 4.4814067 | 1.66E-235 | 32.84037 |
| 34 | CX3CR1 | Core | -1.7290965 | 1.30E-210 | -31.046827 |
| 395 | CX3CR1 | Edge | 1.7290965 | 1.30E-210 | 31.046827 |
| 35 | FCGR1A | Core | -1.2025713 | 4.70E-208 | -30.855753 |
| 396 | FCGR1A | Edge | 1.2025713 | 4.70E-208 | 30.855753 |
| 36 | VCP | Core | -0.670753 | 2.99E-194 | -29.808119 |
| 397 | VCP | Edge | 0.670753 | 2.99E-194 | 29.808119 |
| 37 | MS4A6A | Core | -1.1577812 | 9.40E-182 | -28.82681 |
| 398 | MS4A6A | Edge | 1.1577812 | 9.40E-182 | 28.82681 |

| | | | | | |
|---|---|---|---|---|---|
| 399 | TMIGD3 | Edge | 1.2554789 | 2.20E-173 | 28.150278 |
| 38 | TMIGD3 | Core | -1.2554789 | 2.20E-173 | -28.150278 |
| 400 | KRAS | Edge | 0.7946333 | 5.39E-162 | 27.203096 |
| 39 | KRAS | Core | -0.7946333 | 5.39E-162 | -27.203096 |
| 40 | CALCRL | Core | -1.177118 | 1.26E-161 | -27.170973 |
| 401 | CALCRL | Edge | 1.177118 | 1.26E-161 | 27.170973 |
| 402 | CNTNAP3B | Edge | -0.9647464 | 3.54E-157 | -26.790987 |
| 41 | CNTNAP3B | Core | 0.96474636 | 3.54E-157 | 26.790987 |
| 42 | TTYH1 | Core | -0.7093551 | 2.45E-151 | -26.284046 |
| 403 | TTYH1 | Edge | 0.70935506 | 2.45E-151 | 26.284046 |
| 43 | CXCL14 | Core | 0.49505427 | 1.04E-150 | 26.228088 |
| 404 | CXCL14 | Edge | -0.4950543 | 1.04E-150 | -26.228088 |
| 405 | CCND2 | Edge | 1.3833995 | 1.60E-150 | 26.210873 |
| 44 | CCND2 | Core | -1.3833995 | 1.60E-150 | -26.210873 |
| 45 | NNAT | Core | -1.6509762 | 1.93E-144 | -25.671162 |
| 406 | NNAT | Edge | 1.6509762 | 1.93E-144 | 25.671162 |
| 46 | RGS5 | Core | -0.8502717 | 2.75E-144 | -25.656404 |
| 407 | RGS5 | Edge | 0.8502717 | 2.75E-144 | 25.656404 |
| 408 | HTRA1 | Edge | 1.5256681 | 6.09E-144 | 25.624714 |
| 47 | HTRA1 | Core | -1.5256681 | 6.09E-144 | -25.624714 |
| 48 | HLA-DMB | Core | -1.0291119 | 2.96E-137 | -25.016806 |
| 409 | HLA-DMB | Edge | 1.0291119 | 2.96E-137 | 25.016806 |
| 410 | PTCH1 | Edge | 1.0099678 | 1.94E-130 | 24.381416 |
| 49 | PTCH1 | Core | -1.0099678 | 1.94E-130 | -24.381416 |
| 50 | CCND1 | Core | -0.9317139 | 1.26E-128 | -24.209322 |
| 411 | CCND1 | Edge | 0.9317139 | 1.26E-128 | 24.209322 |
| 412 | SLC17A7 | Edge | 4.0002875 | 1.65E-123 | 23.717442 |
| 51 | SLC17A7 | Core | -4.0002875 | 1.65E-123 | -23.717442 |
| 52 | FCER1G | Core | -0.5600136 | 2.10E-119 | -23.31552 |
| 413 | FCER1G | Edge | 0.5600136 | 2.10E-119 | 23.31552 |
| 414 | RGS10 | Edge | 0.9976627 | 1.46E-118 | 23.231646 |
| 53 | RGS10 | Core | -0.9976627 | 1.46E-118 | -23.231646 |
| 54 | RNASET2 | Core | -0.7655471 | 1.06E-116 | -23.045973 |
| 415 | RNASET2 | Edge | 0.7655471 | 1.06E-116 | 23.045973 |
| 416 | SETD2 | Edge | 0.625222 | 2.53E-116 | 23.007465 |
| 55 | SETD2 | Core | -0.625222 | 2.53E-116 | -23.007465 |
| 56 | PECAM1 | Core | -0.8359903 | 8.94E-116 | -22.95186 |
| 417 | PECAM1 | Edge | 0.8359903 | 8.94E-116 | 22.95186 |
| 418 | CNDP1 | Edge | 3.0716755 | 8.39E-115 | 22.853516 |

| | | | | | |
|---|---|---|---|---|---|
| 57 | CNDP1 | Core | -3.0716755 | 8.39E-115 | -22.853516 |
| 58 | CAMK2A | Core | -1.9096402 | 3.98E-114 | -22.784632 |
| 419 | CAMK2A | Edge | 1.9096402 | 3.98E-114 | 22.784632 |
| 420 | AXL | Edge | 0.8016638 | 6.80E-113 | 22.659246 |
| 59 | AXL | Core | -0.8016638 | 6.80E-113 | -22.659246 |
| 60 | TRIL | Core | -0.7769944 | 3.45E-111 | -22.484877 |
| 421 | TRIL | Edge | 0.7769944 | 3.45E-111 | 22.484877 |
| 422 | GPR37L1 | Edge | 0.79399985 | 3.62E-109 | 22.276672 |
| 61 | GPR37L1 | Core | -0.7939999 | 3.62E-109 | -22.276672 |
| 423 | ADRA1A | Edge | 0.93117076 | 1.13E-107 | 22.121458 |
| 62 | ADRA1A | Core | -0.9311708 | 1.13E-107 | -22.121458 |
| 63 | CTSS | Core | -0.8623501 | 5.17E-103 | -21.63108 |
| 424 | CTSS | Edge | 0.8623501 | 5.17E-103 | 21.63108 |
| 64 | TLR4 | Core | -0.8316662 | 1.00E-102 | -21.59982 |
| 425 | TLR4 | Edge | 0.8316662 | 1.00E-102 | 21.59982 |
| 426 | TREM2 | Edge | 1.3511374 | 3.59E-100 | 21.325674 |
| 65 | TREM2 | Core | -1.3511374 | 3.59E-100 | -21.325674 |
| 66 | PDGFRA | Core | -0.9802988 | 1.99E-93 | -20.585724 |
| 427 | PDGFRA | Edge | 0.98029876 | 1.99E-93 | 20.585724 |
| 428 | MOG | Edge | 2.5841858 | 1.35E-92 | 20.492039 |
| 67 | MOG | Core | -2.5841858 | 1.35E-92 | -20.492039 |
| 429 | PBRM1 | Edge | 0.70211625 | 1.38E-92 | 20.49026 |
| 68 | PBRM1 | Core | -0.7021163 | 1.38E-92 | -20.49026 |
| 69 | CNTN2 | Core | -2.0128617 | 1.53E-90 | -20.25892 |
| 430 | CNTN2 | Edge | 2.0128617 | 1.53E-90 | 20.25892 |
| 70 | KCNJ16 | Core | -0.5646947 | 8.87E-90 | -20.171476 |
| 431 | KCNJ16 | Edge | 0.5646947 | 8.87E-90 | 20.171476 |
| 432 | ST18 | Edge | 2.1599655 | 7.49E-89 | 20.06494 |
| 71 | ST18 | Core | -2.1599655 | 7.49E-89 | -20.06494 |
| 433 | SLC24A3 | Edge | 1.1040711 | 8.90E-88 | 19.940847 |
| 72 | SLC24A3 | Core | -1.1040711 | 8.90E-88 | -19.940847 |
| 434 | SOX10 | Edge | 0.8894636 | 1.88E-85 | 19.670528 |
| 73 | SOX10 | Core | -0.8894636 | 1.88E-85 | -19.670528 |
| 74 | HLA-DQA1 | Core | -1.1687738 | 6.96E-85 | -19.603405 |
| 435 | HLA-DQA1 | Edge | 1.1687738 | 6.96E-85 | 19.603405 |
| 436 | HEPACAM | Edge | 0.9336214 | 2.33E-83 | 19.423382 |
| 75 | HEPACAM | Core | -0.9336214 | 2.33E-83 | -19.423382 |
| 76 | TGFB2 | Core | 0.5664203 | 1.38E-81 | 19.211994 |
| 437 | TGFB2 | Edge | -0.5664203 | 1.38E-81 | -19.211994 |

| | | | | | |
|---|---|---|---|---|---|
| 438 | FCGR3A | Edge | 0.69955885 | 1.34E-79 | 18.972193 |
| 77 | FCGR3A | Core | -0.6995589 | 1.34E-79 | -18.972193 |
| 78 | ITGB2 | Core | -0.7771766 | 6.62E-77 | -18.642662 |
| 439 | ITGB2 | Edge | 0.77717656 | 6.62E-77 | 18.642662 |
| 440 | CLDN11 | Edge | 0.7106517 | 7.57E-77 | 18.634823 |
| 79 | CLDN11 | Core | -0.7106517 | 7.57E-77 | -18.634823 |
| 441 | CENPF | Edge | 0.8900023 | 1.27E-76 | 18.606588 |
| 80 | CENPF | Core | -0.8900023 | 1.27E-76 | -18.606588 |
| 442 | FLT1 | Edge | 0.98552674 | 4.83E-76 | 18.534033 |
| 81 | FLT1 | Core | -0.9855267 | 4.83E-76 | -18.534033 |
| 443 | LY86 | Edge | 1.153127 | 2.10E-74 | 18.329393 |
| 82 | LY86 | Core | -1.153127 | 2.10E-74 | -18.329393 |
| 444 | S100A1 | Edge | 0.55641556 | 3.43E-73 | 18.176058 |
| 83 | S100A1 | Core | -0.5564156 | 3.43E-73 | -18.176058 |
| 445 | HHIP | Edge | 1.7020441 | 4.60E-71 | 17.904715 |
| 84 | HHIP | Core | -1.7020441 | 4.60E-71 | -17.904715 |
| 85 | TGFBI | Core | 1.0064948 | 2.73E-70 | 17.804586 |
| 446 | TGFBI | Edge | -1.0064948 | 2.73E-70 | -17.804586 |
| 447 | ARHGAP24 | Edge | 0.96229446 | 6.22E-70 | 17.757826 |
| 86 | ARHGAP24 | Core | -0.9622945 | 6.22E-70 | -17.757826 |
| 87 | GAD1 | Core | -1.6688731 | 1.09E-65 | -17.199818 |
| 448 | GAD1 | Edge | 1.6688731 | 1.09E-65 | 17.199818 |
| 449 | CCK | Edge | 4.0778713 | 2.25E-65 | 17.157219 |
| 88 | CCK | Core | -4.0778713 | 2.25E-65 | -17.157219 |
| 450 | TP53 | Edge | 0.47462988 | 1.10E-64 | 17.063917 |
| 89 | TP53 | Core | -0.4746299 | 1.10E-64 | -17.063917 |
| 90 | HIF1A | Core | -1.2199996 | 1.87E-64 | -17.032635 |
| 451 | HIF1A | Edge | 1.2199996 | 1.87E-64 | 17.032635 |
| 452 | CAPN3 | Edge | 2.1252468 | 9.25E-64 | 16.938042 |
| 91 | CAPN3 | Core | -2.1252468 | 9.25E-64 | -16.938042 |
| 92 | CTSH | Core | -0.7593692 | 2.80E-62 | -16.735674 |
| 453 | CTSH | Edge | 0.7593692 | 2.80E-62 | 16.735674 |
| 454 | MFSD2A | Edge | 1.0099792 | 1.10E-61 | 16.653305 |
| 93 | MFSD2A | Core | -1.0099792 | 1.10E-61 | -16.653305 |
| 94 | FBLN1 | Core | -0.7845321 | 3.04E-61 | -16.591797 |
| 455 | FBLN1 | Edge | 0.7845321 | 3.04E-61 | 16.591797 |
| 456 | CAMK2D | Edge | -0.2335725 | 1.47E-59 | -16.35653 |
| 95 | CAMK2D | Core | 0.23357254 | 1.47E-59 | 16.35653 |
| 96 | ESAM | Core | -1.0382978 | 5.24E-59 | -16.27827 |

| | | | | | |
|---|---|---|---|---|---|
| 457 | ESAM | Edge | 1.0382978 | 5.24E-59 | 16.27827 |
| 458 | FREM2 | Edge | -0.3446066 | 5.90E-59 | -16.270409 |
| 97 | FREM2 | Core | 0.34460655 | 5.90E-59 | 16.270409 |
| 98 | RORB | Core | 0.3927236 | 1.80E-57 | 16.059248 |
| 459 | RORB | Edge | -0.3927236 | 1.80E-57 | -16.059248 |
| 460 | PROX1 | Edge | 0.7011614 | 3.11E-57 | 16.024612 |
| 99 | PROX1 | Core | -0.7011614 | 3.11E-57 | -16.024612 |
| 100 | CORO1A | Core | -1.2190922 | 7.33E-57 | -15.970527 |
| 461 | CORO1A | Edge | 1.2190922 | 7.33E-57 | 15.970527 |
| 462 | MYO16 | Edge | 0.79083806 | 7.78E-57 | 15.966158 |
| 101 | MYO16 | Core | -0.7908381 | 7.78E-57 | -15.966158 |
| 102 | EZH2 | Core | -0.5674951 | 2.54E-56 | -15.891558 |
| 463 | EZH2 | Edge | 0.5674951 | 2.54E-56 | 15.891558 |
| 464 | SMARCB1 | Edge | 0.50786847 | 6.77E-56 | 15.82939 |
| 103 | SMARCB1 | Core | -0.5078685 | 6.77E-56 | -15.82939 |
| 465 | TOP2A | Edge | 0.8387834 | 8.02E-56 | 15.818173 |
| 104 | TOP2A | Core | -0.8387834 | 8.02E-56 | -15.818173 |
| 105 | EGFR | Core | 0.7248239 | 4.21E-55 | 15.7127905 |
| 466 | EGFR | Edge | -0.7248239 | 4.21E-55 | -15.712791 |
| 467 | IFITM3 | Edge | -0.5815972 | 8.57E-55 | -15.667089 |
| 106 | IFITM3 | Core | 0.5815972 | 8.57E-55 | 15.667089 |
| 107 | FGF13 | Core | -1.5746655 | 8.66E-55 | -15.66581 |
| 468 | FGF13 | Edge | 1.5746655 | 8.66E-55 | 15.66581 |
| 469 | PTPRC | Edge | 0.58207506 | 1.13E-53 | 15.500936 |
| 108 | PTPRC | Core | -0.5820751 | 1.13E-53 | -15.500936 |
| 109 | KDR | Core | -0.9700269 | 2.19E-53 | -15.45796 |
| 470 | KDR | Edge | 0.97002685 | 2.19E-53 | 15.45796 |
| 471 | PCNA | Edge | 0.54617107 | 1.00E-52 | 15.3589325 |
| 110 | PCNA | Core | -0.5461711 | 1.00E-52 | -15.358933 |
| 111 | COL4A1 | Core | 0.08849215 | 2.25E-52 | 15.305848 |
| 472 | COL4A1 | Edge | -0.0884922 | 2.25E-52 | -15.305848 |
| 473 | KIT | Edge | 1.7937459 | 7.06E-50 | 14.926718 |
| 112 | KIT | Core | -1.7937459 | 7.06E-50 | -14.926718 |
| 113 | C1orf162 | Core | -0.657405 | 2.06E-49 | -14.854483 |
| 474 | C1orf162 | Edge | 0.65740496 | 2.06E-49 | 14.854483 |
| 114 | MYRF | Core | -1.3523389 | 6.54E-46 | -14.303439 |
| 475 | MYRF | Edge | 1.3523389 | 6.54E-46 | 14.303439 |
| 476 | MLC1 | Edge | 0.6657969 | 3.25E-44 | 14.02846 |
| 115 | MLC1 | Core | -0.6657969 | 3.25E-44 | -14.02846 |

| | | | | | |
|---|---|---|---|---|---|
| 477 | SULF1 | Edge | 0.3974371 | 6.60E-44 | 13.977459 |
| 116 | SULF1 | Core | -0.3974371 | 6.60E-44 | -13.977459 |
| 117 | CD68 | Core | -0.2940339 | 6.40E-43 | -13.814205 |
| 478 | CD68 | Edge | 0.2940339 | 6.40E-43 | 13.814205 |
| 118 | ERBB3 | Core | -0.5139954 | 1.29E-42 | -13.762952 |
| 479 | ERBB3 | Edge | 0.51399535 | 1.29E-42 | 13.762952 |
| 119 | BRINP3 | Core | -0.5543683 | 2.09E-41 | -13.559497 |
| 480 | BRINP3 | Edge | 0.5543683 | 2.09E-41 | 13.559497 |
| 120 | MSH6 | Core | -0.4559372 | 1.47E-40 | -13.415174 |
| 481 | MSH6 | Edge | 0.45593715 | 1.47E-40 | 13.415174 |
| 482 | THBS1 | Edge | -0.6325565 | 1.16E-39 | -13.260829 |
| 121 | THBS1 | Core | 0.6325565 | 1.16E-39 | 13.260829 |
| 122 | ITGA7 | Core | 0.26823047 | 3.30E-39 | 13.181401 |
| 483 | ITGA7 | Edge | -0.2682305 | 3.30E-39 | -13.181401 |
| 484 | SPI1 | Edge | 0.76336294 | 4.84E-38 | 12.976502 |
| 123 | SPI1 | Core | -0.7633629 | 4.84E-38 | -12.976502 |
| 124 | CDK6 | Core | 0.26247576 | 1.06E-36 | 12.737315 |
| 485 | CDK6 | Edge | -0.2624758 | 1.06E-36 | -12.737315 |
| 486 | MAL | Edge | 1.5833836 | 1.95E-36 | 12.689029 |
| 125 | MAL | Core | -1.5833836 | 1.95E-36 | -12.689029 |
| 126 | MAG | Core | -2.0040274 | 4.66E-36 | -12.619972 |
| 487 | MAG | Edge | 2.0040274 | 4.66E-36 | 12.619972 |
| 127 | CUX2 | Core | -2.5708537 | 2.07E-35 | -12.501279 |
| 488 | CUX2 | Edge | 2.5708537 | 2.07E-35 | 12.501279 |
| 489 | ENG | Edge | 0.5641973 | 2.14E-35 | 12.498208 |
| 128 | ENG | Core | -0.5641973 | 2.14E-35 | -12.498208 |
| 129 | TAC1 | Core | -3.1045418 | 2.14E-35 | -12.497367 |
| 490 | TAC1 | Edge | 3.1045418 | 2.14E-35 | 12.497367 |
| 491 | LAMA2 | Edge | -0.2406216 | 4.72E-35 | -12.433794 |
| 130 | LAMA2 | Core | 0.24062164 | 4.72E-35 | 12.433794 |
| 131 | HS3ST2 | Core | -1.7752228 | 5.95E-34 | -12.228951 |
| 492 | HS3ST2 | Edge | 1.7752228 | 5.95E-34 | 12.228951 |
| 493 | ITGAX | Edge | 0.63041216 | 1.49E-33 | 12.153397 |
| 132 | ITGAX | Core | -0.6304122 | 1.49E-33 | -12.153397 |
| 494 | PCSK6 | Edge | 1.1320989 | 2.48E-33 | 12.111137 |
| 133 | PCSK6 | Core | -1.1320989 | 2.48E-33 | -12.111137 |
| 134 | CD4 | Core | -0.5027005 | 9.89E-33 | -11.996678 |
| 495 | CD4 | Edge | 0.5027005 | 9.89E-33 | 11.996678 |
| 496 | TPM2 | Edge | -0.4187916 | 1.51E-31 | -11.768204 |

| | | | | | |
|---|---|---|---|---|---|
| 135 | TPM2 | Core | 0.41879162 | 1.51E-31 | 11.768204 |
| 136 | SCN8A | Core | -0.8141044 | 1.96E-31 | -11.745811 |
| 497 | SCN8A | Edge | 0.81410444 | 1.96E-31 | 11.7458105 |
| 498 | ANGPT1 | Edge | 0.6580143 | 3.67E-31 | 11.691808 |
| 137 | ANGPT1 | Core | -0.6580143 | 3.67E-31 | -11.691808 |
| 138 | IGFBP4 | Core | -0.2684139 | 8.34E-31 | -11.621284 |
| 499 | IGFBP4 | Edge | 0.2684139 | 8.34E-31 | 11.621284 |
| 500 | COL1A1 | Edge | -1.8003541 | 5.20E-30 | -11.463239 |
| 139 | COL1A1 | Core | 1.8003541 | 5.20E-30 | 11.463239 |
| 140 | P2RY13 | Core | -0.9542248 | 5.36E-29 | -11.258925 |
| 501 | P2RY13 | Edge | 0.9542248 | 5.36E-29 | 11.258925 |
| 141 | NT5E | Core | 0.5793617 | 7.61E-29 | 11.227358 |
| 502 | NT5E | Edge | -0.5793617 | 7.61E-29 | -11.227358 |
| 503 | NTNG2 | Edge | 0.76785743 | 2.49E-28 | 11.121569 |
| 142 | NTNG2 | Core | -0.7678574 | 2.49E-28 | -11.121569 |
| 143 | RIT2 | Core | -1.001066 | 5.02E-27 | -10.849683 |
| 504 | RIT2 | Edge | 1.001066 | 5.02E-27 | 10.849683 |
| 505 | FGFR2 | Edge | 0.49644396 | 1.40E-26 | 10.754613 |
| 144 | FGFR2 | Core | -0.496444 | 1.40E-26 | -10.754613 |
| 506 | MKI67 | Edge | 0.64799404 | 3.25E-26 | 10.676359 |
| 145 | MKI67 | Core | -0.647994 | 3.25E-26 | -10.676359 |
| 146 | EYA4 | Core | 0.5806329 | 5.60E-26 | 10.624946 |
| 507 | EYA4 | Edge | -0.5806329 | 5.60E-26 | -10.624946 |
| 508 | SAMD5 | Edge | 0.96929955 | 1.73E-25 | 10.518382 |
| 147 | SAMD5 | Core | -0.9692996 | 1.73E-25 | -10.518382 |
| 148 | ITGA5 | Core | 0.41104677 | 8.46E-25 | 10.367283 |
| 509 | ITGA5 | Edge | -0.4110468 | 8.46E-25 | -10.367283 |
| 510 | SPON1 | Edge | 0.6522031 | 4.28E-24 | 10.210567 |
| 149 | SPON1 | Core | -0.6522031 | 4.28E-24 | -10.210567 |
| 150 | CD86 | Core | -0.5872008 | 1.09E-23 | -10.118986 |
| 511 | CD86 | Edge | 0.5872008 | 1.09E-23 | 10.118986 |
| 512 | SERPINA3 | Edge | 0.91690826 | 3.29E-23 | 10.009572 |
| 151 | SERPINA3 | Core | -0.9169083 | 3.29E-23 | -10.009572 |
| 152 | THSD4 | Core | -0.9353885 | 3.99E-23 | -9.989741 |
| 513 | THSD4 | Edge | 0.93538845 | 3.99E-23 | 9.989741 |
| 514 | STK32B | Edge | -0.6620354 | 7.53E-23 | -9.925897 |
| 153 | STK32B | Core | 0.66203535 | 7.53E-23 | 9.925897 |
| 154 | PTPRZ1 | Core | -2.6455255 | 1.50E-22 | -9.855936 |
| 515 | PTPRZ1 | Edge | 2.6455255 | 1.50E-22 | 9.855936 |

| | | | | | |
|---|---|---|---|---|---|
| 516 | PTPN12 | Edge | -0.2460427 | 1.59E-22 | -9.850034 |
| 155 | PTPN12 | Core | 0.24604271 | 1.59E-22 | 9.850034 |
| 156 | PAX6 | Core | -0.3718905 | 5.71E-22 | -9.719816 |
| 517 | PAX6 | Edge | 0.37189054 | 5.71E-22 | 9.719816 |
| 518 | HHATL | Edge | 1.5008154 | 1.46E-21 | 9.6229105 |
| 157 | HHATL | Core | -1.5008154 | 1.46E-21 | -9.6229105 |
| 158 | CD83 | Core | -0.5741904 | 3.26E-21 | -9.539573 |
| 519 | CD83 | Edge | 0.5741904 | 3.26E-21 | 9.539573 |
| 520 | VCAM1 | Edge | -0.4843888 | 9.95E-21 | -9.422403 |
| 159 | VCAM1 | Core | 0.48438877 | 9.95E-21 | 9.422403 |
| 160 | NTNG1 | Core | -0.4371251 | 5.80E-20 | -9.234904 |
| 521 | NTNG1 | Edge | 0.43712512 | 5.80E-20 | 9.234904 |
| 522 | ITGA8 | Edge | 1.3441229 | 1.40E-19 | 9.139393 |
| 161 | ITGA8 | Core | -1.3441229 | 1.40E-19 | -9.139393 |
| 162 | FGFR3 | Core | -0.4237613 | 2.29E-19 | -9.085326 |
| 523 | FGFR3 | Edge | 0.42376128 | 2.29E-19 | 9.085326 |
| 524 | NRP1 | Edge | -0.2603067 | 4.05E-19 | -9.022424 |
| 163 | NRP1 | Core | 0.26030672 | 4.05E-19 | 9.022424 |
| 164 | TGFB1 | Core | -0.2302094 | 7.74E-19 | -8.950525 |
| 525 | TGFB1 | Edge | 0.23020938 | 7.74E-19 | 8.950525 |
| 526 | SOX2 | Edge | 0.427594 | 9.24E-19 | 8.930287 |
| 165 | SOX2 | Core | -0.427594 | 9.24E-19 | -8.930287 |
| 166 | GAD2 | Core | -3.290552 | 8.99E-18 | -8.674305 |
| 527 | GAD2 | Edge | 3.290552 | 8.99E-18 | 8.674305 |
| 528 | TACR1 | Edge | 1.3916978 | 1.38E-17 | 8.624543 |
| 167 | TACR1 | Core | -1.3916978 | 1.38E-17 | -8.624543 |
| 168 | C1QL3 | Core | -2.3548565 | 1.88E-17 | -8.588631 |
| 529 | C1QL3 | Edge | 2.3548565 | 1.88E-17 | 8.588631 |
| 530 | TAGLN | Edge | -0.6910105 | 2.58E-17 | -8.551339 |
| 169 | TAGLN | Core | 0.6910105 | 2.58E-17 | 8.551339 |
| 170 | HTR2A | Core | -1.4024544 | 1.99E-16 | -8.311839 |
| 531 | HTR2A | Edge | 1.4024544 | 1.99E-16 | 8.311839 |
| 532 | SLC17A6 | Edge | 1.1314877 | 2.70E-16 | 8.274629 |
| 171 | SLC17A6 | Core | -1.1314877 | 2.70E-16 | -8.274629 |
| 172 | OPALIN | Core | -1.670714 | 2.96E-16 | -8.263319 |
| 533 | OPALIN | Edge | 1.670714 | 2.96E-16 | 8.263319 |
| 534 | SYNPR | Edge | 1.2404021 | 7.42E-16 | 8.152153 |
| 173 | SYNPR | Core | -1.2404021 | 7.42E-16 | -8.152153 |
| 174 | PDGFRB | Core | -0.4186812 | 1.03E-15 | -8.111138 |

| | | | | | |
|---|---|---|---|---|---|
| 535 | PDGFRB | Edge | 0.41868117 | 1.03E-15 | 8.111138 |
| 536 | MCTP2 | Edge | -1.0899403 | 1.85E-15 | -8.039211 |
| 175 | MCTP2 | Core | 1.0899403 | 1.85E-15 | 8.039211 |
| 176 | NPNT | Core | 0.21324582 | 4.33E-15 | 7.9338417 |
| 537 | NPNT | Edge | -0.2132458 | 4.33E-15 | -7.9338417 |
| 538 | GJA1 | Edge | 0.95623374 | 5.61E-15 | -7.900916 |
| 177 | GJA1 | Core | -0.9562337 | 5.61E-15 | 7.900916 |
| 178 | VWC2L | Core | -1.0766319 | 2.03E-14 | -7.738492 |
| 539 | VWC2L | Edge | 1.0766319 | 2.03E-14 | 7.738492 |
| 179 | MYCN | Core | -0.7324212 | 3.49E-14 | -7.6685348 |
| 540 | MYCN | Edge | 0.73242116 | 3.49E-14 | 7.6685348 |
| 541 | GPNMB | Edge | 1.2551246 | 3.65E-14 | 7.6620584 |
| 180 | GPNMB | Core | -1.2551246 | 3.65E-14 | -7.6620584 |
| 181 | PLCH1 | Core | -0.8310558 | 2.45E-13 | -7.4129143 |
| 542 | PLCH1 | Edge | 0.83105576 | 2.45E-13 | 7.4129143 |
| 543 | ELOVL2 | Edge | 0.20103104 | 4.18E-13 | -7.340971 |
| 182 | ELOVL2 | Core | -0.201031 | 4.18E-13 | 7.340971 |
| 183 | CEMIP2 | Core | -0.3179736 | 7.12E-13 | -7.268735 |
| 544 | CEMIP2 | Edge | 0.3179736 | 7.12E-13 | 7.268735 |
| 545 | RYR1 | Edge | 0.8692817 | 7.47E-13 | 7.2614956 |
| 184 | RYR1 | Core | -0.8692817 | 7.47E-13 | -7.2614956 |
| 185 | RYK | Core | -0.2445118 | 1.91E-12 | -7.133051 |
| 546 | RYK | Edge | 0.24451178 | 1.91E-12 | 7.133051 |
| 547 | CD163 | Edge | 0.24682322 | 1.92E-12 | 7.1312933 |
| 186 | CD163 | Core | -0.2468232 | 1.92E-12 | -7.1312933 |
| 548 | MYO5B | Edge | -0.3978985 | 4.10E-12 | -7.0252857 |
| 187 | MYO5B | Core | 0.39789852 | 4.10E-12 | 7.0252857 |
| 188 | COL3A1 | Core | 1.5891498 | 9.89E-12 | 6.900589 |
| 549 | COL3A1 | Edge | -1.5891498 | 9.89E-12 | -6.900589 |
| 189 | ITGAM | Core | -0.3800021 | 1.19E-11 | -6.873495 |
| 550 | ITGAM | Edge | 0.3800021 | 1.19E-11 | 6.873495 |
| 551 | CEMIP | Edge | -0.5342739 | 1.55E-11 | -6.834973 |
| 190 | CEMIP | Core | 0.5342739 | 1.55E-11 | 6.834973 |
| 552 | PSEN1 | Edge | 0.26131484 | 2.71E-11 | 6.7536716 |
| 191 | PSEN1 | Core | -0.2613148 | 2.71E-11 | -6.7536716 |
| 192 | LAMP5 | Core | -0.6713654 | 2.74E-11 | -6.7515635 |
| 553 | LAMP5 | Edge | 0.6713654 | 2.74E-11 | 6.7515635 |
| 554 | CSPG4 | Edge | 0.5353543 | 3.58E-11 | 6.7117715 |
| 193 | CSPG4 | Core | -0.5353543 | 3.58E-11 | -6.7117715 |

| | | | | | |
|---|---|---|---|---|---|
| 194 | CAPG | Core | -0.2384242 | 5.50E-11 | -6.648024 |
| 555 | CAPG | Edge | 0.23842418 | 5.50E-11 | 6.648024 |
| 556 | CCNE1 | Edge | 0.5789075 | 6.11E-11 | 6.631668 |
| 195 | CCNE1 | Core | -0.5789075 | 6.11E-11 | -6.631668 |
| 196 | CDK1 | Core | -0.4416989 | 1.18E-10 | -6.5329227 |
| 557 | CDK1 | Edge | 0.4416989 | 1.18E-10 | 6.5329227 |
| 558 | LHX6 | Edge | 1.1724633 | 2.06E-10 | 6.4483037 |
| 197 | LHX6 | Core | -1.1724633 | 2.06E-10 | -6.4483037 |
| 198 | FRZB | Core | -0.5092416 | 2.77E-10 | -6.402616 |
| 559 | FRZB | Edge | 0.5092416 | 2.77E-10 | 6.402616 |
| 560 | CCNB2 | Edge | 0.48973027 | 3.31E-10 | 6.374667 |
| 199 | CCNB2 | Core | -0.4897303 | 3.31E-10 | -6.374667 |
| 200 | ACTA2 | Core | 0.3699132 | 4.13E-10 | 6.3398123 |
| 561 | ACTA2 | Edge | -0.3699132 | 4.13E-10 | -6.3398123 |
| 562 | PDGFD | Edge | -0.3220327 | 4.24E-10 | -6.335155 |
| 201 | PDGFD | Core | 0.32203266 | 4.24E-10 | 6.335155 |
| 202 | STXBP2 | Core | -0.3128127 | 3.56E-09 | -5.9975023 |
| 563 | STXBP2 | Edge | 0.3128127 | 3.56E-09 | 5.9975023 |
| 564 | ANK1 | Edge | 0.8036745 | 5.04E-09 | 5.9400744 |
| 203 | ANK1 | Core | -0.8036745 | 5.04E-09 | -5.9400744 |
| 204 | CCL4 | Core | -1.0995011 | 1.10E-08 | -5.809337 |
| 565 | CCL4 | Edge | 1.0995011 | 1.10E-08 | 5.809337 |
| 566 | SNTB2 | Edge | 0.27509 | 1.12E-08 | 5.806532 |
| 205 | SNTB2 | Core | -0.27509 | 1.12E-08 | -5.806532 |
| 206 | CD48 | Core | -0.4410904 | 2.95E-08 | -5.641028 |
| 567 | CD48 | Edge | 0.44109043 | 2.95E-08 | 5.641028 |
| 568 | RYR2 | Edge | 1.2450243 | 3.17E-08 | 5.627429 |
| 207 | RYR2 | Core | -1.2450243 | 3.17E-08 | -5.627429 |
| 569 | FREM1 | Edge | 0.78304315 | 3.35E-08 | 5.61739 |
| 208 | FREM1 | Core | -0.7830432 | 3.35E-08 | -5.61739 |
| 209 | FSTL4 | Core | -1.3371712 | 3.86E-08 | -5.592034 |
| 570 | FSTL4 | Edge | 1.3371712 | 3.86E-08 | 5.592034 |
| 210 | CRYM | Core | -0.8210229 | 4.28E-08 | -5.5730352 |
| 571 | CRYM | Edge | 0.82102287 | 4.28E-08 | 5.5730352 |
| 572 | HS3ST4 | Edge | 1.4129283 | 4.95E-08 | 5.5468087 |
| 211 | HS3ST4 | Core | -1.4129283 | 4.95E-08 | -5.5468087 |
| 573 | FLT4 | Edge | 0.83333766 | 8.28E-08 | 5.45536 |
| 212 | FLT4 | Core | -0.8333377 | 8.28E-08 | -5.45536 |
| 213 | LRRK1 | Core | -0.7478058 | 1.98E-07 | -5.297441 |

| | | | | | |
|---|---|---|---|---|---|
| 574 | LRRK1 | Edge | 0.74780583 | 1.98E-07 | 5.297441 |
| 575 | SFRP2 | Edge | 0.37326762 | 2.20E-07 | 5.277207 |
| 214 | SFRP2 | Core | -0.3732676 | 2.20E-07 | -5.277207 |
| 215 | PTCHD4 | Core | -0.6594997 | 2.45E-07 | -5.2566776 |
| 576 | PTCHD4 | Edge | 0.6594997 | 2.45E-07 | 5.2566776 |
| 577 | MGST1 | Edge | -0.2345571 | 2.47E-07 | 5.254276 |
| 216 | MGST1 | Core | 0.2345571 | 2.47E-07 | -5.254276 |
| 217 | CDH6 | Core | 0.11958496 | 2.65E-07 | 5.2404585 |
| 578 | CDH6 | Edge | -0.119585 | 2.65E-07 | -5.2404585 |
| 218 | LXN | Core | -0.6109744 | 2.97E-07 | -5.218569 |
| 579 | LXN | Edge | 0.61097443 | 2.97E-07 | 5.218569 |
| 219 | TSHZ2 | Core | 0.36460632 | 2.97E-07 | 5.217595 |
| 580 | TSHZ2 | Edge | -0.3646063 | 2.97E-07 | -5.217595 |
| 220 | PLCE1 | Core | -0.2563904 | 4.46E-07 | -5.141015 |
| 581 | PLCE1 | Edge | 0.25639042 | 4.46E-07 | 5.141015 |
| 221 | ERBB2 | Core | -0.2566824 | 8.54E-07 | -5.016768 |
| 582 | ERBB2 | Edge | 0.25668237 | 8.54E-07 | 5.016768 |
| 583 | SNCG | Edge | 2.6551414 | 1.32E-06 | 4.9313455 |
| 222 | SNCG | Core | -2.6551414 | 1.32E-06 | -4.9313455 |
| 584 | GLI1 | Edge | 0.5172234 | 1.34E-06 | 4.9274635 |
| 223 | GLI1 | Core | -0.5172234 | 1.34E-06 | -4.9274635 |
| 224 | CCKBR | Core | -0.936617 | 1.53E-06 | -4.9014406 |
| 586 | RFTN1 | Edge | -0.2393168 | 1.53E-06 | -4.9006214 |
| 585 | CCKBR | Edge | 0.93661696 | 1.53E-06 | 4.9014406 |
| 225 | RFTN1 | Core | 0.23931676 | 1.53E-06 | 4.9006214 |
| 587 | ADAMTS3 | Edge | -0.4145953 | 1.71E-06 | -4.8773174 |
| 226 | ADAMTS3 | Core | 0.41459525 | 1.71E-06 | 4.8773174 |
| 227 | SLIT3 | Core | -0.8976896 | 2.13E-06 | -4.8331566 |
| 588 | SLIT3 | Edge | 0.89768964 | 2.13E-06 | 4.8331566 |
| 228 | CDH1 | Core | -0.6653818 | 3.27E-06 | -4.7458797 |
| 589 | CDH1 | Edge | 0.6653818 | 3.27E-06 | 4.7458797 |
| 229 | CTNNA3 | Core | -0.8648211 | 4.07E-06 | -4.7005386 |
| 590 | CTNNA3 | Edge | 0.86482114 | 4.07E-06 | 4.7005386 |
| 230 | CRHBP | Core | -1.0255715 | 4.51E-06 | -4.678556 |
| 591 | CRHBP | Edge | 1.0255715 | 4.51E-06 | 4.678556 |
| 592 | SDK1 | Edge | 0.44482732 | 6.52E-06 | 4.6018147 |
| 231 | SDK1 | Core | -0.4448273 | 6.52E-06 | -4.6018147 |
| 232 | GRAPL | Core | -0.6765979 | 1.03E-05 | -4.5047603 |
| 593 | GRAPL | Edge | 0.6765979 | 1.03E-05 | 4.5047603 |

| | | | | | |
|---|---|---|---|---|---|
| 594 | GRAP | Edge | 0.5517763 | 1.09E-05 | 4.492432 |
| 233 | GRAP | Core | -0.5517763 | 1.09E-05 | -4.492432 |
| 234 | LEF1 | Core | 0.08366995 | 1.21E-05 | 4.4681287 |
| 595 | LEF1 | Edge | -0.0836699 | 1.21E-05 | -4.4681287 |
| 596 | KLF2 | Edge | 0.40274355 | 1.90E-05 | 4.3702397 |
| 235 | KLF2 | Core | -0.4027436 | 1.90E-05 | -4.3702397 |
| 236 | PLD5 | Core | -0.1975315 | 1.95E-05 | -4.363112 |
| 597 | PLD5 | Edge | 0.19753145 | 1.95E-05 | 4.363112 |
| 598 | DCN | Edge | 0.7621513 | 2.04E-05 | 4.3527336 |
| 237 | DCN | Core | -0.7621513 | 2.04E-05 | -4.3527336 |
| 238 | MYC | Core | -0.4324891 | 2.17E-05 | -4.338046 |
| 599 | MYC | Edge | 0.4324891 | 2.17E-05 | 4.338046 |
| 600 | ANKRD18A | Edge | 0.5016557 | 4.28E-05 | 4.185688 |
| 239 | ANKRD18A | Core | -0.5016557 | 4.28E-05 | -4.185688 |
| 240 | SORCS1 | Core | -1.1319035 | 5.82E-05 | -4.1142936 |
| 601 | SORCS1 | Edge | 1.1319035 | 5.82E-05 | 4.1142936 |
| 602 | UNC5B | Edge | 0.2517519 | 6.45E-05 | 4.0896325 |
| 241 | UNC5B | Core | -0.2517519 | 6.45E-05 | -4.0896325 |
| 242 | APP | Core | -0.5607659 | 0.00012535 | -3.9316251 |
| 603 | APP | Edge | 0.5607659 | 0.00012535 | 3.9316251 |
| 604 | HIGD1B | Edge | 0.7638349 | 0.00013276 | 3.916806 |
| 243 | HIGD1B | Core | -0.7638349 | 0.00013276 | -3.916806 |
| 244 | TMEM132C | Core | -0.4883207 | 0.00013458 | -3.912529 |
| 605 | TMEM132C | Edge | 0.48832074 | 0.00013458 | 3.912529 |
| 606 | ANO3 | Edge | 0.36676556 | 0.00015491 | 3.8774328 |
| 245 | ANO3 | Core | -0.3667656 | 0.00015491 | -3.8774328 |
| 246 | NPY1R | Core | -0.499893 | 0.00018823 | -3.8287647 |
| 607 | NPY1R | Edge | 0.49989304 | 0.00018823 | 3.8287647 |
| 608 | CD47 | Edge | 0.14775482 | 0.0003058 | 3.7065768 |
| 247 | CD47 | Core | -0.1477548 | 0.0003058 | -3.7065768 |
| 248 | STAT3 | Core | -0.1246382 | 0.00033129 | -3.685216 |
| 609 | STAT3 | Edge | 0.12463817 | 0.00033129 | 3.685216 |
| 610 | SOX9 | Edge | 0.6555757 | 0.00041485 | 3.6264992 |
| 249 | SOX9 | Core | -0.6555757 | 0.00041485 | -3.6264992 |
| 250 | MGP | Core | 0.25999224 | 0.00066103 | 3.5032542 |
| 611 | MGP | Edge | -0.2599922 | 0.00066103 | -3.5032542 |
| 612 | NODAL | Edge | 0.82841176 | 0.00091718 | 3.4139647 |
| 251 | NODAL | Core | -0.8284118 | 0.00091718 | -3.4139647 |

G

**Supplemental Table 8. Survival of Differential Gene Expression**

| Gene | Region | Logfoldchange | p-value | Region | Logfoldchange | p-value | Survival p-value |
|---|---|---|---|---|---|---|---|
| POSTN | Core | 3.1 | 0 | Edge | -3 | 0 | 6.66E-05 |
| ADRA1A | Core | -0.9 | 1.13E-107 | Edge | 0.9 | 1.13E-107 | 0.00085 |
| TGFBI | Core | 1 | 2.73E-70 | Edge | -1 | 2.73E-70 | 0.000904 |
| CORO1A | Core | -1.2 | 7.33E-57 | Edge | 1.2 | 7.33E-57 | 0.000963 |
| ELOVL2 | Core | -0.2 | 4.18E-13 | Edge | -0.2 | 4.18E-13 | 0.001225 |
| SOX4 | Core | -2.8 | 0 | Edge | 2.8 | 0 | 0.003573 |
| GPNMB | Core | -1.2 | 3.65E-14 | Edge | 1.2 | 3.65E-14 | 0.00725 |
| IGFBP5 | Core | 8.1 | 0 | Edge | -8.1 | 0 | 0.008161 |
| LOX | Core | 3.6 | 9.96E-302 | Edge | -3.6 | 9.96E-302 | 0.01364 |

Supplemental Table 9

| channel | Expression | Coverage | Celltype-xenium | Metab Peak Name | core_id | gene-xenium | Region |
|---|---|---|---|---|---|---|---|
| **channel_168** | 0.13050485 | overlap | Mes-like | 162.0567 m/z Â± 10 ppm | GBM1-Core | LOX | Core |
| **channel_177** | 0.061371956 | overlap | Mes-like | Terephthalic acid | GBM10-Core | LOX | Edge |
| **channel_184** | 0.093231596 | overlap | Mes-like | Isovanillic acid | GBM10-Edge | LOX | Core |
| **channel_184** | 0.054492284 | overlap | Mes-like | Isovanillic acid | GBM2-Core | LOX | Core |
| **channel_184** | 0.0483286 | overlap | Mes-like | Isovanillic acid | GBM2-Edge | LOX | Core |
| **channel_184** | 0.026006455 | overlap | Mes-like | Isovanillic acid | GBM2-Edge | LOX | Core |
| **channel_198** | 0.12822813 | overlap | Mes-like | Hippuric acid | GBM2-Edge | LOX | Core |
| **channel_207** | 0.056506097 | overlap | Mes-like | 182.0482 m/z Â± 10 ppm | GBM3-Core | LOX | Core |
| **channel_227** | 0.5439773 | overlap | Mes-like | D-Glucuronic acid | GBM3-Core | LOX | Edge |
| **channel_250** | 0.026586771 | overlap | Mes-like | 200.1708 m/z Â± 10 ppm | GBM3-Core | LOX | Edge |
| **channel_281** | 0.027498333 | overlap | Mes-like | 213.0658 m/z Â± 10 ppm | GBM3-Edge | LOX | Core |
| **channel_316** | 0.060800873 | overlap | Mes-like | 228.0845 m/z Â± 10 ppm | GBM3-Edge | LOX | Core |
| **channel_316** | 0.04915527 | overlap | Mes-like | 228.0845 m/z Â± 10 ppm | GBM4-Core | LOX | Core |
| **channel_34** | 0.48366567 | overlap | Mes-like | 106.0671 m/z Â± 10 ppm | GBM4-Core | LOX | Edge |
| **channel_343** | 0.68520206 | overlap | Mes-like | 243.0752 m/z Â± 10 ppm | GBM4-Edge | LOX | Edge |
| **channel_346** | 0.03612054 | overlap | Mes-like | 244.0824 m/z Â± 10 ppm | GBM5-Core | LOX | Core |
| **channel_348** | 0.057909112 | overlap | Mes-like | N-Acetyl-DL-tryptophan | GBM5-Core | LOX | Edge |
| **channel_378** | 0.07554762 | overlap | Mes-like | 257.2365 m/z Â± 10 ppm | GBM5-Edge | LOX | Edge |
| **channel_389** | 0.028969109 | overlap | Mes-like | 269.1276 m/z Â± 10 ppm | GBM6-Core | LOX | Core |
| **channel_417** | 0.09551092 | overlap | Mes-like | Oleic acid | GBM6-Core | LOX | Edge |
| **channel_417** | 0.07303304 | overlap | Mes-like | Oleic acid | GBM6-Core | LOX | Edge |
| **channel_433** | 0.11059541 | overlap | Mes-like | 303.1251 m/z Â± 10 ppm | GBM6-Edge | LOX | Edge |
| **channel_442** | 0.027689682 | overlap | Mes-like | 311.1735 m/z Â± 10 ppm | GBM6-Edge | LOX | Core |
| **channel_50** | 0.024742188 | overlap | Mes-like | Creatinine | GBM6-Edge | LOX | Edge |
| **channel_532** | 0.049123686 | overlap | Mes-like | 87.0574 m/z Â± 10 ppm | GBM7-Core | LOX | Edge |
| **channel_537** | 0.06570686 | overlap | Mes-like | Glycerol | GBM8-Core | LOX | Core |
| **channel_575** | 0.038358748 | overlap | Mes-like | Phenylacetaldehyde | GBM8-Core | LOX | Edge |
| **channel_587** | 0.12755431 | overlap | Mes-like | Tyrosol | GBM8-Core | LOX | Core |
| **channel_591** | 0.08105323 | overlap | Mes-like | 1,3-Dimethyluracil | GBM8-Edge | LOX | Core |
| **channel_74** | 0.025681956 | overlap | Mes-like | 123.0815 m/z Â± 10 ppm | GBM8-Edge | LOX | Core |
| **channel_95** | 0.06546255 | overlap | Mes-like | 131.0615 m/z Â± 10 ppm | GBM9-Edge | LOX | Edge |

| channel | Expression | Coverage | Celltype-xenium | Metab Peak Name | core_id | gene-xenium | Region |
|---|---|---|---|---|---|---|---|
| **channel_159** | 0.025771044 | overlap | Mes-like | 154.0514 m/z Â± 10 ppm | GBM6-Core | IGFBP5 | Core |
| **channel_168** | 0.050224155 | overlap | Mes-like | 162.0567 m/z Â± 10 ppm | GBM8-Core | IGFBP5 | Core |
| **channel_184** | 0.014569905 | overlap | Mes-like | Isovanillic acid | GBM1-Core | IGFBP5 | Core |
| **channel_184** | 0.022439323 | overlap | Mes-like | Isovanillic acid | GBM2-Core | IGFBP5 | Core |
| **channel_184** | 0.09576897 | overlap | Mes-like | Isovanillic acid | GBM4-Core | IGFBP5 | Core |
| **channel_184** | 0.032371353 | overlap | Mes-like | Isovanillic acid | GBM7-Core | IGFBP5 | Core |
| **channel_187** | 0.02275742 | overlap | Mes-like | 169.0774 m/z Â± 10 ppm | GBM10-Edge | IGFBP5 | Edge |
| **channel_281** | 0.028617295 | overlap | Mes-like | 213.0658 m/z Â± 10 ppm | GBM6-Core | IGFBP5 | Core |
| **channel_366** | 0.043689422 | overlap | Mes-like | 253.0978 m/z Â± 10 ppm | GBM8-Core | IGFBP5 | Core |
| **channel_389** | 0.026226856 | overlap | Mes-like | 269.1276 m/z Â± 10 ppm | GBM6-Core | IGFBP5 | Core |
| **channel_396** | 0.042110726 | overlap | Mes-like | 271.1089 m/z Â± 10 ppm | GBM8-Core | IGFBP5 | Core |
| **channel_417** | 0.05450298 | overlap | Mes-like | Oleic acid | GBM1-Edge | IGFBP5 | Edge |
| **channel_442** | 0.02786883 | overlap | Mes-like | 311.1735 m/z Â± 10 ppm | GBM6-Core | IGFBP5 | Core |
| **channel_442** | 0.005521739 | overlap | Mes-like | 311.1735 m/z Â± 10 ppm | GBM5-Core | IGFBP5 | Core |
| **channel_537** | 0.06492386 | overlap | Mes-like | Glycerol | GBM4-Core | IGFBP5 | Core |
| **channel_554** | 0.23332223 | overlap | Mes-like | 100.0047 m/z Â± 10 ppm | GBM2-Edge | IGFBP5 | Edge |
| **channel_636** | 0.019963615 | overlap | Mes-like | Lipoamide | GBM10-Edge | IGFBP5 | Edge |
| **channel_665** | 0.02064679 | overlap | Mes-like | 296.1188 m/z Â± 10 ppm | GBM10-Edge | IGFBP5 | Edge |
| **channel_72** | 0.019713743 | overlap | Mes-like | 4-Methylcatechol | GBM8-Edge | IGFBP5 | Edge |

| channel | Expression | Coverage | Celltype-xenium | Metab Peak Name | core_id | gene-xenium | Region |
|---|---|---|---|---|---|---|---|
| **channel_100** | 0.035565615 | overlap | NPC-like | Homocysteine | GBM8-Core | SOX4 | Core |
| **channel_155** | 0.04899725 | overlap | NPC-like | Dopamine | GBM6-Core | SOX4 | Core |
| **channel_157** | 0.03742808 | overlap | NPC-like | 2,3-Dihydroxbenzoic acid | GBM6-Core | SOX4 | Core |
| **channel_168** | 0.04145586 | overlap | NPC-like | 162.0567 m/z Â± 10 ppm | GBM8-Core | SOX4 | Core |
| **channel_173** | 0.036950856 | overlap | NPC-like | 164.0558 m/z Â± 10 ppm | GBM6-Core | SOX4 | Core |
| **channel_184** | 0.08235717 | overlap | NPC-like | Isovanillic acid | GBM4-Core | SOX4 | Core |
| **channel_184** | 0.025795674 | overlap | NPC-like | Isovanillic acid | GBM1-Core | SOX4 | Core |
| **channel_198** | 0.036074474 | overlap | NPC-like | Hippuric acid | GBM8-Core | SOX4 | Core |
| **channel_396** | 0.03749559 | overlap | NPC-like | 271.2283 m/z Â± 10 ppm | GBM8-Core | SOX4 | Core |
| **channel_417** | 0.060738802 | overlap | NPC-like | Oleic acid | GBM1-Edge | SOX4 | Edge |
| **channel_442** | 0.029613141 | overlap | NPC-like | 311.1735 m/z Â± 10 ppm | GBM2-Core | SOX4 | Core |
| **channel_537** | 0.06671638 | overlap | NPC-like | Glycerol | GBM4-Core | SOX4 | Core |
| **channel_545** | 0.03282926 | overlap | NPC-like | 96.0489 m/z Â± 10 ppm | GBM10-Edge | SOX4 | Edge |
| **channel_556** | 0.25758934 | overlap | NPC-like | Valeric acid | GBM2-Edge | SOX4 | Edge |
| **channel_575** | 0.031850662 | overlap | NPC-like | Phenylacetaldehyde | GBM10-Edge | SOX4 | Edge |
| **channel_586** | 0.017074626 | overlap | NPC-like | 133.0662 m/z Â± 10 ppm | GBM5-Core | SOX4 | Core |
| **channel_631** | 0.01994482 | overlap | NPC-like | 3-Hydroxymethylglutaric acid | GBM5-Core | SOX4 | Core |
| **channel_9** | 0.038088154 | overlap | NPC-like | 92.0519 m/z Â± 10 ppm | GBM8-Core | SOX4 | Core |

| channel | Expression | Coverage | Celltype-xenium | Metab Peak Name | core_id | gene-xenium | Region |
|---|---|---|---|---|---|---|---|
| **channel_109** | 0.05942491 | overlap | AC-like | Tyramine | GBM6-Core | ELOVL2 | Core |
| **channel_118** | 0.041087344 | overlap | AC-like | O-Phosphoethanolamine | GBM10-Edge | ELOVL2 | Edge |
| **channel_172** | 0.043898083 | overlap | AC-like | Lumazine | GBM6-Core | ELOVL2 | Core |
| **channel_173** | 0.17603827 | overlap | AC-like | 164.0558 m/z Â± 10 ppm | GBM8-Core | ELOVL2 | Core |
| **channel_184** | 0.022192262 | overlap | AC-like | Isovanillic acid | GBM1-Core | ELOVL2 | Core |
| **channel_184** | 0.08533944 | overlap | AC-like | Isovanillic acid | GBM4-Core | ELOVL2 | Core |
| **channel_184** | 0.034343902 | overlap | AC-like | Isovanillic acid | GBM7-Core | ELOVL2 | Core |
| **channel_187** | 0.039270815 | overlap | AC-like | 169.0774 m/z Â± 10 ppm | GBM10-Edge | ELOVL2 | Edge |
| **channel_235** | 0.024733523 | overlap | AC-like | Aminoadipic acid | GBM9-Edge | ELOVL2 | Edge |
| **channel_266** | 0.04579407 | overlap | AC-like | L-Kynurenine | GBM8-Edge | ELOVL2 | Edge |
| **channel_382** | 0.17558357 | overlap | AC-like | 262.096 m/z Â± 10 ppm | GBM8-Core | ELOVL2 | Core |
| **channel_395** | 0.17253087 | overlap | AC-like | 271.0222 m/z Â± 10 ppm | GBM8-Core | ELOVL2 | Core |
| **channel_417** | 0.061862137 | overlap | AC-like | Oleic acid | GBM1-Edge | ELOVL2 | Edge |
| **channel_442** | 0.046765637 | overlap | AC-like | 311.1735 m/z Â± 10 ppm | GBM6-Core | ELOVL2 | Core |
| **channel_524** | 0.27254343 | overlap | AC-like | 81.9539 m/z Â± 10 ppm | GBM8-Core | ELOVL2 | Core |
| **channel_531** | 0.038300127 | overlap | AC-like | Acetoin | GBM10-Edge | ELOVL2 | Edge |
| **channel_537** | 0.07502741 | overlap | AC-like | Glycerol | GBM4-Core | ELOVL2 | Core |
| **channel_540** | 0.0484335 | overlap | AC-like | 92.9291 m/z Â± 10 ppm | GBM8-Edge | ELOVL2 | Edge |
| **channel_588** | 0.07573779 | overlap | AC-like | 137.0981 m/z Â± 10 ppm | GBM4-Core | ELOVL2 | Core |
| **channel_603** | 0.034376908 | overlap | AC-like | 153.9991 m/z Â± 10 ppm | GBM10-Edge | ELOVL2 | Edge |
| **channel_622** | 0.050466094 | overlap | AC-like | Methylglutaric acid | GBM6-Core | ELOVL2 | Core |
| **channel_68** | 0.025340898 | overlap | AC-like | Nicotinic acid | GBM9-Edge | ELOVL2 | Edge |

| channel | Expression | Coverage | Celltype-xenium | Metab Peak Name | core_id | gene-xenium | Region |
|---|---|---|---|---|---|---|---|
| **channel_118** | 0.08656756 | overlap | Mes-like | O-Phosphoethanolamine | GBM4-Edge | TGFBI | Edge |
| **channel_159** | 0.04307685 | overlap | Mes-like | 154.0514 m/z Â± 10 ppm | GBM3-Core | TGFBI | Core |
| **channel_159** | 0.027094899 | overlap | Mes-like | 154.0514 m/z Â± 10 ppm | GBM6-Core | TGFBI | Core |
| **channel_168** | 0.18133661 | overlap | Mes-like | 162.0567 m/z Â± 10 ppm | GBM8-Core | TGFBI | Core |
| **channel_168** | 0.029011989 | overlap | Mes-like | 162.0567 m/z Â± 10 ppm | GBM6-Core | TGFBI | Core |
| **channel_184** | 0.023074498 | overlap | Mes-like | Isovanillic acid | GBM2-Core | TGFBI | Core |
| **channel_184** | 0.091883786 | overlap | Mes-like | Isovanillic acid | GBM4-Core | TGFBI | Core |
| **channel_184** | 0.040753428 | overlap | Mes-like | Isovanillic acid | GBM7-Core | TGFBI | Core |
| **channel_184** | 0.026445951 | overlap | Mes-like | Isovanillic acid | GBM10-Core | TGFBI | Core |
| **channel_187** | 0.017706556 | overlap | Mes-like | 169.0774 m/z Â± 10 ppm | GBM9-Edge | TGFBI | Edge |
| **channel_194** | 0.55867976 | overlap | Mes-like | Pyrophosphoric Acid | GBM2-Edge | TGFBI | Edge |
| **channel_227** | 0.71426934 | overlap | Mes-like | D-Glucuronic acid | GBM2-Edge | TGFBI | Edge |
| **channel_238** | 0.07535851 | overlap | Mes-like | 196.1346 m/z Â± 10 ppm | GBM10-Edge | TGFBI | Edge |
| **channel_26** | 0.057564665 | overlap | Mes-like | 100.0422 m/z Â± 10 ppm | GBM6-Edge | TGFBI | Edge |
| **channel_281** | 0.03325049 | overlap | Mes-like | 213.0658 m/z Â± 10 ppm | GBM6-Core | TGFBI | Core |
| **channel_287** | 0.058134116 | overlap | Mes-like | 215.0458 m/z Â± 10 ppm | GBM5-Core | TGFBI | Core |
| **channel_343** | 0.71545666 | overlap | Mes-like | 243.0752 m/z Â± 10 ppm | GBM2-Edge | TGFBI | Edge |
| **channel_389** | 0.026219582 | overlap | Mes-like | 269.1276 m/z Â± 10 ppm | GBM1-Core | TGFBI | Core |
| **channel_417** | 0.07284638 | overlap | Mes-like | Oleic acid | GBM1-Edge | TGFBI | Edge |
| **channel_442** | 0.04177016 | overlap | Mes-like | 311.1735 m/z Â± 10 ppm | GBM9-Core | TGFBI | Core |
| **channel_452** | 0.22031853 | overlap | Mes-like | 324.1011 m/z Â± 10 ppm | GBM8-Core | TGFBI | Core |
| **channel_453** | 0.092887245 | overlap | Mes-like | 325.109 m/z Â± 10 ppm | GBM4-Edge | TGFBI | Edge |
| **channel_512** | 0.30089906 | overlap | Mes-like | 399.2649 m/z Â± 10 ppm | GBM8-Core | TGFBI | Core |
| **channel_586** | 0.1703939 | overlap | Mes-like | 133.0662 m/z Â± 10 ppm | GBM8-Core | TGFBI | Core |
| **channel_591** | 0.062055413 | overlap | Mes-like | 1,3-Dimethyluracil | GBM5-Core | TGFBI | Core |
| **channel_601** | 0.09039967 | overlap | Mes-like | Perillyl alcohol | GBM4-Edge | TGFBI | Edge |
| **channel_64** | 0.0664031 | overlap | Mes-like | Benzamide | GBM8-Edge | TGFBI | Edge |
| **channel_665** | 0.07813882 | overlap | Mes-like | 296.1188 m/z Â± 10 ppm | GBM5-Edge | TGFBI | Edge |
| **channel_88** | 0.17848629 | overlap | Mes-like | 128.052 m/z Â± 10 ppm | GBM8-Core | TGFBI | Core |

| channel | Expression | Coverage | Celltype-xenium | Metab Peak Name | core_id | gene-xenium | Region |
|---|---|---|---|---|---|---|---|
| **channel_481** | 1.0505718 | overlap | AC-like | 347.1477 m/z Â± 10 ppm | GBM2-Edge | POSTN | Edge |
| **channel_394** | 0.88394094 | overlap | AC-like | 270.2482 m/z Â± 10 ppm | GBM2-Edge | POSTN | Edge |
| **channel_234** | 0.8075721 | overlap | AC-like | 195.1388 m/z Â± 10 ppm | GBM2-Edge | POSTN | Edge |
| **channel_591** | 0.1868783 | overlap | AC-like | 1,3-Dimethyluracil | GBM8-Core | POSTN | Core |
| **channel_406** | 0.16891211 | overlap | AC-like | 277.181 m/z Â± 10 ppm | GBM8-Core | POSTN | Core |
| **channel_62** | 0.14826262 | overlap | AC-like | Erythrose | GBM4-Core | POSTN | Core |
| **channel_395** | 0.10483249 | overlap | AC-like | 271.0222 m/z Â± 10 ppm | GBM3-Core | POSTN | Core |
| **channel_163** | 0.1037368 | overlap | AC-like | Indole-3-carboxylic acid | GBM4-Edge | POSTN | Edge |
| **channel_440** | 0.09935482 | overlap | AC-like | 310.1188 m/z Â± 10 ppm | GBM4-Edge | POSTN | Edge |
| **channel_106** | 0.09911834 | overlap | AC-like | Adenine | GBM4-Edge | POSTN | Edge |
| **channel_50** | 0.09476058 | overlap | AC-like | Creatinine | GBM4-Edge | POSTN | Edge |
| **channel_48** | 0.08370923 | overlap | AC-like | 111.9572 m/z Â± 10 ppm | GBM1-Edge | POSTN | Edge |
| **channel_187** | 0.08337231 | overlap | AC-like | 169.0774 m/z Â± 10 ppm | GBM10-Edge | POSTN | Edge |
| **channel_538** | 0.07972165 | overlap | AC-like | 92.0252 m/z Â± 10 ppm | GBM3-Core | POSTN | Core |
| **channel_197** | 0.07364221 | overlap | AC-like | 5-Phenylvaleric acid | GBM3-Core | POSTN | Core |
| **channel_232** | 0.0675795 | overlap | AC-like | Alpha-hydroxyhippuric acid | GBM9-Edge | POSTN | Edge |
| **channel_302** | 0.06737968 | overlap | AC-like | N,N,N-Trimethyllysine | GBM9-Edge | POSTN | Edge |
| **channel_422** | 0.05903401 | overlap | AC-like | 285.1159 m/z Â± 10 ppm | GBM8-Edge | POSTN | Edge |
| **channel_522** | 0.05565382 | overlap | AC-like | 81.0351 m/z Â± 10 ppm | GBM8-Edge | POSTN | Edge |
| **channel_440** | 0.051983852 | overlap | AC-like | 310.1188 m/z Â± 10 ppm | GBM8-Edge | POSTN | Edge |
| **channel_184** | 0.05016152 | overlap | AC-like | Isovanillic acid | GBM7-Core | POSTN | Core |
| **channel_326** | 0.04727498 | overlap | AC-like | 235.0365 m/z Â± 10 ppm | GBM3-Edge | POSTN | Edge |
| **channel_30** | 0.04687174 | overlap | AC-like | 102.9488 m/z Â± 10 ppm | GBM8-Edge | POSTN | Edge |
| **channel_417** | 0.04519724 | overlap | AC-like | Oleic acid | GBM6-Edge | POSTN | Edge |
| **channel_639** | 0.043717388 | overlap | AC-like | L-Arginine | GBM3-Edge | POSTN | Edge |
| **channel_50** | 0.04067854 | overlap | AC-like | Creatinine | GBM7-Edge | POSTN | Edge |
| **channel_432** | 0.037840463 | overlap | AC-like | 299.0514 m/z Â± 10 ppm | GBM3-Edge | POSTN | Edge |
| **channel_429** | 0.034170438 | overlap | AC-like | 295.1092 m/z Â± 10 ppm | GBM7-Edge | POSTN | Edge |
| **channel_281** | 0.03240716 | overlap | AC-like | 213.0658 m/z Â± 10 ppm | GBM6-Core | POSTN | Core |
| **channel_159** | 0.029950732 | overlap | AC-like | 154.0514 m/z Â± 10 ppm | GBM6-Core | POSTN | Core |
| **channel_611** | 0.02755815 | overlap | AC-like | Leucinic acid | GBM6-Core | POSTN | Core |
| **channel_184** | 0.02726676 | overlap | AC-like | Isovanillic acid | GBM1-Core | POSTN | Core |
| **channel_389** | 0.027051091 | overlap | AC-like | 269.1276 m/z Â± 10 ppm | GBM6-Core | POSTN | Core |
| **channel_442** | 0.022419294 | overlap | AC-like | 311.1735 m/z Â± 10 ppm | GBM10-Core | POSTN | Core |
| **channel_442** | 0.005785395 | overlap | AC-like | Guanine | GBM5-Core | POSTN | Core |

| channel | Expression | Coverage | Celltype-xenium | Metab Peak Name | core_id | gene-xenium | Region |
|---|---|---|---|---|---|---|---|
| **channel_115** | 0.050113566 | overlap | Mes-like | Phosphonoacetate | GBM8-Edge | GPNMB | Edge |
| **channel_126** | 0.04563749 | overlap | Mes-like | 142.0511 m/z Â± 10 ppm | GBM4-Edge | GPNMB | Edge |
| **channel_15** | 0.030231813 | overlap | Mes-like | 95.0509 m/z Â± 10 ppm | GBM6-Core | GPNMB | Core |
| **channel_175** | 0.041298907 | overlap | Mes-like | Terephthalic acid | GBM5-Core | GPNMB | Core |
| **channel_184** | 0.06507892 | overlap | Mes-like | Isovanillic acid | GBM1-Core | GPNMB | Core |
| **channel_184** | 0.093612194 | overlap | Mes-like | Isovanillic acid | GBM4-Core | GPNMB | Core |
| **channel_184** | 0.043530107 | overlap | Mes-like | Isovanillic acid | GBM7-Core | GPNMB | Core |
| **channel_223** | 0.031927846 | overlap | Mes-like | 2-Keto-D-gluconic Acid | GBM6-Core | GPNMB | Core |
| **channel_231** | 0.0334946 | overlap | Mes-like | 112.0407 m/z Â± 10 ppm | GBM3-Edge | GPNMB | Edge |
| **channel_235** | 0.020502541 | overlap | Mes-like | Oxoadipate | GBM9-Edge | GPNMB | Edge |
| **channel_24** | 0.04682198 | overlap | Mes-like | 99.0101 m/z Â± 10 ppm | GBM8-Edge | GPNMB | Edge |
| **channel_264** | 0.077113226 | overlap | Mes-like | 208.037 m/z Â± 10 ppm | GBM6-Edge | GPNMB | Edge |
| **channel_35** | 0.42297605 | overlap | Mes-like | 106.9811 m/z Â± 10 ppm | GBM2-Edge | GPNMB | Edge |
| **channel_382** | 0.036543734 | overlap | Mes-like | 262.096 m/z Â± 10 ppm | GBM9-Core | GPNMB | Core |
| **channel_396** | 0.18195796 | overlap | Mes-like | 271.1089 m/z Â± 10 ppm | GBM8-Core | GPNMB | Core |
| **channel_40** | 0.038921636 | overlap | Mes-like | 4-Aminophenol | GBM10-Core | GPNMB | Core |
| **channel_430** | 0.072743766 | overlap | Mes-like | 13-HODE | GBM6-Edge | GPNMB | Edge |
| **channel_432** | 0.031454742 | overlap | Mes-like | 299.0514 m/z Â± 10 ppm | GBM3-Edge | GPNMB | Edge |
| **channel_442** | 0.0280965 | overlap | Mes-like | 311.1735 m/z Â± 10 ppm | GBM6-Core | GPNMB | Core |
| **channel_451** | 0.38008234 | overlap | Mes-like | 323.0947 m/z Â± 10 ppm | GBM2-Edge | GPNMB | Edge |
| **channel_452** | 0.03672933 | overlap | Mes-like | 324.1011 m/z Â± 10 ppm | GBM5-Core | GPNMB | Core |
| **channel_458** | 0.030360922 | overlap | Mes-like | 326.1429 m/z Â± 10 ppm | GBM10-Core | GPNMB | Core |
| **channel_52** | 0.019674784 | overlap | Mes-like | 113.0248 m/z Â± 10 ppm | GBM9-Edge | GPNMB | Edge |
| **channel_537** | 0.09172344 | overlap | Mes-like | Glycerol | GBM4-Core | GPNMB | Core |
| **channel_540** | 0.036899395 | overlap | Mes-like | 92.9291 m/z Â± 10 ppm | GBM4-Edge | GPNMB | Edge |
| **channel_541** | 0.04683135 | overlap | Mes-like | 93.9744 m/z Â± 10 ppm | GBM5-Core | GPNMB | Core |
| **channel_543** | 0.036312245 | overlap | Mes-like | 95.0257 m/z Â± 10 ppm | GBM4-Edge | GPNMB | Edge |
| **channel_545** | 0.02934975 | overlap | Mes-like | 96.0489 m/z Â± 10 ppm | GBM10-Edge | GPNMB | Edge |
| **channel_563** | 0.029757567 | overlap | Mes-like | 109.0408 m/z Â± 10 ppm | GBM6-Core | GPNMB | Core |
| **channel_586** | 0.035994876 | overlap | Mes-like | 133.0662 m/z Â± 10 ppm | GBM9-Core | GPNMB | Core |
| **channel_6** | 0.08003692 | overlap | Mes-like | 85.0307 m/z Â± 10 ppm | GBM4-Core | GPNMB | Core |
| **channel_602** | 0.037108332 | overlap | Mes-like | 4-Hydroxy-3-methylbenzoic acid | GBM4-Edge | GPNMB | Edge |
| **channel_602** | 0.038460597 | overlap | Mes-like | 4-Hydroxy-3-methylbenzoic acid | GBM10-Edge | GPNMB | Edge |
| **channel_603** | 0.108643614 | overlap | Mes-like | 153.9991 m/z Â± 10 ppm | GBM1-Edge | GPNMB | Edge |
| **channel_622** | 0.034099355 | overlap | Mes-like | Methylglutaric acid | GBM10-Core | GPNMB | Core |
| **channel_645** | 0.057279337 | overlap | Mes-like | 217.0344 m/z Â± 10 ppm | GBM5-Edge | GPNMB | Edge |
| **channel_649** | 0.030605122 | overlap | Mes-like | 227.1172 m/z Â± 10 ppm | GBM10-Core | GPNMB | Core |
| **channel_77** | 0.03153541 | overlap | Mes-like | 5-Methylcytosine | GBM10-Edge | GPNMB | Edge |
| **channel_84** | 0.1435745 | overlap | Mes-like | 126.9955 m/z Â± 10 ppm | GBM8-Core | GPNMB | Core |
| **channel_89** | 0.1668127 | overlap | Mes-like | Itaconic acid | GBM8-Core | GPNMB | Core |
